# *Plasmodium falciparum* and helminth coinfections increase IgE and parasite-specific IgG responses

**DOI:** 10.1101/2021.05.26.445753

**Authors:** Rebeca Santano, Rocío Rubio, Berta Grau-Pujol, Valdemiro Escola, Osvaldo Muchisse, Inocência Cuamba, Marta Vidal, Pau Cisteró, Gemma Ruiz-Olalla, Ruth Aguilar, Maria Demontis, Jose Carlos Jamine, Anélsio Cossa, Charfudin Sacoor, Jorge Cano, Luis Izquierdo, Chetan E Chitnis, Ross L Coppel, Virander Chauhan, David Cavanagh, Sheetij Dutta, Evelina Angov, Deepak Gaur, Lisette van Lieshout, Bin Zhan, José Muñoz, Gemma Moncunill, Carlota Dobaño

**Affiliations:** ISGlobal, Hospital Clínic - Universitat de Barcelona, Barcelona, Spain; Centro de Investigação em Saúde de Manhiça (CISM), Maputo, Mozambique; Fundación Mundo Sano, Argentina; Department of Parasitology, Centre of Infectious Diseases, Leiden University Medical Centre (LUMC), Leiden, The Netherlands; Communicable and Non-communicable Diseases Cluster (UCN), WHO Regional Office for Africa, Brazzaville, Republic of Congo; Malaria Parasite Biology and Vaccines, Department of Parasites & Insect Vectors, Institut Pasteur, Paris, France; Department of Microbiology, Faculty of Medicine, Nursing and Health Sciences Monash University, Melbourne, Australia; Malaria Group, International Centre for Genetic Engineering and Biotechnology (ICGEB), New Delhi, India; Institute of Immunology and Infection Research, University of Edinburgh, Edinburgh, UK; Walter Reed Army Institute of Research (WRAIR), Maryland, USA; Laboratory of Malaria & Vaccine Research, School of Biotechnology, Jawaharlal Nehru University, New Delhi, India; Baylor College of Medicine (BCM), Texas, USA

## Abstract

Coinfection with *Plasmodium falciparum* and helminths may impact the immune response to these parasites since they induce different immune profiles. We studied the effects of coinfections on the antibody profile in a cohort of 715 Mozambican children and adults using the Luminex technology with a panel of 16 antigens from *P. falciparum* and 11 antigens from helminths (*Ascaris lumbricoides*, hookworm, *Trichuris trichiura*, *Strongyloides stercoralis* and *Schistosoma* spp.) and measured antigen-specific IgG and total IgE responses. We compared the antibody profile between groups defined by *P. falciparum* and helminth previous exposure (based on serology) and/or current infection (determined by microscopy and/or qPCR). In multivariable regression models adjusted by demographic, socioeconomic, water and sanitation variables, individuals exposed/infected with *P. falciparum* and helminths had significantly higher total IgE and antigen-specific IgG levels, magnitude (sum of all levels) and breadth of response to both types of parasites compared to individuals exposed/infected with only one type of parasite (p≤ 0.05). There was a positive association between exposure/infection to *P. falciparum* and exposure/infection to helminths or the number of helminth species, and *vice versa* (p≤ 0.001). In addition, children coexposed/coinfected tended (p= 0.062) to have higher *P. falciparum* parasitemia than those single exposed/infected. Our results suggest that an increase in the antibody responses in coexposed/coinfected individuals may reflect higher exposure and be due to a more permissive immune environment to infection in the host.

**GRAPHICAL ABSTRACT:** 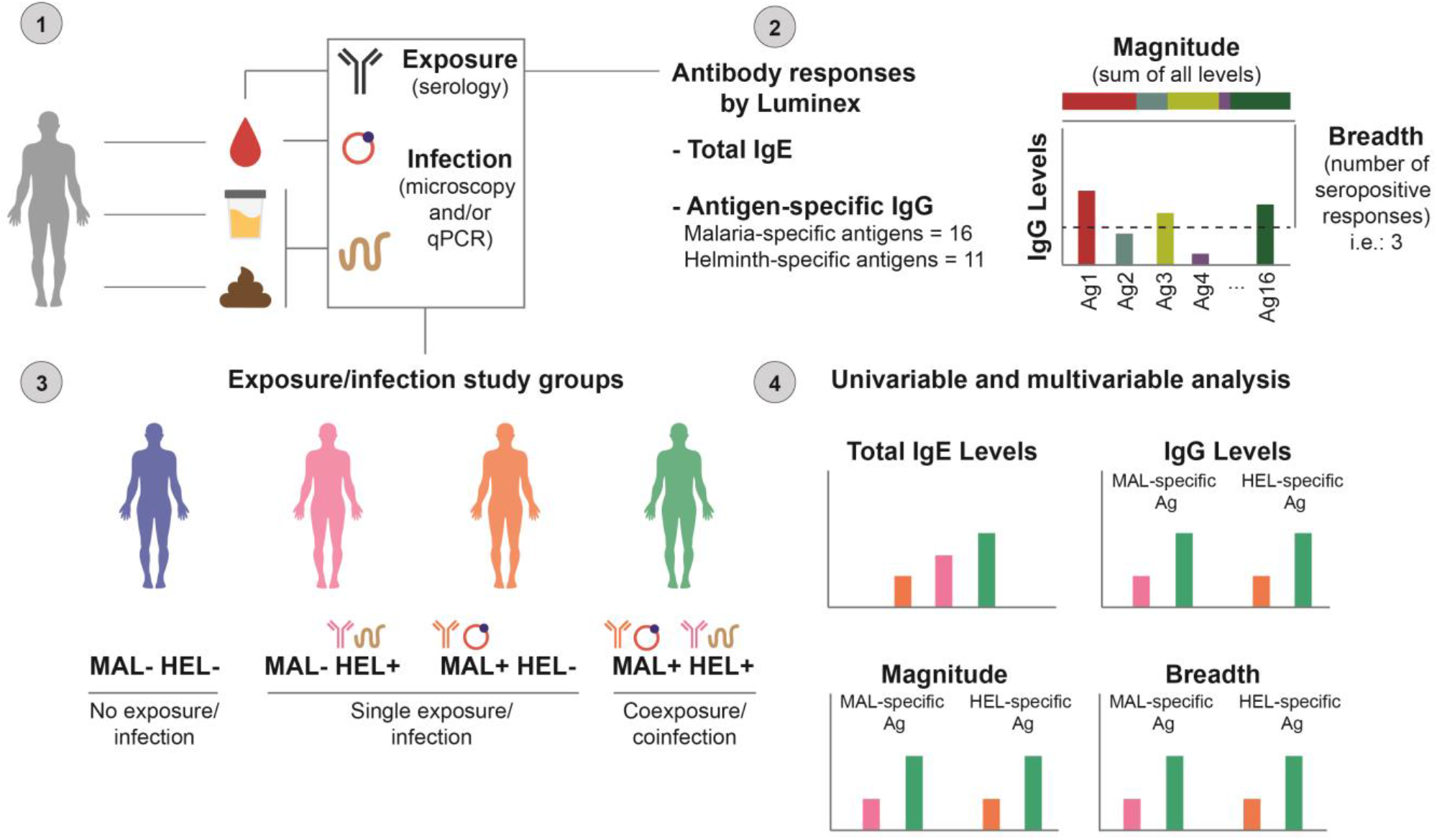

## INTRODUCTION

Malaria and helminthiasis are endemic parasitic diseases in tropical and subtropical areas, especially in impoverished countries with poor water and sanitation access. Their overlapping spatial distribution makes coinfections with these pathogens a frequent event (Brooker et al., 2006). Both cause high burden of morbidity, and malaria is a leading cause of mortality, particularly in children (World Health Organization, 2020a, 2020b, 2020c). The highest burden of malaria cases (82%) and malaria deaths (94%) occurs in sub-Saharan Africa (World Health Organization, 2020a), where *Plasmodium falciparum* is the most prevalent species causing malaria (World Health Organization, 2021). Helminths such as *Schistosoma* spp. and soil-transmitted helminths (STH) (*Ascaris lumbricoides, Trichuris trichiura, Ancylostoma duodenale, Necator americanus and Strongyloides stercoralis*) are also prevalent in Sub-Saharan Africa affecting more than 1.5 billion people worldwide (World Health Organization, 2020b, 2020c).

*P. falciparum* and helminths induce different types of immune responses. On one hand, the clearance of *P. falciparum* infection is achieved by an initial T helper (Th)1 response with the production of IgG antibodies, mainly from the cytophilic IgG1 and IgG3 subclasses (Bouharoun-Tayoun and Druilhe, 1992; Cohen et al., 1961). Evidence from animal models and humans suggest that later, a Th2/T regulatory (Treg) response is required to counteract severe immunopathological consequences of an excessive proinflammatory Th1 milieu (Hartgers and Yazdanbakhsh, 2006; Langhorne et al., 1998; Malaguarnera and Musumeci, 2002). On the other hand, helminths generally induce a Th2 polarization with the production of IgE and IgG4 antibodies (Hartgers and Yazdanbakhsh, 2006; Maizels et al., 2004; Maizels and Yazdanbakhsh, 2003). However, helminths are a heterogeneous group of parasites, each with a complex life cycle; therefore, differences exist by species and life-cycle stage. For example, *Schistosoma* spp. infection induces an acute proinflammatory Th1 response while there is a shift towards a Th2 response upon deposition of eggs (Pearce and MacDonald, 2002). Adding another layer of complexity, helminths are well known for their immunomodulating effects that deviate the cellular response towards a regulatory profile. This allows helminths to survive in the host for years and this may affect not only helminth antigens but also bystander antigens (Hartgers and Yazdanbakhsh, 2006; Maizels et al., 2004; Maizels and Yazdanbakhsh, 2003; McSorley and Maizels, 2012).

By altering the immunological balance, coinfections can impact immunity and course of infections. A paradox exists for *P. falciparum*: a Th2 and Treg response could protect against severe disease, but it may render subjects more susceptible to *P. falciparum* infection. This may explain the conflicting results existing in the literature (Degarege and Erko, 2016). In some cases, helminths have been associated with detrimental effects on *P. falciparum* infection, complications and parasite load (Babamale et al., 2018; Degarege et al., 2012, 2009; Le Hesran et al., 2004; Nacher et al., 2002b, 2001a; Ntonifor et al., 2021; Sokhna et al., 2004; Spiegel et al., 2003; Tshikuka et al., 1996) with protective effects in other cases (Briand et al., 2005; Brutus et al., 2007, 2006; Efunshile et al., 2015; Lemaitre et al., 2014; Lyke et al., 2005; Murray et al., 1978; Nacher et al., 2002a, 2001b, 2000) or no effect (Shapiro et al., 2005). The outcome may be critically influenced by the species, timing (Salazar-Castañón et al., 2018), parasite load, (Briand et al., 2005; Lemaitre et al., 2014; Lyke et al., 2005), and endpoint analyzed (infection or disease) (Degarege et al., 2016). Additionally, the limited information available regarding the effect of *P. falciparum* on helminth infections in humans suggests that *P. falciparum* could also affect the frequency and course of helminth infections, perhaps by delaying or dampening the production of required Th2 cytokines. In fact, a study showed that coinfection with malaria reduced the Th2 cytokine IL-4 in response to hookworm antigens (Quinnell et al., 2004). Lessons learned from other pathogens that induce Th1 responses support these findings, where the human T cell lymphotropic virus type 1 (HTLV-1) in coinfection with *S. stercoralis* also reduced the Th2 cytokine IL-5 and IgE (Porto et al., 2001), and previous leishmaniasis increased the probability to subsequently be infected with *S. mansoni* (Miranda et al., 2021). This is relevant since a Th1 polarization, depending on the species and life-stage, has been associated with persistence of helminth infection (Artis and Grencis, 2008).

Based on the different immune responses triggered by *P. falciparum* and helminths and the immunomodulating effect of helminths, we hypothesized that prior exposure to *P. falciparum* and helminths or their coinfection, would have an impact on the immune response to these parasites. Thus, our objective was to describe the effects of *P. falciparum* and helminth coinfections on the antibody profiles in an endemic cohort from Southern Mozambique. We measured antigen-specific IgG and total IgE responses with a multiplex suspension array technology (Luminex) including a panel of 16 antigens from different life stages of *P. falciparum* and 11 antigens from helminths (STH and *Schistosoma* spp.). The antibody profile was compared between different exposure/infection groups defined by previous exposure (based on serology) and/or the presence of current infection (determined by microscopy and/or qPCR).

## RESULTS

### Characteristics of the study population

The study included 715 subjects (363 children and 352 adults) from six different locations of Manhiça district, in the Maputo province, Southern Mozambique (**Table 1**). From all the participants, 9.93% were asymptomatically infected with *P. falciparum* as diagnosed by qPCR, and 53.57% carried a helminth infection (diagnosed by either qPCR and/or microscopy) (**Table S1**). Twenty-eight percent were infected by hookworm (*A. duodenale* and/or *N. americanus*), 14.41% by *T. trichiura*, 12.03% by *Schistosoma* spp., 10.35% by *S. stercoralis* and 7.41% by *A. lumbricoides.* From those infected with helminths, 29.24% were infected with more than one helminth species, with the coinfection of hookworm with *S. stercoralis* being the most common (**Table S2** and **Figure S1A**). Assessed by serology, 91.05% of individuals had been exposed to *P. falciparum* and 79.16% to any helminth (69.51% to hookworm, 52.73% to *T. trichiura,* 43.92% to *Schistosoma* spp. 27.27% to *S. stercoralis* and 20.70% to *A. lumbricoides*) (**Table S1**). Most of the helminth exposed individuals (77.21%) had also been exposed to multiple helminth species, mainly to hookworm and *T. trichiura* (**Table S2** and **Figure S1B**).

**Table 1.**
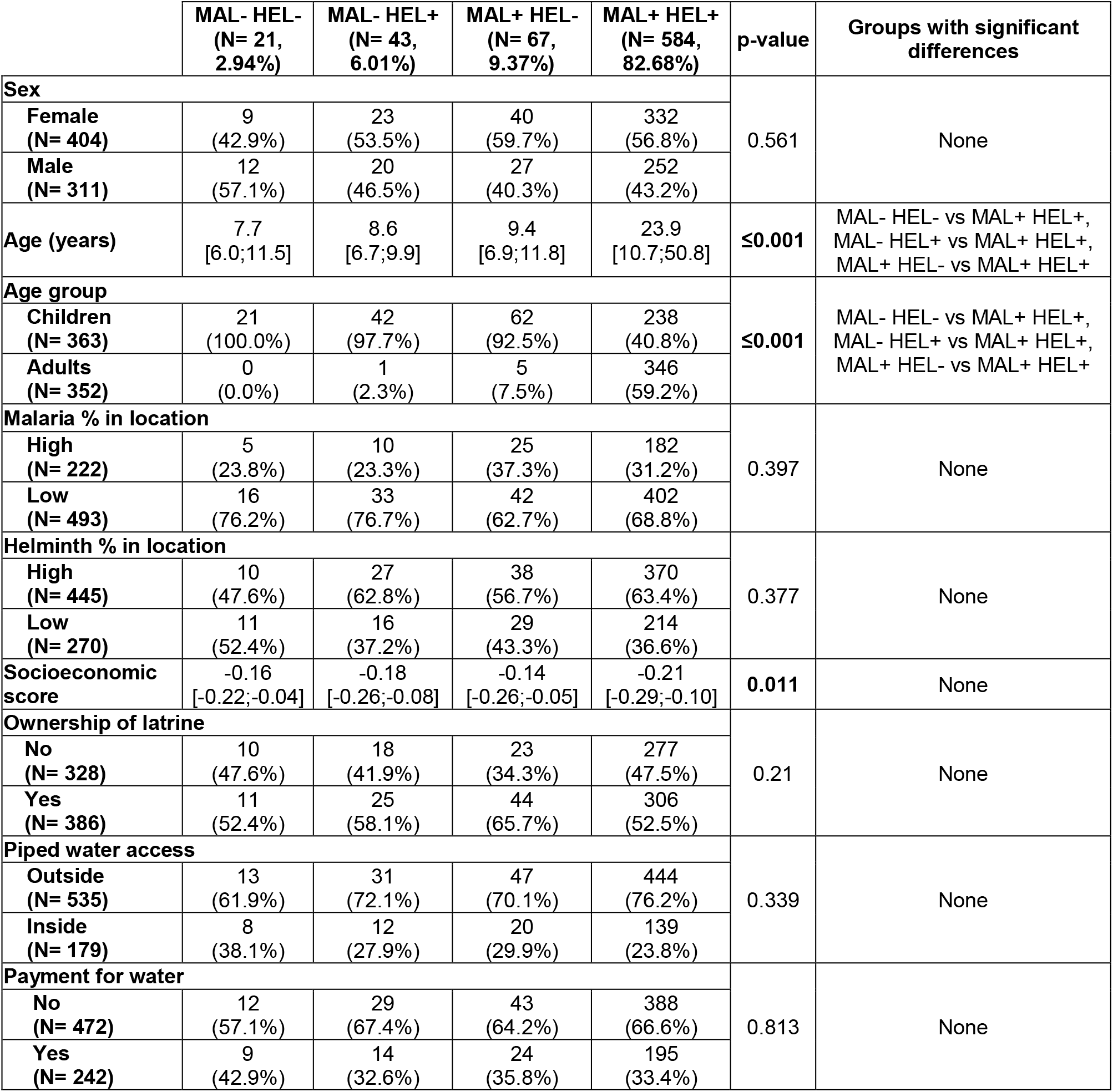
Demographic, socioeconomic, water and sanitation characteristics of the study population by exposure/infection group. Comparisons were made between study groups: no exposure/infection (MAL- HEL-), only exposure/infection with helminths (MAL- HEL+), only exposure/infection with *P. falciparum* (MAL+ HEL-) or coexposure/coinfection with *P. falciparum* and helminths (MAL+ HEL+). Malaria and helminth percentage of infection in each location of the study were calculated and participants were classified into groups of locations with higher or lower percentage of infection compared to the overall median by type of parasite. Total sample size was N= 715, except for the socioeconomic score, ownership of latrine, piped water access and payment for water that was N= 714.

|  | MAL- HEL-<br>(N= 21,<br>2.94%) | MAL- HEL+<br>(N= 43,<br>6.01%) | MAL+ HEL-<br>(N= 67,<br>9.37%) | MAL+ HEL+<br>(N= 584,<br>82.68%) | p-value | Groups with significant<br>differences |
| --- | --- | --- | --- | --- | --- | --- |
| Sex |  |  |  |  | 0.561 | None |
| Female<br>(N= 404) | 9<br>(42.9%) | 23<br>(53.5%) | 40<br>(59.7%) | 332<br>(56.8%) |  |  |
| Male<br>(N= 311) | 12<br>(57.1%) | 20<br>(46.5%) | 27<br>(40.3%) | 252<br>(43.2%) |  |  |
| Age (years) | 7.7<br>[6.0;11.5] | 8.6<br>[6.7;9.9] | 9.4<br>[6.9;11.8] | 23.9<br>[10.7;50.8] | ≤0.001 | MAL- HEL- vs MAL+ HEL+,<br>MAL- HEL+ vs MAL+ HEL+,<br>MAL+ HEL- vs MAL+ HEL+ |
| Age group |  |  |  |  | ≤0.001 | MAL- HEL- vs MAL+ HEL+,<br>MAL- HEL+ vs MAL+ HEL+,<br>MAL+ HEL- vs MAL+ HEL+ |
| Children<br>(N= 363) | 21<br>(100.0%) | 42<br>(97.7%) | 62<br>(92.5%) | 238<br>(40.8%) |  |  |
| Adults<br>(N= 352) | 0<br>(0.0%) | 1<br>(2.3%) | 5<br>(7.5%) | 346<br>(59.2%) |  |  |
| Malaria % in location |  |  |  |  | 0.397 | None |
| High<br>(N= 222) | 5<br>(23.8%) | 10<br>(23.3%) | 25<br>(37.3%) | 182<br>(31.2%) |  |  |
| Low<br>(N= 493) | 16<br>(76.2%) | 33<br>(76.7%) | 42<br>(62.7%) | 402<br>(68.8%) |  |  |
| Helminth % in location |  |  |  |  | 0.377 | None |
| High<br>(N= 445) | 10<br>(47.6%) | 27<br>(62.8%) | 38<br>(56.7%) | 370<br>(63.4%) |  |  |
| Low<br>(N= 270) | 11<br>(52.4%) | 16<br>(37.2%) | 29<br>(43.3%) | 214<br>(36.6%) |  |  |
| Socioeconomic<br>score | -0.16<br>[-0.22;-0.04] | -0.18<br>[-0.26;-0.08] | -0.14<br>[-0.26;-0.05] | -0.21<br>[-0.29;-0.10] | 0.011 | None |
| Ownership of latrine |  |  |  |  | 0.21 | None |
| No<br>(N= 328) | 10<br>(47.6%) | 18<br>(41.9%) | 23<br>(34.3%) | 277<br>(47.5%) |  |  |
| Yes<br>(N= 386) | 11<br>(52.4%) | 25<br>(58.1%) | 44<br>(65.7%) | 306<br>(52.5%) |  |  |
| Piped water access |  |  |  |  | 0.339 | None |
| Outside<br>(N= 535) | 13<br>(61.9%) | 31<br>(72.1%) | 47<br>(70.1%) | 444<br>(76.2%) |  |  |
| Inside<br>(N= 179) | 8<br>(38.1%) | 12<br>(27.9%) | 20<br>(29.9%) | 139<br>(23.8%) |  |  |
| Payment for water |  |  |  |  | 0.813 | None |
| No<br>(N= 472) | 12<br>(57.1%) | 29<br>(67.4%) | 43<br>(64.2%) | 388<br>(66.6%) |  |  |
| Yes<br>(N= 242) | 9<br>(42.9%) | 14<br>(32.6%) | 24<br>(35.8%) | 195<br>(33.4%) |  |  |

For subsequent analyses, we considered only helminth infection regardless of the species. Combining the diagnosis of exposure and/or infections, 91.05% and 87.69% of individuals had been exposed to and/or were infected with *P. falciparum* and helminths, respectively (**Table S1**), with the following breakdown into exposure/infection groups: no exposure/infection (MAL- HEL- [N= 21, 2.94%]), single exposure/infection (MAL- HEL+ [N= 43, 6.01%] or MAL+ HEL- [N= 67, 9.37%]) and coexposure/coinfection (MAL+ HEL+ [N= 584, 82.68%]) (**Table 1**). The median antibody levels for each group defined by exposure/infection followed a similar pattern but were lower compared to the same groups defined by current infection only (**Figure S2**). Using a combination of past exposure and current infection, we were able to capture a large proportion of exposed individuals not detected by current infection diagnosis (**Table S3**).

Females (N= 404) and males (N= 311) were equally distributed across exposure/infection study groups but there were significant differences in terms of age (**Table 1**). As expected, the MAL- HEL- group had the lowest age median (7.7 years), while the MAL+ HEL+ group had the highest age median (23.9 years) (**Table 1**). Further, the breakdown by age group revealed a very unbalanced distribution of children (5-14 years old) and adults (≥ 15 years old) across study groups, with the vast majority of adults in the MAL+ HEL+ group (**Table 1**).

The percentage of individuals with asymptomatic malaria and infection by any helminth was calculated for each of the 6 locations. Then, these were classified into “high” and “low” prevalence locations based on higher or lower percentage of infected individuals compared to the median percentage of infections from all locations (**Tables 1** and **S4**). No statistically significant differences were found by high or low location between study groups (**Table 1**).

Regarding socioeconomic, water and sanitation metrics, such as socioeconomic score (SES), ownership of latrine, accessibility to piped water inside the household and payment for water, there were no statistically significant differences between exposure/infection groups (**Table 1**).

### Association of demographic, SES, water and sanitation variables with antibody levels

Sex had a different impact depending on the immunoglobulin isotype. For antigen-specific IgG, there was a clear and consistent trend for higher antibody levels in females than males, which was statistically significant (p≤ 0.05) for IgG to EXP1, AMA1, As37, AyCp2, NaAPR1, NaGST1, NaSAA2 and MEA (**Figures 1** and **S3A**). For total IgE, there was a trend in the opposite direction, with males exhibiting higher levels (p= 0.087) (**Figures 1** and **S3A**). When differences by sex were assessed stratifying by age group, there were no differences for antigen-specific IgG (data not shown), whereas total IgE was statistically significantly higher in male compared to female children (**Figure S3B**).

**Figure 1.**
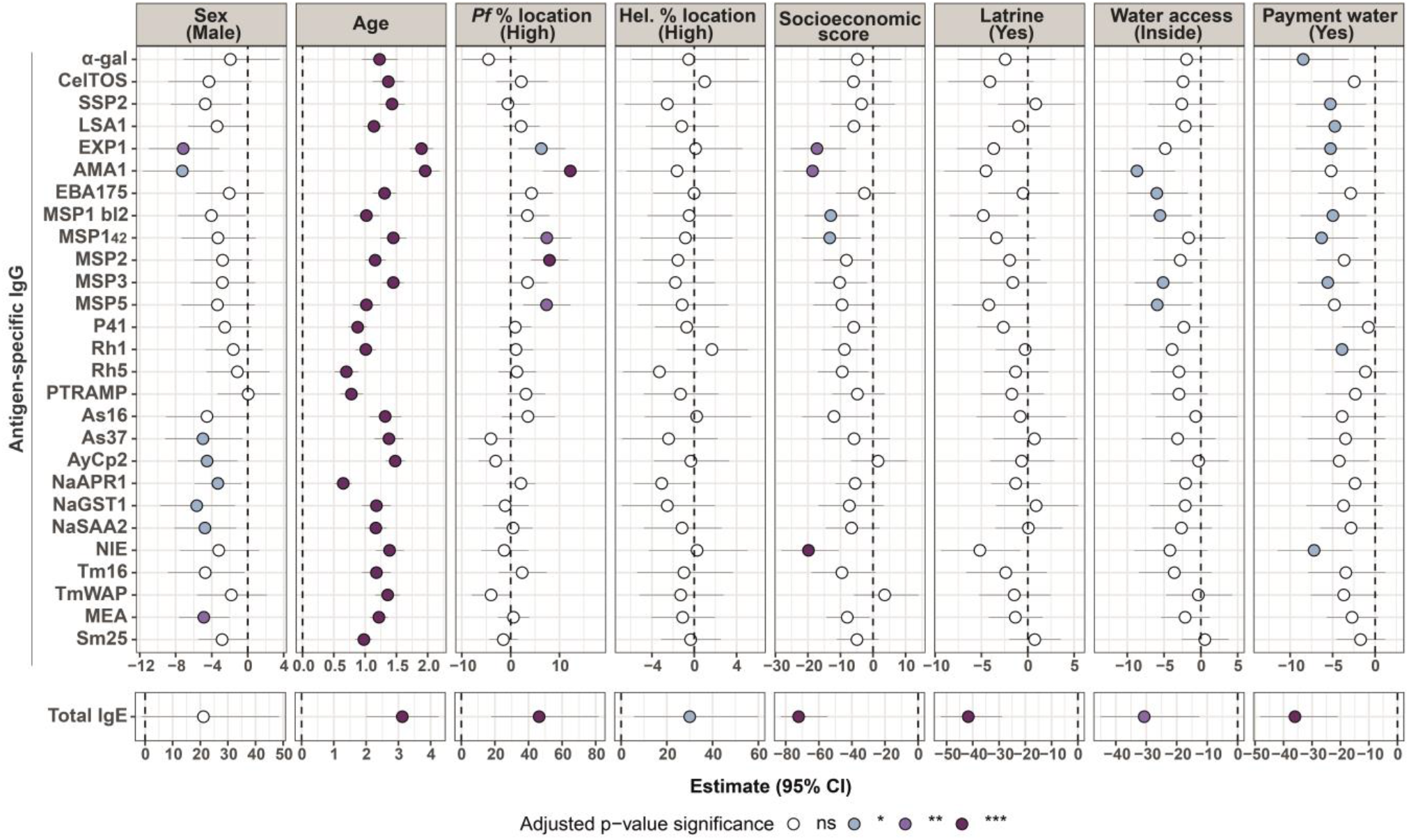
Association of demographic, socioeconomic, water and sanitation metrics with antibody levels in univariable linear regression models. Forest plots show the association of sex (reference: female), age, percentage of *P. falciparum* infected individuals by location (reference: locations with percentage lower than the median of all locations), percentage of helminth infected individuals by location (reference: locations with percentage lower than the median of all locations), socioeconomic score, owning a latrine (reference: no), piped water accessibility (reference: outside the household), and payment for water (reference: no) with specific IgG levels to *P. falciparum* and helminth antigens and total IgE levels. Univariable linear regression models were fitted to calculate the estimates (dots) and 95% confidence intervals (CI) (lines). The represented values are the transformed betas and CI in percentage. The color of the dots represents the p-value significance after adjustment for multiple testing by Benjamini-Hochberg, where ns= not significant, * = p-value ≤ 0.05, ** = p-value ≤ 0.01 and *** = p-value ≤ 0.001. Pf: *P. falciparum*; Hel.: Helminth.

There was a positive correlation between age and all antigen-specific IgG (rho= 0.31 – 0.64) and total IgE levels (rho= 0.2) (**Figures 1** and **S4**). IgG and IgE antibody levels rose until 15−24 years old and then remained relatively stable during the rest of adulthood (**Figure 2**).

**Figure 2.**
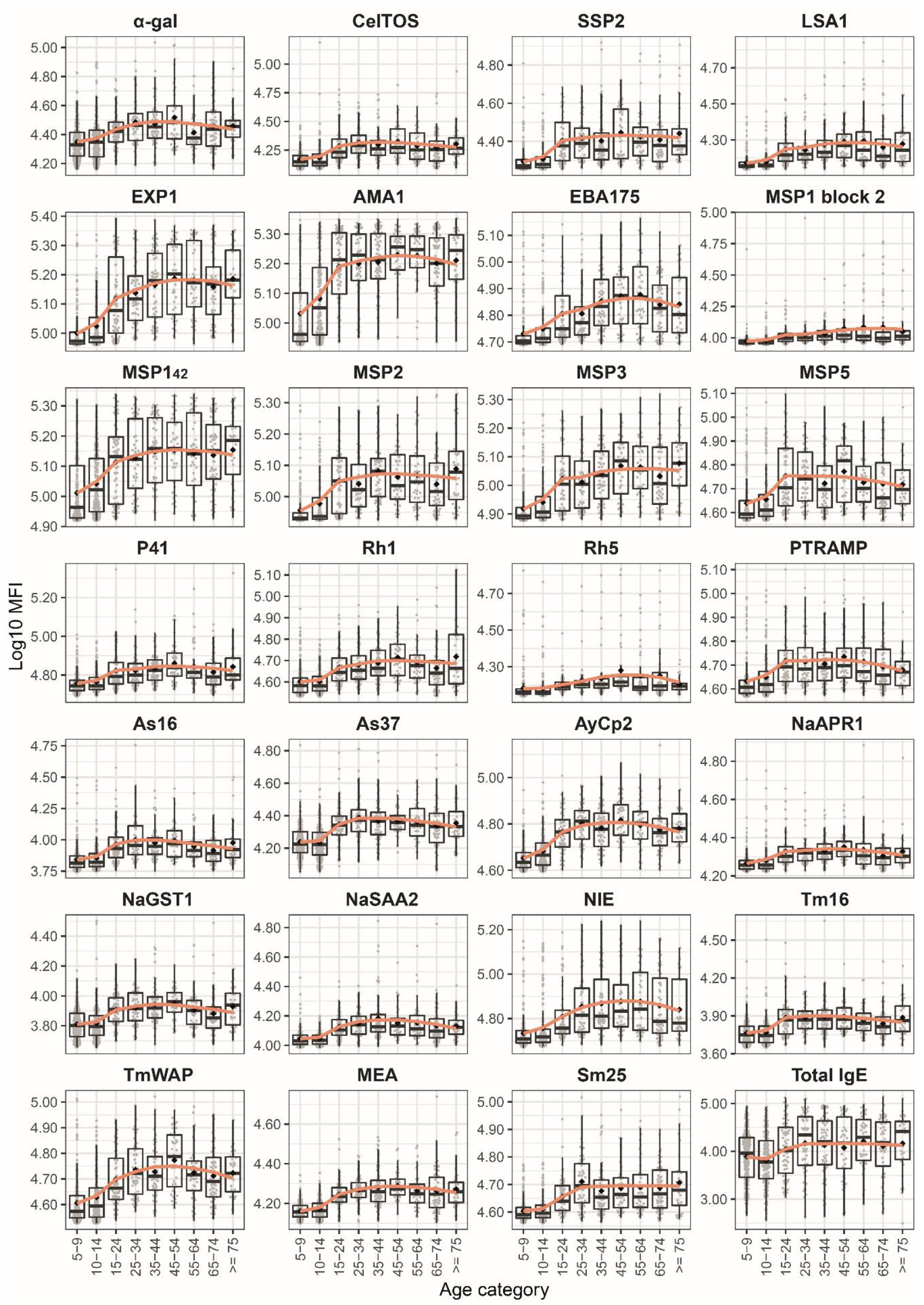
Antibody levels by age groups. Antigen-specific IgG levels against *Plasmodium falciparum* and helminth antigens and total IgE levels are expressed as log_10_-transformed median fluorescence levels (MFI). Age categories are grouped in 5 years for children and 10 years for adults and the boxplots represent the median (bold line), the mean (black diamond), the 1^st^ and 3^rd^ quartiles (box) and the largest and smallest values within 1.5 times the inter-quartile range (whiskers). Data beyond the end of the whiskers are outliers. The kinetics curves in orange were calculated using the locally estimated scatterplot smoothing (LOESS) method. *P. falciparum* antigens: α-gal, CelTOS, SSP2, LSA1, EXP1, AMA1, EBA175, MSP1 block 2, MSP1_42_, MSP2, MSP3, MSP5, P41, Rh5, PTRAMP. Helminth antigens: As16 and As37 (*Ascaris lumbricoides*), AyCp2, NaAPR1, NaGST1, NaSAA2 (Hookworm), NIE (*Strongyloides stercoralis*), Tm16 and TmWAP (*Trichuris trichiura*), MEA and Sm25 (*Schistosoma* spp).

Being from a location with a high prevalence of malaria was associated with higher total IgE levels (p≤ 0.001) and *P. falciparum*-specific IgG levels, specifically for EXP1, AMA1, MSP1_42_, MSP2 and MSP5 but not with helminth-specific IgG levels (**Figures 1** and **S5**). Living in a location with a higher burden of helminth infection was associated with higher total IgE levels (p≤ 0.05), but not with helminth-specific nor *P. falciparum*-specific IgG levels (**Figures 1** and **S6**).

SES was negatively association with IgG anti-EXP1, AMA1, MSP1 block 2, MSP1_42_, and NIE and total IgE (**Figure 1**). Overall, there was a weak but significant negative correlation (rho= -0.17 to - 0.1) between antigen-specific IgG levels and SES, whereas the negative correlation with total IgE levels was somewhat stronger (rho= -0.21) (**Figure S7**).

Regarding water and sanitation metrics, individuals with a latrine at home had significantly lower total IgE levels than individuals without a latrine at home (p≤ 0.001), while specific-IgG levels were not significantly associated (**Figures 1** and **S8**). Additionally, living in a household with piped water inside and paying for water access were also significantly associated with lower total IgE levels compared to living in a household with piped water in the yard (p = 0.002) or a household that did not pay for water (p≤ 0.001) (**Figures 1**). Piped water accessibility outside the household was associated with higher antigen-specific IgG levels against AMA1, EBA175, MSP1 block 2, MSP3 and MSP5 and no payment for water with IgG levels against α-gal, SSP2, LSA1, EXP1, MSP1 block 2, MSP1_42_, MSP3, Rh1 and NIE (p≤ 0.05) (**Figures 1, S9** and **S10**).

### Association of coexposure/coinfection with antibody levels

The MAL+ HEL+ group consistently showed higher IgG levels against all *P. falciparum* and helminth antigens and higher total IgE levels (**Figures 3A, 3B** and **S11**), higher magnitude (defined as the sum of antibody levels against *P. falciparum* or helminth antigens) (**Figure 3C**) and breadth of response (defined as the number of *P. falciparum* or helminth seropositive responses) (**Figure 3D**), compared to the MAL+ HEL- or MAL- HEL+ groups (p ≤ 0.001 except for IgE levels between MAL+ HEL+ and MAL-HEL+ where p= 0.033). Since all antibody levels increased with age, and there was a biased distribution of age groups across infection groups, this analysis was also performed stratified by age, where only the children had enough sample size in all the groups. The results in children confirmed the results in the whole cohort, with statistically significant differences in all comparisons except for IgG against highly immunogenic antigens such as AMA1, EBA175 and MSP1_42_, where after adjusting for multiple comparisons statistical significance disappeared, although a clear trend remained (**Figure S12**).

**Figure 3.**
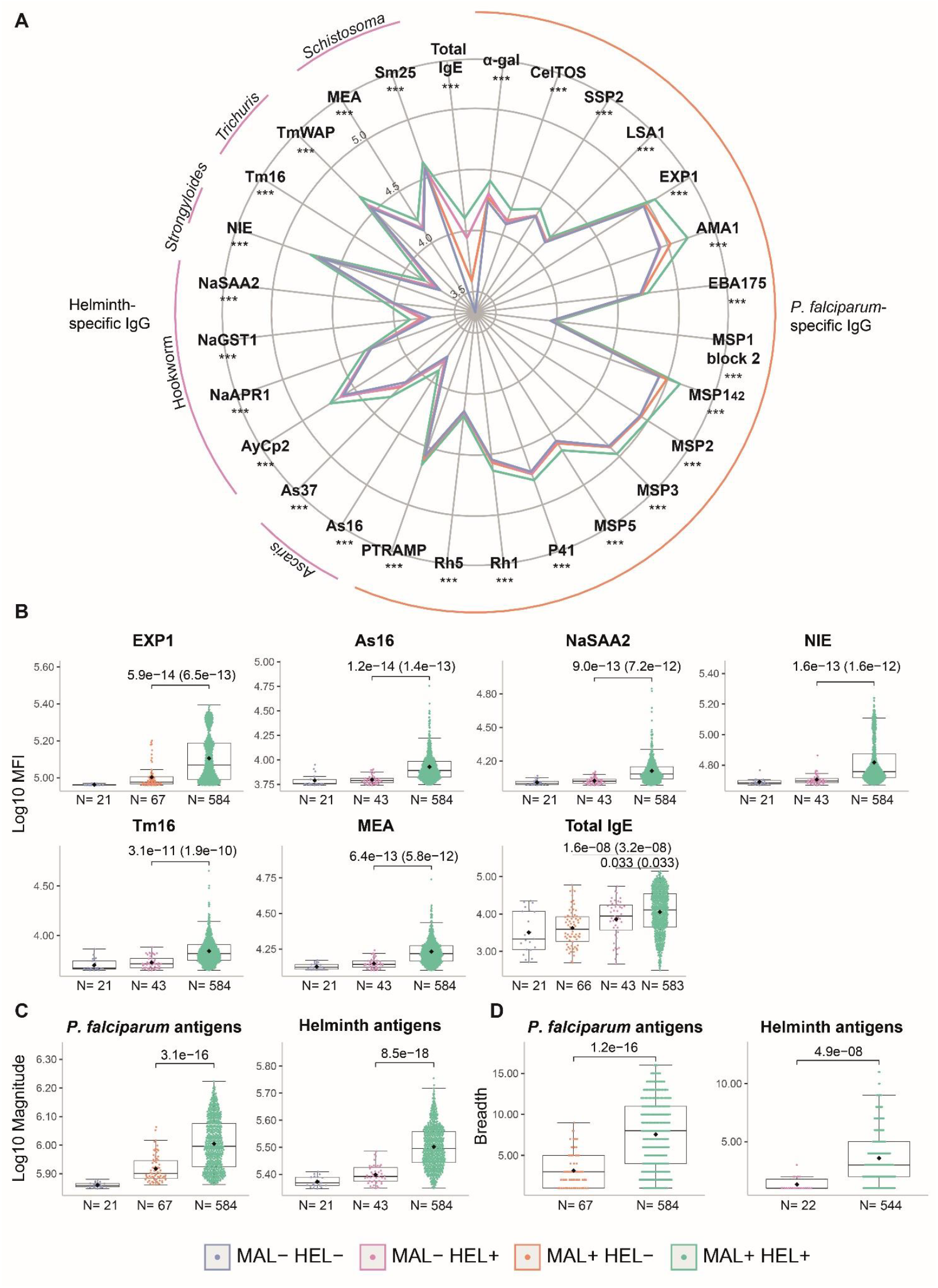
Comparison of antibody responses between exposure/infection groups. **(A)** Radarplot summarizing the antibody responses per exposure/infection group. In the vertices, the antibody responses are grouped by specific IgG against *Plasmodium falciparum* antigens, helminth antigens or total IgE. The colored lines represent the median of antibody levels expressed as log_10_-transformed median fluorescence levels (MFI). For *P. falciparum* and helminth antigen-specific IgG, the medians of the MAL+ HEL+ group were compared with the median of the MAL+ HEL- or MAL- HEL+ groups, respectively, by Wilcoxon rank sum test. For total IgE, the median of the MAL+ HEL+, MAL+ HEL- and MAL- HEL+ groups were compared by Kruskall-Wallis test. Statistically significant differences are indicated with asterisks (*** p ≤ 0.001). The boxplots compare infection groups based on **(B)** the log_10_-transformed MFI specific IgG levels against representative antigens per parasite species and total IgE, **(C)** the log_10_-transformed magnitude of response (sum of MFI), and **(D)** the breadth of response (number of seropositive antigen-specific IgG responses). The boxplots represent the median (bold line), the mean (black diamond), the 1^st^ and 3^rd^ quartiles (box) and the largest and smallest values within 1.5 times the inter-quartile range (whiskers). Data beyond the end of the whiskers are outliers. Statistical comparison between groups was performed by Wilcoxon rank sum test and the exact p-values are shown. Adjusted p-values by the Benjamini-Hochberg approach are shown in brackets. Study groups are shown in colors: no exposure/infection (violet) (MAL- HEL-), only exposure/infection with helminths (pink) (MAL- HEL+), only exposure/infection with *P. falciparum* (orange) (MAL+ HEL-) and coexposure/coinfection with *P. falciparum* and helminths (green) (MAL+ HEL+). The rest of the antigens are showed in Supplementary Figure 10.

In multivariable analysis, linear regression models adjusted by their corresponding covariables revealed that the MAL+ HEL+ group had significantly higher antibody levels against most of the antigens, magnitude and breadth of response compared to single infection/exposure groups (MAL+ HEL- or MAL- HEL+) (**Figure 4**).

**Figure 4.**
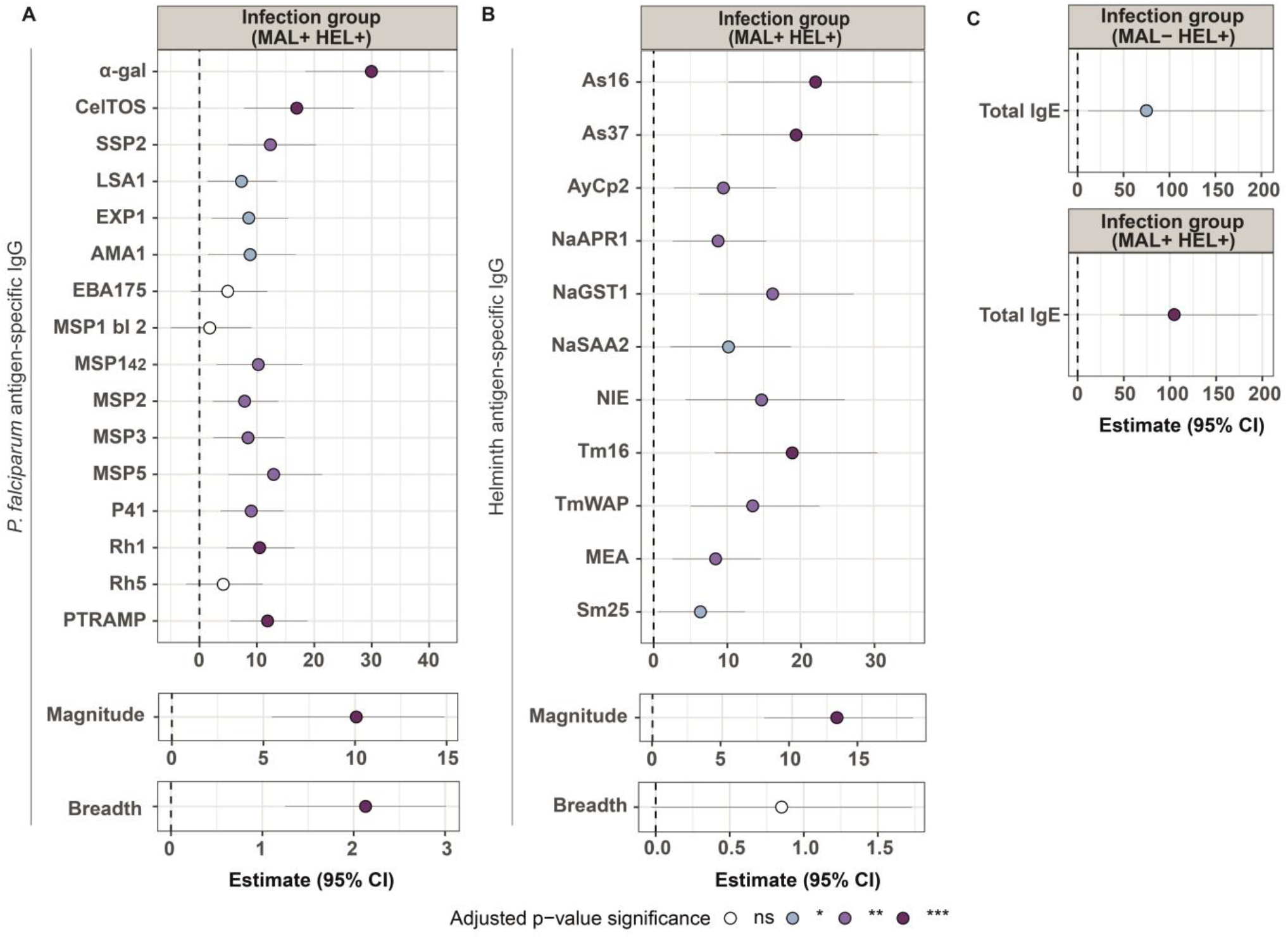
Association of exposure/infection group with antibody levels in multivariable linear regression models. Forest plots show the association of the MAL+ HEL+ group with **(A)** IgG levels to *P. falciparum* antigens, magnitude and breadth of response (reference: MAL+ HEL- group) or **(B)** IgG levels to helminth antigens, magnitude and breadth of response (reference: MAL- HEL+ group). In **(C)**, the association of MAL+ HEL+ group or the MAL- HEL+ group with IgE levels was assessed (reference: MAL+ HEL- group). The magnitude of response was calculated as the sum of all specific IgG levels to the different antigens belonging to *P. falciparum* or helminths. The breadth was defined as the number of seropositive antigen-specific IgG responses. Multivariable linear regression models were fitted to calculate the estimates (dots) and 95% confidence intervals (CI) (lines). The represented values are the transformed betas and CI in percentage for IgG levels to antigens, total IgE and magnitude of response (log-linear models), and the untransformed betas and CI for the breadth of response (linear-linear models). The color of the dots represents the p-value significance after adjustment for multiple testing by Benjamini-Hochberg, where ns= not significant, * = p-value ≤ 0.05, ** = p-value ≤ 0.01 and *** =  p-value ≤ 0.001. Models were adjusted by age and percentage of *P. falciparum* infected individuals by location for *P. falciparum* antigens, age for helminth antigens, and sex, percentage of helminth infected individuals by location, ownership of latrine, piped water accessibility, payment for water and socioeconomic score for total IgE. MSP1 bl 2: MSP1 block 2. MAL+ HEL+: coexposure/coinfection with *P. falciparum* and helminths; MAL+ HEL-: only exposure/infection with *P. falciparum;* MAL- HEL+: only exposure/infection with helminths.

For most *P. falciparum* antigens, IgG levels increased between 5% and 15% in the MAL+ HEL+ group compared to the MAL+ HEL- group, although it reached a 17% increase for CelTOS (95% CI = 7.750 – 26.932) and a 30% for α-gal (95% CI = 18.438 - 42.570) (**Figure 4A**). The only antigens without a significant increase associated with coinfection were EBA175, MSP1 block 2 and Rh5 (**Figure 4A**). The MAL+ HEL+ group resulted in an increase of up to a 10% (95% CI = 5.462 - 14.889) in the magnitude of response, and an increase of 2 in the total number of seropositive responses (95% CI = 1.254 - 3.001) (**Figure 4A**).

Regarding helminth antigens, the increase in IgG levels in the MAL+ HEL+ group compared to the MAL- HEL+ group ranged from 6% for Sm25 (95% CI = 0.624 - 12.463) to 22% for As16 (95% CI = 10.213 - 35.249) (**Figure 4B**). The magnitude increased up to a 13% (95% CI = 7.995 - 18.684) while the breadth was not significantly affected (0.852, 95% CI = -0.030 - 1.734) (**Figure 4B**).

For total IgE levels, MAL- HEL+ or MAL+ HEL+ groups were significantly associated with a 74% (95% CI = 6.193 – 185.409) or 106% (95% CI = 46.453 – 191.300) increase in total IgE levels, respectively, compared to the MAL+ HEL- group (**Figure 4C**).

### Association of magnitude of response and total IgE with antibody levels

We also performed additional multivariable models in the whole cohort using total IgE or the magnitude of response against malaria and helminth antigens as predictor exposure variables instead of exposure/infection groups (**Figure S13** and **S14**). Total IgE was significantly positively associated with the magnitude and breadth of response and IgG levels against *P. falciparum* and helminth antigens (**Figure S13**). The magnitude of response against helminth antigens was also positively associated with IgG levels against *P. falciparum* antigens. Magnitude of response against *P. falciparum* antigens was also positively associated with IgG levels against helminth antigens, and both magnitudes with total IgE levels (**Figure S14**). Lastly, as expected, the magnitude of response against helminth antigens was more associated with an increase in total IgE levels (31%, 95% CI = 22.314 - 39.504) than the magnitude of response against *P. falciparum* antigens (9%, 95% CI = 2.778 – 15.956) (**Figure S14**).

### Association of coexposure/coinfection with susceptibility to infection

Logistic regression analysis adjusted by their corresponding covariables showed a consistent trend for higher odds of having malaria or helminth infections if exposed to or currently having the other parasite type (**Table S5**). In support of this trend, a higher magnitude of antibody response to helminth or *P. falciparum* antigens was significantly associated to having *P. falciparum* or helminth infection, respectively (**Figure 5**). In addition, there was a positive association between being exposed/infected with *P. falciparum* and the number of helminth species exposed/infected with (**Table 2**). Furthermore, there was a statistically significant higher proportion of individuals exposed/infected with multiple helminth species in the MAL+ HEL+ group than in the MAL- HEL+ group (p≤ 0.001) (**Table S2**).

**Table 2.** Association between exposure/infection with Plasmodium falciparum and number of helminth species defined by exposure/infection. In the linear regression model, the reference category was MAL-. The models were adjusted by covariables selected based on their significant association with the outcome variable in univariable models: * log_10_ age *** log_10_ age, sex and socioeconomic score N: number of individuals, OR: odds ratio, CI: Confidence interval, MAL+: *Plasmodium falciparum* positive infection or exposure/infection, MAL-: *P. falciparum* negative infection or exposure/infection.

| Outcome |  | Predictor | Estimate | 95% CI | p-value |
| --- | --- | --- | --- | --- | --- |
| Multivariable logistic regression |  |  |  |  |  |
| MAL+<br>(N) | MAL-<br>(N) | Number of<br>helminth<br>species | OR*= 1.607 | [1.214 – 2.167] | ≤ 0.001 |
| 67 | 21 | 0 |  |  |  |
| 109 | 26 | 1 |  |  |  |
| 144 | 12 | 2 |  |  |  |
| 148 | 4 | 3 |  |  |  |
| 114 | 0 | 4 |  |  |  |
| 69 | 0 | 5 |  |  |  |
| Multivariable linear regression |  |  |  |  |  |
| Number of<br>helminth<br>species |  | MAL+ | β***= 0.637 | [0.316 – 0.958] | ≤ 0.001 |

**Figure 5.**
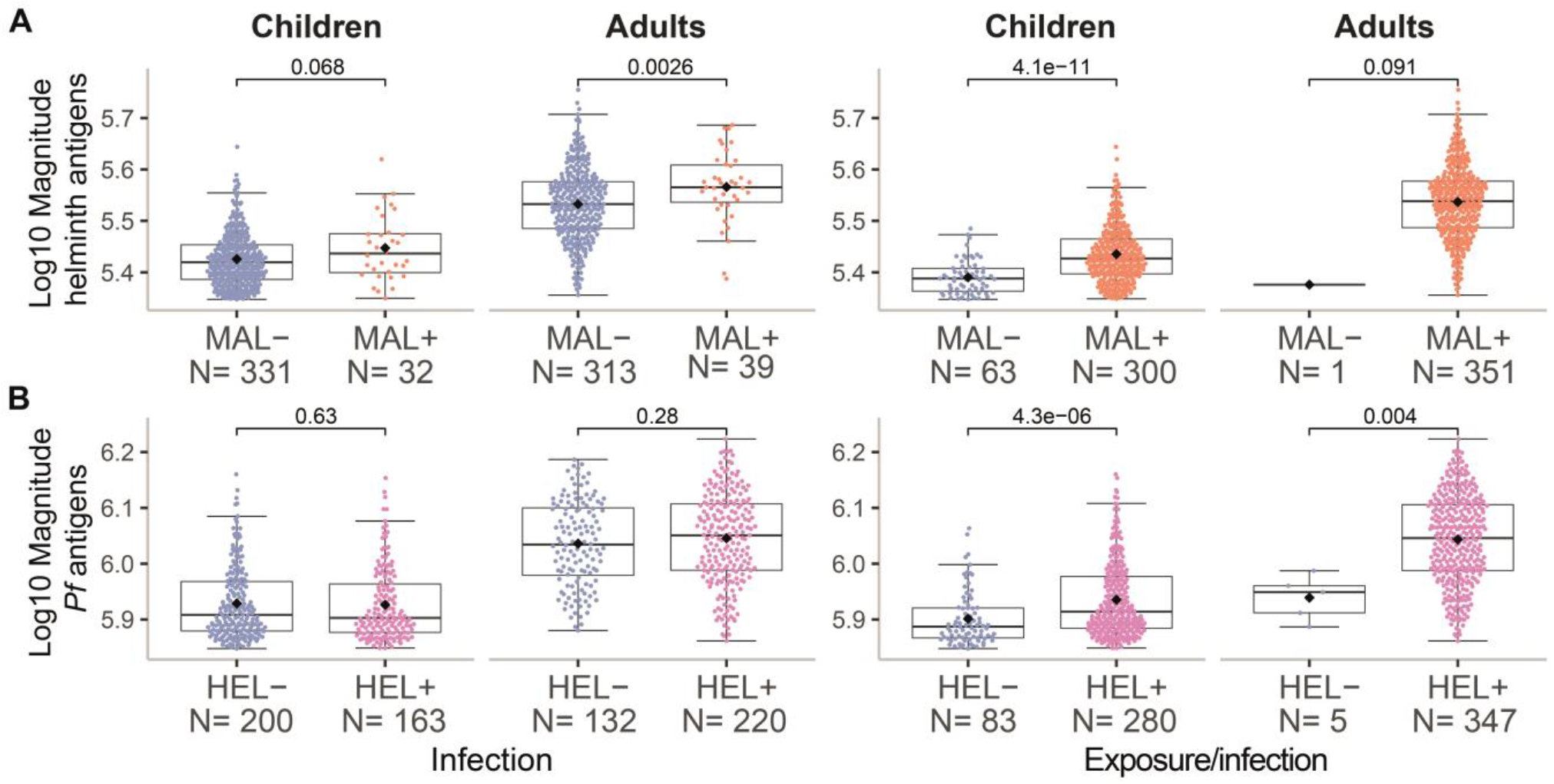
Magnitude of response by infection or exposure/infection to malaria or helminths. The magnitude of response represents the sum of all specific IgG responses against **(A)** helminth or **(B)** *P. falciparum* antigens. The X-axis shows the classification of samples into negative (-) or positive (+) based on either past exposure and/or present infection (detected by serology and molecular and/or microscopic diagnosis) or present infection (detected by molecular and/or microscopic diagnosis) for **(A)** malaria (MAL) or **(B)** helminths (HEL). The boxplots represent the median (bold line), the mean (black diamond), the 1^st^ and 3^rd^ quartiles (box) and the largest and smallest values within 1.5 times the inter-quartile range (whiskers). Data beyond the end of the whiskers are outliers. Statistical comparison between groups was performed by Wilcoxon rank sum test and the exact p-values are shown.

### Association of coexposure/coinfection with parasite burden

Next, we evaluated whether being exposed and/or having a coinfection had an impact on parasite burden. In the case of malaria, 93% of the people with current *P. falciparum* infection were coexposed/coinfected with helminths (66/71), leaving the single infection group with a much-reduced sample size. In this case, the coexposure/coinfection group was not significantly associated (p= 0.34) with *P. falciparum* parasitemia (parasites/μL) (**Figure 6A**). However, since immunity develops over time and most of the coexposed/coinfected individuals were adults, we stratified by age group, which resulted in a trend (p= 0.062) for higher parasitemia in the coexposure/coinfection group compared to the malaria only group in children (**Figure 6A**). In the case of helminths, coexposure/coinfection had a significant protective effect on parasite burden (expressed as qPCR Ct values) of *T. trichiura* (p≤ 0.01), also after stratification by age group (p≤ 0.05) (**Figure 6B**). For the remainder of the helminth species there was no significant association between coexposure/coinfection and parasite burden (**Figure S15**).

**Figure 6.**
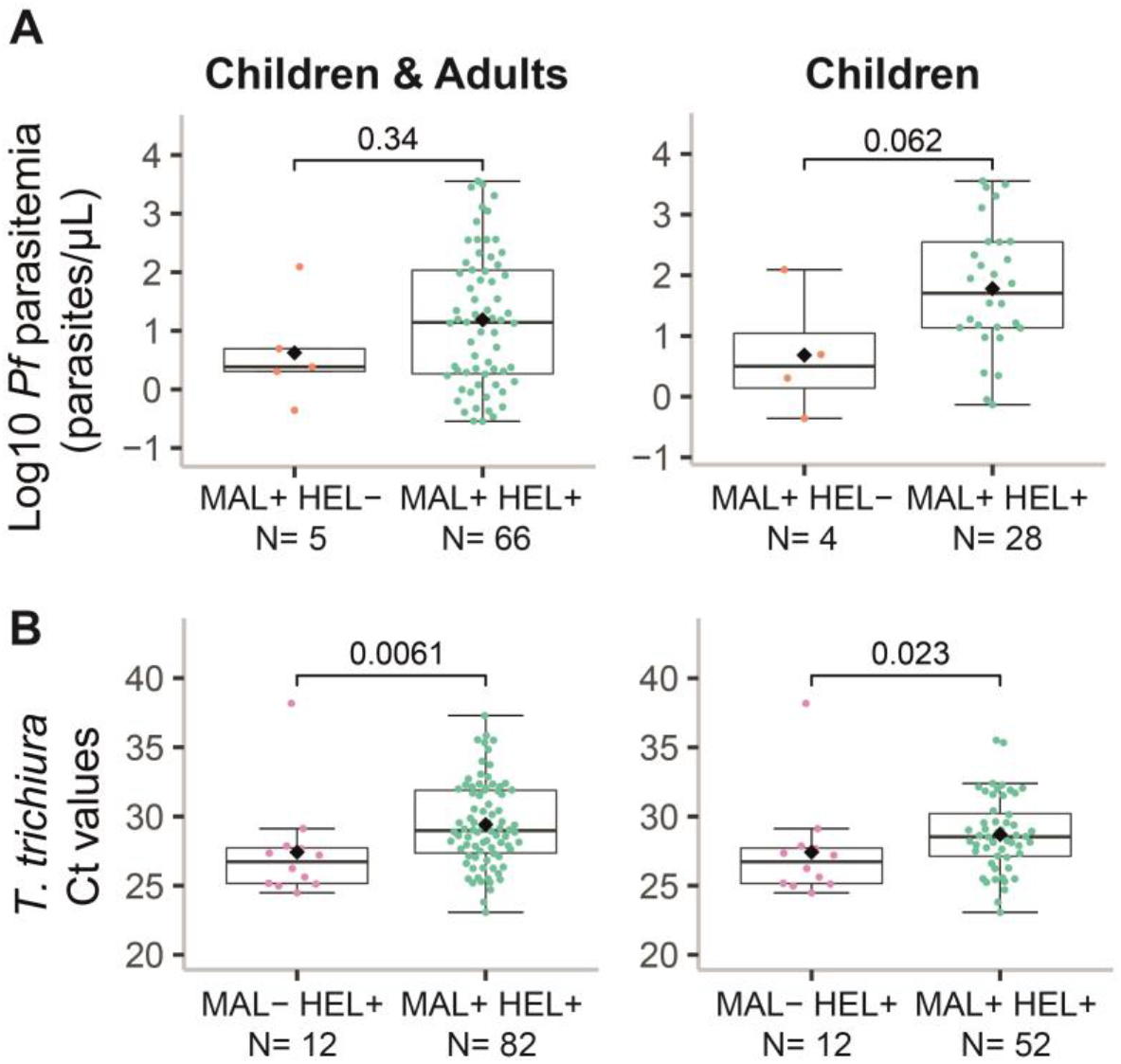
Parasite burden by exposure/infection groups. The levels of parasite burden are represented as **(A)** the log_10_-transformed parasitemia of *Plasmodium falciparum* (parasites/μL) or **(B)** the *Trichuris trichiura* Ct values, and are compared between the coinfection group (MAL+ HEL+) and the only malaria group (MAL+ HEL-) for *P. falciparum,* or the only helminths group (MAL- HEL+) for *T. trichiura*. For each parasite, data for all individuals with parasite burden information or only children are displayed. The boxplots represent the median (bold line), the mean (black diamond), the 1^st^ and 3^rd^ quartiles (box) and the largest and smallest values within 1.5 times the inter-quartile range (whiskers). Data beyond the end of the whiskers are outliers. Statistical comparison between groups was performed by Wilcoxon rank sum test and the exact p-values are shown. See supplementary Figure S15 for the plots of the rest of parasites.

### Association of antibody levels with parasite burden

To investigate the associations of antibody responses with parasite burden and their possible involvement in protection, we performed linear regression models adjusted by age. For *P. falciparum* parasitemia, there was a significant (raw p≤ 0.05) positive association of specific IgG responses to CelTOS (17.793, 95% CI = 2.99 – 35.634), SSP2 (22.229, 95% CI = 1.38 – 47.361), AMA1 (34.277, 95% CI = 2.410 – 76.060), MSP5 (26.824, 95% CI = 5.440 – 52.545) and the breadth of response to *P. falciparum* antigens (20.838, 95% CI = 1.111 – 44.415) only before adjustment for multiple comparisons (**Figure 7**).

**Figure 7.**
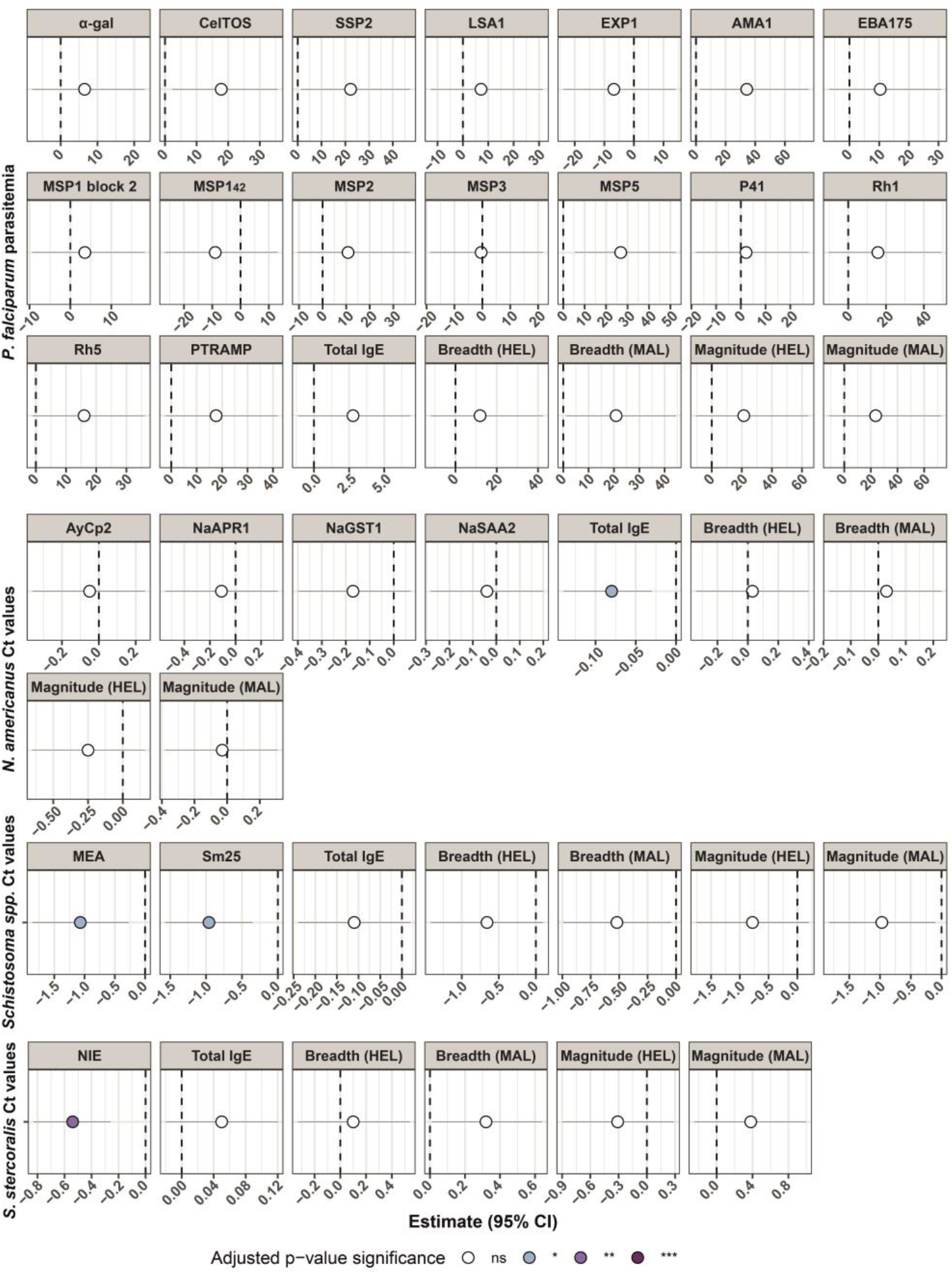
Association of antibody responses with parasite burden in multivariable linear regression models. Forest plots show the association of antibody responses on parasite burden represented as the log_10_-transformed parasitemia (parasites/μL) of *Plasmodium falciparum* or the Ct values of *Schistosoma* spp., *Necator americanus* and *Strongyloides stercoralis*. For each parasite, linear regression models adjusted by age were fitted, where specific IgG levels against the antigens from the corresponding parasite, total IgE levels and magnitude (sum of antigen-specific IgG levels) and breadth (number of seropositive antigen-specific IgG responses) of response were the predictor variables, and the parasite burden was the outcome variable. The estimates (dots) and 95% confidence intervals (CI) (lines) of the models were transformed into percentages for *P. falciparum* parasitemia models (log-log or log-linear models). For helminths models, when the predictor variables were antibody levels or magnitude of response (linear-log models), the betas and 95% CI were transformed to represent the additive effect on the Ct values of a 10% increase in the predictor variable. For the breadth of response in helminths models (linear-linear models), the untransformed betas were used. The color of the dots represents the adjusted p-value significance, where ns= not significant, * = p-value ≤ 0.05, ** = p-value ≤ 0.01 and *** = p-value ≤ 0.001. See supplementary Figure S16 for *Ascaris lumbricoides* and *Trichuris trichiura* models.

As for helminths, we found significant negative associations of qPCR Ct values with total IgE (-0.085, 95% CI = -0.141 – -0.028) for hookworm, specific IgG to Sm25 (-0.963, 95% CI = -1.573 – -0.353) and MEA (-1.065, 95% CI = -1.862 – -0.268) for *Schistosoma* spp. and NIE for *S. stercoralis* (-0.542, 95% CI = -0.828 – -0.256) (**Figure 7**). These negative associations with Ct values are indicative of a positive association with parasite burden. We found no association between *A. lumbricoides* and *T. trichiura* antibody levels and Ct values (**Figure S16**).

## DISCUSSION

In this study we found that 83% of a cohort of children and adults from Southern Mozambique was infected or had been exposed to *P. falciparum* and helminths, mainly hookworm. Our main finding was that coexposure/coinfection was associated with increased antibody levels, magnitude and breadth of response to both types of parasites compared to the single exposure/infection groups.

We initially expected a deviation in the coexposure/coinfection group towards one of the branches of the immune response, which would result in reduced antibody production of the specificity and type required for one or other type of infection. On the contrary, we found that antigen-specific IgG and total IgE responses were higher in the coexposure/coinfection group compared to the single exposure/infection groups. For *P. falciparum*, this is consistent with previous studies that reported increased *P. falciparum*-specific IgG1, IgG2 and IgG3 responses in coinfection with *S. haematobium* (Diallo et al., 2010; Tokplonou et al., 2020). In contrast, other studies have reported a detrimental effect of coinfection with helminths in the humoral response to malaria with a reduction in *P. falciparum*-specific IgG (Ateba-Ngoa et al., 2016), IgG1 and IgG3 (Courtin et al., 2011; Roussilhon et al., 2010), and an increase in the non-cytophilic IgG4 (Roussilhon et al., 2010). As for helminths, the effect of *P. falciparum* infection on antibody levels has been mainly restricted to *Schistosoma* spp. and, in line with our results, coinfection with *P. falciparum* has been associated with an increase in both antigen-specific IgG and IgE responses to *Schistosoma* spp. antigens compared to the single infection group (Remoue et al., 2003). Does this increase mean there is a synergy at the immune humoral compartment or does it simply reflect an increased susceptibility, and therefore exposure to infections in the coinfection group?

In support of the first option, antibody production requires the aid of classical Th2-like cytokines such as IL-4 (Kotowicz and Callard, 1993; Troye-Blomberg et al., 1990). The fact that both types of parasites are able to induce Th2 responses, at least at some point during the course of infection, could favor higher levels of antibodies. However, whether these antibodies are protective or not needs to be evaluated in longitudinal studies and by looking at the subclass composition of the IgG responses. A Th2 milieu could favor the production of antibodies, but without the appropriate Th1 balance, production would be expected to be skewed towards the non-cytophilic IgG2 and IgG4 (Druilhe et al., 2005) and IgE, which are suggested to increase the risk to *P. falciparum* infection or severe disease (Bouharoun-Tayoun and Druilhe, 1992; Perlmann et al., 1994). In fact, low levels of IL-4 are required for induction of IgG1, IgG2 and IgG3. In contrast, high levels of IL-4 had no effects or even suppressed those IgG subclasses while IgE and IgG4 were enhanced (Kotowicz and Callard, 1993). This hypothesis, however, is not supported by data of increase in *P. falciparum*-specific IgG1 and IgG3 during coinfection with *S. haematobium* (Diallo et al., 2010). Similarly, *P. falciparum* coinfection with hookworm increased IgG1 and IgG3 levels to the GMZ2 vaccine and related antigens, while deworming treatment reduced these IgG1 and IgG3 levels (Amoani et al., 2021). On the other hand, following the synergy hypothesis, our results could be beneficial for helminth immunity assuming that IgE has a protective effect. However, in our study, total IgE levels had a significant positive association with hookworm burden and with all antigen-specific IgG levels, indirectly suggesting total IgE as a marker of exposure/infection rather than protection. Other studies have also reported a positive correlation between parasite burden or helminth-specific IgG levels and total IgE levels (Cooper et al., 2008; Mangano et al., 2020; Mulu et al., 2014).

In addition to higher antibody levels, coexposed/coinfected individuals had higher magnitude and breadth of response to *P. falciparum* and helminth antigens compared to single exposure/infection groups. A possible explanation is the ability of *P. falciparum* and helminths to induce polyclonal B cell responses (Donati et al., 2004; Harris and Gause, 2011), which is particularly evident for IgE during helminth infections (Maizels and Yazdanbakhsh, 2003). This phenomenon consists in the induction and differentiation of antibody-secreting cells from different B cell clones, regardless of their antigen specificity. While this may play an important role in defense against infections by enhancing natural antibody production, it could also be a strategy to avoid host-specific immune responses by “diluting” antibodies against important epitopes for protection (Montes et al., 2007). Either way, bystander antigens in coinfections could be affected by this spillover effect (Maizels and Yazdanbakhsh, 2003).

Nonetheless, our results point towards another more plausible option, which is that antibody responses are a consequence of an increased exposure due to higher susceptibility to infection in the coexposure/coinfection group. Firstly, antibody levels were positively associated with parasite burden, indicating that they reflect exposure rather than protection. These antibody responses that reflect exposure were increased in the coexposure/coinfection compared to single exposure/infection groups. We checked whether differences in the proportion of current infections between groups could be the cause due to boosting of the immune response, but there were no differences between groups except for *T. trichiura*. In fact, the proportion of *T. trichiura* current infection was higher in the helminth single exposure/infection group, but even so *T. trichiura* specific IgG levels were higher in the coexposure/coinfection group.

Secondly, there was a trend for higher odds of *P. falciparum* or helminths infection if coinfected or exposed to the other type of parasite. In fact, when using the magnitude of response to *P. falciparum* or helminth antigens as a surrogate of exposure, we found that *P. falciparum* exposed/infected individuals had a significantly higher exposure to helminths and *vice versa*. In line with this, others were able to confirm that infection with helminths increases the odds of *P. falciparum* infection (Afolabi et al., 2021; Babamale et al., 2018; Degarege et al., 2016; Nacher et al., 2002b). This is consistent with our finding that exposure/infection to an increasing number of helminth species augmented the odds of being infected or exposed to *P. falciparum*. Several studies have revealed similar results in terms of higher *P. falciparum* infection risk or density with increasing number of helminth species (Babamale et al., 2018; Degarege et al., 2012, 2009; Nacher et al., 2002b). In relation to this, one study indicated that helminth polyparasitism is more important than the effects of individual species when driving immunoregulatory effects (Turner et al., 2008). This supports the idea of higher susceptibility to *P. falciparum* infection, and in fact to any infection, with increasing number of helminths and aligns with our observation of higher proportion of exposure/infection with multiple helminth species in the coexposure/coinfection group compared to the only helminths group. This observation is equally consistent with the concept of hyper-susceptible hosts, who are individuals infected to a greater degree as a result of exposure to pathogens or underlying conditions.

Thirdly, there was a trend for higher *P. falciparum* burden in the coexposure/coinfection group compared to the *P. falciparum* single exposure/infection group in children, which is consistent with other reports for *P. falciparum* (Degarege et al., 2016) and *P. vivax* (Easton et al., 2020), but contrary to others (Amoani et al., 2019; Tokplonou et al., 2020). The only significant association for helminths with regards to parasite burden was for *T. trichiura*, where having been exposed to or currently infected with *P. falciparum* seemed to have a protective effect.

Therefore, if higher antibody levels simply reflect higher exposure, is there any underlying factor that renders these individuals in the coexposure/coinfection group more prone to multiple infections? It has been suggested that factors favoring coinfections may be common socioeconomic or environmental factors rather than true immunological interactions (Brooker et al., 2007; Mwangi et al., 2006). One of the strengths of this study is that we accounted for socioeconomical and water and sanitation factors, besides demographic variables, that could influence the exposure to the parasites. Among all variables, age had the strongest association with antibody levels. We clearly showed that the acquisition of immunity is similar across all antibody responses, with levels rising during childhood and stabilizing during adulthood. This is consistent with a cumulative exposure over time, which resulted in a clear age bias whereby most of the adults were in the coexposure/coinfection group. A study that detected IgG against some of the helminth antigens measured in this study (NIE, Sm25) and other *P. falciparum* antigens also observed similar age trends in Kenya (Njenga et al., 2020). Socioeconomic and water and sanitation metrics had the strongest associations with total IgE, with higher SES and improved water and sanitation conditions showing the lowest IgE levels. This is consistent with a higher exposure to helminths, potent inducers of IgE, under poor sanitation and hygiene conditions, considering that transmission of STH and *Schistosoma* spp. relies on fecal spread of eggs and larvae. It is noteworthy that also specific IgG levels against some *P. falciparum* and helminth antigens were affected by these variables, suggesting shared transmission risks, such as fecal contaminated water in the household surroundings that can serve as a reservoir of helminth eggs and mosquito larvae. However, only age was significantly higher in the coexposure/coinfection group compared to the rest. Therefore, immunological permissiveness seems to be a likely cause for the observed increase in antibody responses. Nonetheless, we cannot rule out other underlying conditions, such as malnutrition, that could make this group of individuals more susceptible to infections.

Total IgE levels were, as expected, associated with locations with high prevalence of helminths but surprisingly also with locations with high prevalence of *P. falciparum*. However, this could be because the location with the highest percentage of helminth infected individuals also had a high percentage of *P. falciparum* infected individuals. Nonetheless, it is remarkable that in the absence of infection and exposure to helminths, *P. falciparum* seems to be an inducer of total IgE. Additionally, in the coexposure/coinfection group, total IgE levels were also higher than in the helminth single exposure/infection group. This suggests that *P. falciparum* also contributes to these IgE levels, since exposure to or coinfection with *P. falciparum* does not seem to increase helminth burden. Interestingly, another study has also found elevated total IgE levels in malaria patients regardless of helminth coinfection (Mulu et al., 2014). IgE is a neglected isotype in the field of malaria, with limited studies focusing on it. However, there is evidence that *P. falciparum* indeed induces IgE but the role it plays in protection or susceptibility remains to be elucidated. Some have argued it has a protective role (Bereczky et al., 2004; Duarte et al., 2012, 2007; Farouk et al., 2005) while others indicate that it is involved in malaria pathogenesis (Calissano et al., 2003; Maeno et al., 2000; Perlmann et al., 1994, 2000, 1997; Seka-Seka et al., 2004; Tangteerawatana et al., 2007). Total IgE seems to be a good proxy of a Th2 polarization and, due to its short half-life when free in serum, it also serves as a proxy of recent infection with Th2 inducers such as helminths (Geha, 1984). In our linear models, IgE was positively associated with *P. falciparum*-specific IgG levels that have been shown to reflect exposure rather than protection. This indirectly supports the idea of a non-protective role of IgE in terms of *P. falciparum* infection, as it would be associated with increased exposure. However, we could not asses its implication in clinical outcomes or associations for *P. falciparum* antigen-specific IgE, which might be different than those of total IgE (Duarte et al., 2007) probably due to the polyclonal, non-specific nature of total IgE responses (Wang et al., 2011). A possible explanation for high IgE levels in malaria comes from the observation that establishment of antigen-specific IgE responses during the early years of life primes the immune system to produce IgE responses upon antigen contact without the need of cytokines controlling class switch to IgE (Kotowicz and Callard, 1993). This might be the case of people in the coexposure/coinfection group infected with *P. falciparum* and previously primed into a predominant Th2 profile by helminths.

A caveat of this study is the inability to determine causality due to its cross-sectional nature. A longitudinal design, on the other hand, would be more appropriate to determine whether people with *P. falciparum* or helminths are more susceptible to subsequent infections. Furthermore, this study only included asymptomatic malaria cases, which may be hindering the detection of important associations with disease severity and masking the true burden of coinfections. Nonetheless, even with only asymptomatic cases, we were able to identify significant associations that suggest a higher susceptibility to infection. Therefore, we expect that the inclusion of symptomatic cases would only accentuate these associations. An additional limitation is the pooling of all helminth species together due to small sample size of the groups at species level, possibly resulting in masked effects of individual species (Abbate et al., 2018).

In conclusion, this study shows a significant increase in antibody responses in individuals coexposed to or coinfected with *P. falciparum* and helminths in comparison with individuals infected and/or exposed to only one of these parasites. Although we cannot rule out a possible synergistic effect of these parasites at the immunological level, our results suggest that the increase in the antibody responses is due to a more permissive immune environment to infection in the host. Additional studies are needed for a better understanding of the mechanisms involved during coinfections at the immunological level, but also that lead to coinfections in first place. We also highlight the importance of taking previous exposure into account, especially in endemic areas where continuous infections imprint and shape the immune system. For future studies, we suggest including the measurement of antigen-specific IgE and IgG subclasses, cytokines and cell populations. Deciphering the implications of coinfections deserves attention since accounting for the real interactions that occur in nature could improve the design of integrated disease control strategies.

## MATERIALS AND METHODS

### Study design

This immunological analysis was done in the context of a community-based, case-cohort study (MARS) that recruited 819 individuals. MARS aimed to design and implement a surveillance platform to identify and characterize genotypically and phenotypically anthelminthic resistance in Manhiça district, Maputo province, Southern Mozambique. MARS included inhabitants aged 5 years or above, censed in the Manhiça Health Research Centre (CISM) Demographic Surveillance System, who had not taken anthelmintics at any time during the previous 30 days. The locations included were: 3 Fevereiro, Calanga, Ilha Josina, Maluana, Manhiça-sede and Xinavane. The study protocol was approved by the National Bioethics Committee for Health in Mozambique with the approval code number Ref.:517/CNBS/17. An informed consent from individuals who were willing to participate was obtained and, in the case of children, from their parents or guardians. Additionally, participants aged 15 to 17 also gave inform assent.

The final sample size for this analysis was 715. One hundred and three samples were excluded from the analysis because they were either not shipped to Barcelona (Spain) (N= 5) or due to study withdrawal (N= 25), sample insufficiency (N= 4) or incorrect serum elution (N= 69).

### Detection and quantification of *P. falciparum* and helminth infections

Stool, urine, and blood samples were collected from participants for laboratory analyses. Microscopic diagnosis of *N. americanus*, *A. duodenale*, *T. trichiura, A. lumbricoides* and *S. mansoni* was performed at the CISM by search of eggs in two stool samples using duplicate Kato-Katz thick smear technique. *S. stercoralis* was detected in two stool samples using the Telemann technique. *S. haematobium* infection was detected by urine filtration. In addition, molecular diagnosis of STH infections and schistosomiasis was performed at the Leiden University Medical Center (The Netherlands). In brief, the presence of parasite specific DNA was determined in one stool sample per participant by multiplex semi-quantitative real-time Polymerase Chain Reactions (qPCR), using sequences of primers and probes and a general PCR set-up as described before (Kaisar et al., 2017). Individuals were considered to be positive for helminth infection if they had a positive result in at least one of the techniques (microscopy or qPCR) and in at least one of the stool samples.

Capillary blood samples were collected through finger pricking on Whatman 903 filter paper to obtain dried blood spots (DBS). *P. falciparum* infection was determined at the Barcelona Institute for Global Health (ISGlobal) by 18S ribosomal RNA gene detection through qPCR from DBS samples. Briefly, six DBS punches of 3 mm in diameter were used for DNA extraction by the Chelex method (Taylor et al., 2014) along with negative controls (NC) consisting of DBS from non-infected erythrocytes (Banc de Sang i Teixits [BST], Barcelona, Spain) and positive controls (PC) of DBS with *P. falciparum* (3D7 isolate) ring-infected red blood cells diluted in whole blood. Purified DNA templates were amplified following a previously described method (Mayor et al., 2009). A NC with ddH2O instead of DNA template as well as the punch controls were included in each plate. Each specimen was run in duplicate and the standard curve in triplicate for each plate. Parasite density was quantified by Ct interpolation with a standard curve ranging from 18,000 to 1.8 parasites/μL.

### Measurement of antibody responses

#### Antigen panel

A total of 27 antigens, 16 from *P. falciparum* (α-gal, CelTOS, SSP2, LSA1, EXP1, AMA1 FVO, EBA175 reg2 PfF2, MSP1 block 2 MAD20, MSP1_42_ 3D7, MSP2 full length CH150, MSP3 3D7, MSP5, P41, Rh1, Rh5 and PTRAMP) and 11 from helminths (STH [NaGST1, NaAPR1, NaSAA2, AyCp2, TmWAP, Tm16, As16, As37, NIE] and *Schistosoma* spp. [MEA, Sm25]), were selected based on their important role as vaccine candidates and markers of exposure (**Table S6**). They are representative of the different stages of *P. falciparum* lifecycle and of the most common intestinal helminth infections. In addition, glutathione S-transferase (GST) was added to the panel as control for unspecific binding to this tag fused to MSP1 and MSP2 antigens.

As16, As37, AyCp2, NaAPR1, NaGST1, NaSAA2, Tm16 and TmWAP were expressed and purified from yeast *Pichia pastoris* X-33 culture at Baylor University (USA) (**Table S6**). NIE was kindly provided by Thomas Nutman (National Institute of Health and National Institute of Allergy and Infectious Diseases, USA) and Sukwan Handali (Centers for Disease Control and Prevention, USA); MEA and Sm25 were purchased from MyBioSource (USA); α-gal from Dextra Laboratories (UK); and CelTOS, SSP2, LSA1 and EXP1 from Sanaria (USA). AMA1 FVO, MSP1_42_ 3D7 and GST were produced at Walter Reed Army Institute of Research (USA); MSP5 at Monash University (Australia); MSP1 block 2 MAD20 and MSP2 full length CH150 at University of Edinburgh (UK) and EBA175 R2 F2, Rh1, Rh5, PTRAMP and P41 at International Centre for Genetic Engineering and Biotechnology (India).

#### Serum elution from DBS

Antibodies were obtained from DBS samples through an adaptation of a described method (Fonseca et al., 2017). Briefly, serum was eluted from 4 DBS punches, collected as described before, and incubated with 100μl of PBS-BN [filtrated PBS with 1% BSA and 0.05% sodium azide (ref. S8032, MilliporeSigma, St. Louis, USA)] and 0.05% Tween20 (ref. P1379, MilliporeSigma, St. Louis, USA) at 4°C overnight (ON) with gentle mixing (∼600 rpm). Next day, the tubes were centrifuged at 10,000 rpm for 10 min and the supernatant containing the eluted serum was transferred into a new tube, ready for serological measurement. Assuming a haematocrit of 50%, the eluted blood proteins concentration resulted equivalent to a 1:25 plasma dilution. Samples with reddish-brown spots against a pale background were discarded due to incorrect elution. To assure the validity of the punch and the serum elution procedure, a NC of filter paper without DBS and three elution control (EC) samples with known antibody levels (volunteer donors) were included in each elution batch.

#### Luminex

Quantitative suspension array technology (qSAT) assays were used to measure *P. falciparum* and helminth specific-IgG and total IgE as described previously (Vidal et al., 2018). Briefly, the antigens and a capture anti-human IgE (Mouse anti-Human IgE Fc) (ref. ab99834, Abcam PLC, Cambridge, UK) were coupled to magnetic MagPlex® 6.5μm COOH-microspheres (Luminex Corporation, Austin, USA) at the optimal coupling conditions (**Table S7**). Coupled beads were validated (in multiplex in the case of antigen-specific IgG) by measuring antigen-specific IgG and total IgE in serial dilutions of a PC. The antigen-coupled beads were used in multiplex at 1,000 beads per region per sample, while the anti-human IgE-coupled beads were used in singleplex at 3,000 beads per sample.

Coupled beads were incubated with eluted serum samples, eluted controls, a PC curve, and blanks in 96-well μClear® flat bottom plates (ref. 655096, Greiner Bio-One, Frickenhausen, Germany) at 4°C ON in a shaker at 600 rpm. For the antigen-specific IgG quantification we assayed the samples at 1:100 and 1:2000 dilutions, and included a WHO malaria pool was used as PC [First WHO Reference Reagent for Anti-malaria (*P. falciparum*) human serum, NIBSC code: 10/198 (NIBSC, Herts, UK)] at 12 serial dilutions (3-fold) starting at 1:100. For total IgE quantification we assayed the samples at 1:50 dilution and, included a WHO IgE PC [Third WHO International Standard Immunoglobulin E, human serum, NIBSC code: 11/234 (NIBSC, Herts, UK)] at 12 serial dilutions (2-fold) starting at 1/50. After the incubation of coupled beads with samples, beads were washed three times with PBS + 0.05% Tween20, using a manual magnetic washer platform (ref. 40-285, Bio-Rad, Hercules, USA), and the biotinylated secondary anti-human IgG (ref. B1140-1ML, MilliporeSigma, St. Louis, USA)] and the anti-human IgE (ref. A18803, Invitrogen, Waltham, USA) antibodies were added at 1:1250 and 1:100 dilutions, respectively, and incubated for 45 min at RT shaking at 600 rpm. Then, beads were washed three times and streptavidin-R-phycoerythrin (ref. 42250-1ML, MilliporeSigma, St. Louis, USA) was added at 1:1000 dilution and incubated at RT for 30 min at 600 rpm. Finally, beads were washed, resuspended, and at least 50 beads per analyte and sample were acquired in a FlexMap 3D xMAP® instrument (Luminex Corporation, Austin, USA). Crude median fluorescent intensities (MFI) and background fluorescence from blank wells were exported using the xPONENT software v.4.3 (Luminex Corporation, Austin, USA).

### Determination of seropositivity

Gaussian Mixture Models were used to estimate a seropositivity threshold for each antigen-specific IgG and total IgE response and classify the samples into positive and negative by serology. The models were estimated by the Expectation-Maximization (EM) algorithm. For each antigen, two models were fitted assuming either equal or unequal variances, and each of those two models were built with two clusters for classification. The two clusters of classification are meant to represent the distribution of seropositive and seronegative samples. For each antigen, classification of samples was done using the model showing the optimal Bayesian Information Criterion (BIC). Briefly, the EM algorithm is an unsupervised clustering algorithm that works in two steps. The first step is the Expectation step, where the data are assigned to the closest centroid of the two clusters. The second step is the Maximization step, where the centroids of the clusters are recalculated with the newly assigned data. These two steps are repeated until the model converges. The models were fitted using the R CRAN package mclust (Scrucca et al., 2016).

### Definition of study groups

Study participants were classified into study groups based on diagnosis of current infection (defined by microscopy and/or qPCR) and prior exposure (defined by seropositivity to any of the antigens for each type of parasite). The resulting groups were: a) no exposure/infection (MAL- HEL-), which included people that have not been exposed to or currently infected with *P. falciparum* nor helminths; b) single exposure/infections (MAL- HEL+ or MAL+ HEL-), which included people that have been exposed to and/or currently infected with only one type of parasite; and c) coexposure/coinfection (MAL+ HEL+), which included people that have been exposed to and/or currently infected with both types of parasites.

### Definition of study variables

In this study we included demographic variables such as sex, age, age group (children [5-15 years old] or adults [≥15 years old]) and percentage of infection by location. The latter was calculated based on the *P. falciparum* or helminth percentage of infection in each location of the study. Then, participants were classified into *P. falciparum* or helminth “high” and “low” groups of locations defined according to higher or lower percentage of *P. falciparum* or helminths compared to the overall median by type of parasite (**Table S4**). Additionally, we also included socioeconomic (socioeconomic score), water (accessibility to piped water, payment for water) and sanitation (ownership of latrine) variables (Grau-Pujol et al., 2021, in preparation). Finally, we calculated the magnitude and breadth of response to *P. falciparum* and helminth antigens as previously described (Tran et al., 2019). The magnitude of response corresponds to the sum of all specific IgG levels (MFI) to the different antigens belonging to *P. falciparum* or helminths, and the breadth of response is defined as the number of antigens with seropositive responses.

### Statistical analysis

To determine the seropositivity and perform the statistical analysis, normalized and dilution corrected data (see Supplementary Material) were log_10_-transformed. The Shapiro-Wilk test was used to check the normality of the data and since most of the data did not follow a normal distribution, non-parametric tests were used. Then, we explored differences in the distribution of study variables across study groups. For continuous variables, the median and first and third quartiles were calculated and differences in the median between more than two groups were assessed by Kruskall-Wallis test followed by pair-wise comparisons by Wilcoxon rank sum test and Benjamini-Hochberg (BH) adjustment for multiple testing. For categorical variables, differences in proportions between groups were calculated by Chi-square test or Fisher’s exact test when applicable. This was automatically determined by the R CRAN package compareGroups (Subirana et al., 2014). For comparisons between two groups we applied the Wilcoxon rank sum test and, in the case of antigen-specific IgG responses, p-values were adjusted for multiple testing by the BH. Correlations between variables were assessed with the Spearman’s rank correlation coefficient ρ (rho). For Spearman’s test, p-values were computed via the asymptotic *t* approximation.

For regression models, assumptions on the data were checked by exploratory plots. For linear regression models, we checked linearity of the data, normality of residuals, homogeneity of residuals variance and independence of residuals error terms. For logistic regression models, we checked the linearity between the logit of the outcome and each predictor variable, the presence of influential values and multicollinearity among the predictor variables. All the assumptions were met without the presence of extreme patterns, except for normality of the residuals in most cases.

The first linear regression models were performed to assess the association between antibody responses (including antigen-specific IgG levels, total IgE levels, magnitude of response and breadth of response) as outcome variables and sex, log_10_ age, percentage of infected individuals by location, socioeconomic score, accessibility to piped water, payment for water and ownership of latrine in univariable models. The second linear regression models were multivariable models fitted to assess the association between antibody responses as outcome variables and three predictor variables: a) the exposure/infection status as categorical variable (coexposed/coinfected or single exposed/infected), b) log_10_ total IgE levels as continuous variable or c) the magnitude of response to *P. falciparum* or helminth antigens as continuous variable. We evaluated the association of these three predictor variables independently by building three different base models. To select the adjusting covariables, we first selected those variables significantly associated with each block of antigens by univariable models. Then, we explored all possible combinations of the predictor covariables with the base models and chose per block of antigens those combinations that provided the simplest models (based on BIC) without a significant compromise of the Akaike information criterion (AIC) and adjusted r-square. The third linear regression models explored the association between *P. falciparum* exposure/infection (predictor variable) and the number of helminth species (outcome variable) adjusted by the corresponding covariables selected as explained above. The fourth linear regression models were performed to assess the association between parasite burden as outcome variable and antibody responses as predictor variables adjusted by age. We adjusted for multiple comparisons by BH, where the different antibody responses penalized.

The logistic regression models evaluated the association of exposure/infection with either *P. falciparum* or helminths (predictor variable) with being exposed/infected with the counterpart parasite (outcome variable). For *P. falciparum*, we also evaluated by logistic regression the association of the number of helminth species (predictor variable) with exposure/infection with *P. falciparum* (outcome variable). The selection of adjusting covariables was done based on significant associations by univariable models.

The resulting betas and 95% confidence intervals (CI) obtained in each linear regression model were transformed when appropriate for interpretation if the outcome (log-linear model), predictor (linear-log model), or both (log-log model) variables were log_10_-transormed. For log-log models, the transformed values (%) were calculated with the formula ((10^(beta*log_10_(1.1)))-1)*100. This represents the effect (in percentage) on the outcome variable of a 10% increase in the corresponding predictor variable. For log-linear models, the transformed values (%) were calculated with the formula ((10^beta)-1)*100. This represents the effect (in percentage) on the outcome variable of an increase in one unit of the predictor variable, or with respect to the reference category in cases of categorical variables. Finally, for linear-log models, the transformed value was calculated with the formula beta*log_10_(1.1). This represents the additive effect on the outcome variable of a 10% increase in the corresponding predictor variable.

Adjusted and raw (if not adjusted for multiple testing) p-values of ≤0.05 were considered statistically significant. All data processing and statistical analyses were performed using the statistical software R version 4.0.3 and R Studio Version 1.1.463. Other R CRAN packages not mentioned above but used to manage data, generate tables and plots were tidyverse (Wickham et al., 2019), ggiraphExtra (Moon, 2020), ggpubr (Kassambara, 2020), ggbeeswarm (Clarke and Sherrill-Mix, 2017).

## ACKNOWLEDGEMENTS

We are grateful to the volunteers and their families and to all field workers and lab technicians that participated in this study. We would also like to thank CISM Demography and Social Sciences department for their indispensable assistance during field work. We are thankful to Javier Gandasegui for parasitology lab training. Special thanks to the team members from ISGlobal, specifically Laura Puyol for the organization and coordination of shipment of samples, Diana Barrios and Alfons Jiménez for the technical support in the lab, Miquel Vázquez-Santiago and Llorenç Quintó for the statistical support and Chenjerai Jairoce for the support in assay optimization. For antigen procurement, we also thank Thomas Nutman (NIH/NIAID, USA) and Sukwan Handali (CDC, USA). On qPCR analysis, we particularly would like to thank Eric Brienen (LUMC, The Netherlands).

## SUPPLEMENTARY TABLES

**Table S1.** Percentage of exposure, infection or both by parasite species and groups.

|  | Total | MAL- HEL-<br>(N=21) | MAL- HEL+<br>(N=43) | MAL+ HEL-<br>(N=67) | MAL+ HEL+<br>(N=584) | p-value |
| --- | --- | --- | --- | --- | --- | --- |
| **Present infection** |  |  |  |  |  |  |
| Hookworm | 201 (28.11%) | 0 (0.00%) | 17 (39.5%) | 0 (0.00%) | 184 (31.5%) | 0.358 |
| *T. trichiura* | 103 (14.41%) | 0 (0.00%) | 14 (32.6%) | 0 (0.00%) | 89 (15.2%) | **0.008** |
| *A. lumbricoides* | 53 (7.41%) | 0 (0.00%) | 5 (11.6%) | 0 (0.00%) | 48 (8.22%) | 0.399 |
| *S. stercoralis* | 74 (10.35%) | 0 (0.00%) | 1 (2.33%) | 0 (0.00%) | 73 (12.5%) | 0.158 |
| *Schistosoma spp.* | 86 (12.03%) | 0 (0.00%) | 1 (2.33%) | 0 (0.00%) | 85 (14.6%) | 0.097 |
| Any helminth | 383 (53.57%) | 0 (0.00%) | 31 (72.1%) | 0 (0.00%) | 352 (60.3%) | 0.170 |
| *P. falciparum* | 71 (9.93%) | 0 (0.00%) | 0 (0.00%) | 5 (7.46%) | 66 (11.3%) | 0.455 |
| **Positive serology** |  |  |  |  |  |  |
| Hookworm | 497 (69.51%) | 0 (0.00%) | 17 (39.5%) | 0 (0.00%) | 480 (82.2%) | **≤0.001** |
| *T. trichiura* | 377 (52.73%) | 0 (0.00%) | 7 (16.3%) | 0 (0.00%) | 370 (63.4%) | **≤0.001** |
| *A. lumbricoides* | 148 (20.7%) | 0 (0.00%) | 0 (0.00%) | 0 (0.00%) | 148 (25.3%) | **≤0.001** |
| *S. stercoralis* | 195 (27.27%) | 0 (0.00%) | 1 (2.33%) | 0 (0.00%) | 194 (33.2%) | **≤0.001** |
| *Schistosoma spp.* | 314 (43.92%) | 0 (0.00%) | 4 (9.30%) | 0 (0.00%) | 310 (53.1%) | **≤0.001** |
| Any helminth | 566 (79.16%) | 0 (0.00%) | 22 (51.2%) | 0 (0.00%) | 544 (93.2%) | **≤0.001** |
| *P. falciparum* | 651 (91.05%) | 0 (0.00%) | 0 (0.00%) | 67(100%) | 584 (100%) | NA |
| **Exposure/infection** |  |  |  |  |  |  |
| Hookworm | 540 (75.52%) | 0 (0.00%) | 31 (72.1%) | 0 (0.00%) | 509 (87.2%) | **0.011** |
| *T. trichiura* | 426 (59.58%) | 0 (0.00%) | 20 (46.5%) | 0 (0.00%) | 406 (69.5%) | **0.003** |
| *A. lumbricoides* | 190 (26.57%) | 0 (0.00%) | 5 (11.6%) | 0 (0.00%) | 185 (31.7%) | **0.012** |
| *S. stercoralis* | 210 (29.37%) | 0 (0.00%) | 2 (4.65%) | 0 (0.00%) | 208 (35.6%) | **≤0.001** |
| *Schistosoma spp.* | 339 (47.41%) | 0 (0.00%) | 5 (11.6%) | 0 (0.00%) | 334 (57.2%) | **≤0.001** |
| Any helminth | 627 (87.69%) | 0 (0.00%) | 43 (100%) | 0 (0.00%) | 584 (100%) | NA |
| *P. falciparum* | 651 (91.05%) | 0 (0.00%) | 0 (0.00%) | 67(100%) | 584 (100%) | NA |

**Table S2.** Percentage of single or multiple helminth species exposure, infection or both by helminth positive groups.

| Helminth species | Total | MAL- HEL+ | MAL+ HEL+ | p-value |
| --- | --- | --- | --- | --- |
| Present infection |  |  |  |  |
| Single | 271 (70.76%) | 25 (80.6%) | 246 (69.9%) | 0.291 |
| Multiple | 112 (29.24%) | 6 (19.4%) | 106 (30.1%) |  |
| Total | 383 (100%) | 31 (100%) | 352 (100%) |  |
| Positive serology |  |  |  |  |
| Single | 129 (22.79%) | 16 (72.7%) | 113 (20.8%) | ≤0.001 |
| Multiple | 437 (77.21%) | 6 (27.3%) | 431 (79.2%) |  |
| Total | 566 (100%) | 22 (100%) | 544 (100%) |  |
| Exposure/Infection |  |  |  |  |
| Single | 136 (21.69%) | 27 (62.8%) | 109 (18.7%) | ≤0.001 |
| Multiple | 491 (78.31%) | 16 (37.2%) | 475 (81.3%) |  |
| Total | 627 (100%) | 41 (100%) | 584 (100%) |  |

**Table S3.** Comparison of results by diagnostic methods and serology.

| Species | Diagnostic method | Diagnostic method result | Serology |  |
| --- | --- | --- | --- | --- |
|  |  |  | - | + |
| *P. falciparum* | qPCR | - | 64 | 580 |
|  |  | + | 0 | 71 |
| *A. lumbricoides* | Microscopy | - | 541 | 141 |
|  |  | + | 26 | 7 |
|  | qPCR | - | 517 | 126 |
|  |  | + | 36 | 11 |
|  | Microscopy and/or PCR | - | 525 | 137 |
|  |  | + | 42 | 11 |
| *S. stercoralis* | Microscopy | - | 519 | 188 |
|  |  | + | 1 | 7 |
|  | qPCR | - | 489 | 130 |
|  |  | + | 14 | 57 |
|  | Microscopy and/or qPCR | - | 505 | 136 |
|  |  | + | 15 | 59 |
| Hookworm | Microscopy | - | 196 | 401 |
|  |  | + | 22 | 96 |
|  | qPCR | - | 173 | 333 |
|  |  | + | 38 | 146 |
|  | Microscopy and/or PCR | - | 175 | 339 |
|  |  | + | 43 | 158 |
| *T. trichiura* | Microscopy | - | 318 | 342 |
|  |  | + | 20 | 35 |
|  | qPCR | - | 283 | 313 |
|  |  | + | 45 | 49 |
|  | Microscopy and/or qPCR | - | 289 | 323 |
|  |  | + | 49 | 54 |
| *Schistosoma spp.* | Microscopy | - | 393 | 290 |
|  |  | + | 8 | 24 |
|  | qPCR | - | 371 | 248 |
|  |  | + | 19 | 52 |
|  | Microscopy and/or qPCR | - | 376 | 253 |
|  |  | + | 25 | 61 |

Gain in positive results by qPCR and/or microscopy in dark grey. Gain in positive results by serology in light grey.

**Table S4.** Classification of locations based on percentage of individuals infected with Plasmodium falciparum or helminths.

| Location | MAL- | MAL+ | Total | MAL + (%) | Median % of infection | Q1 | Q3 | Classification with respect to the median % of infection |
| --- | --- | --- | --- | --- | --- | --- | --- | --- |
| 3 Fevereiro | 101 | 15 | 116 | 12.931 | 13.203 | 6.823 | 21.569 | Low |
| Calanga | 122 | 19 | 141 | 13.475 |  |  |  | High |
| Ilha Josina | 22 | 8 | 30 | 26.667 |  |  |  | High |
| Maluana | 179 | 9 | 188 | 4.787 |  |  |  | Low |
| Manhiça-sede | 180 | 9 | 189 | 4.762 |  |  |  | Low |
| Xinavane | 40 | 11 | 51 | 21.569 |  |  |  | High |
|  | **HEL-** | **HEL+** |  | **HEL + (%)** |  |  |  |  |
| 3 Fevereiro | 51 | 65 | 116 | 56.034 | 48.814 | 45.331 | 51.596 | High |
| Calanga | 36 | 105 | 141 | 74.468 |  |  |  | High |
| Ilha Josina | 24 | 6 | 30 | 20 |  |  |  | Low |
| Maluana | 91 | 97 | 188 | 51.596 |  |  |  | High |
| Manhiça-sede | 102 | 87 | 189 | 46.032 |  |  |  | Low |
| Xinavane | 28 | 23 | 51 | 45.098 |  |  |  | Low |

Malaria (MAL) and helminth (HEL) percentage of infection in each location of the study were calculated and these were classified into “high” and “low” prevalence locations based on higher or lower percentage of infected individuals compared to the median percentage of infections from all locations by type of parasite.

**Table S5.**
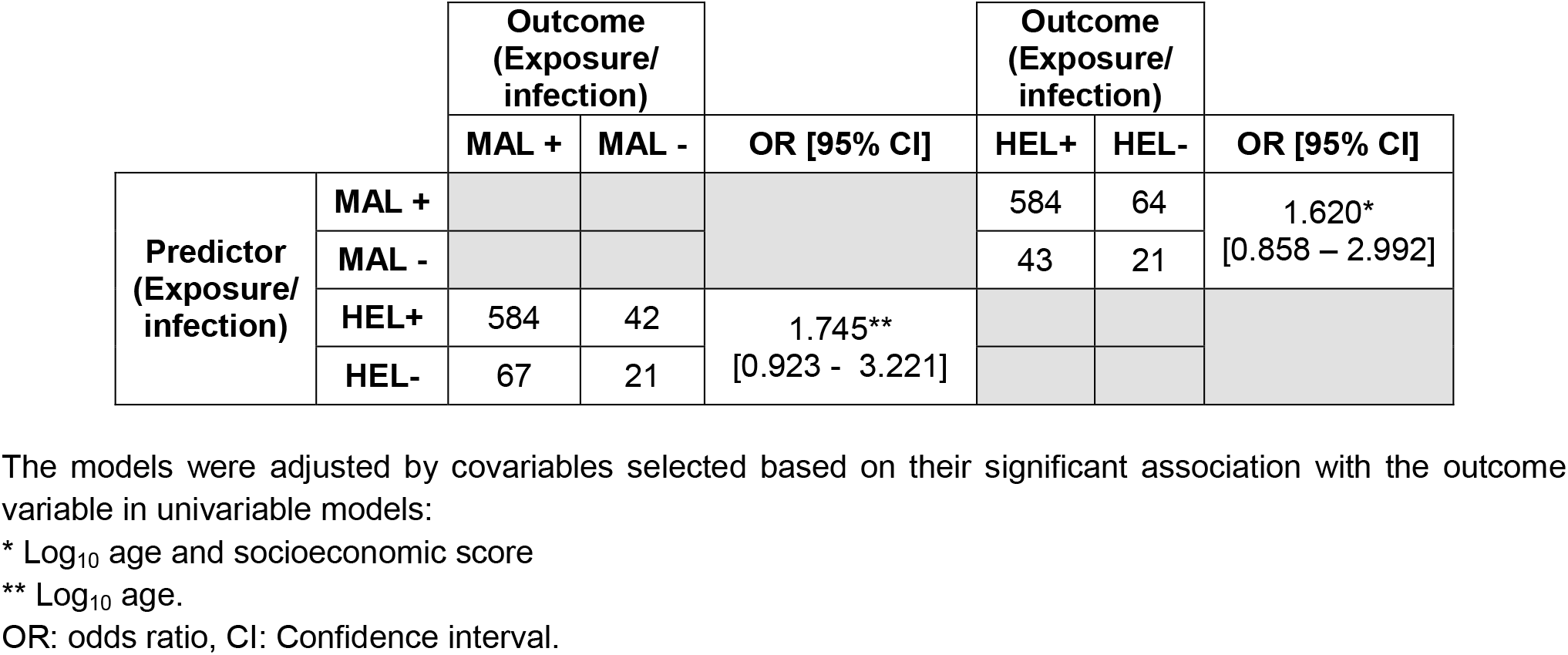
Risk of exposure to or infection with *Plasmodium falciparum* or helminths.

## SUPPLEMENTARY FIGURES

**Figure S1.**
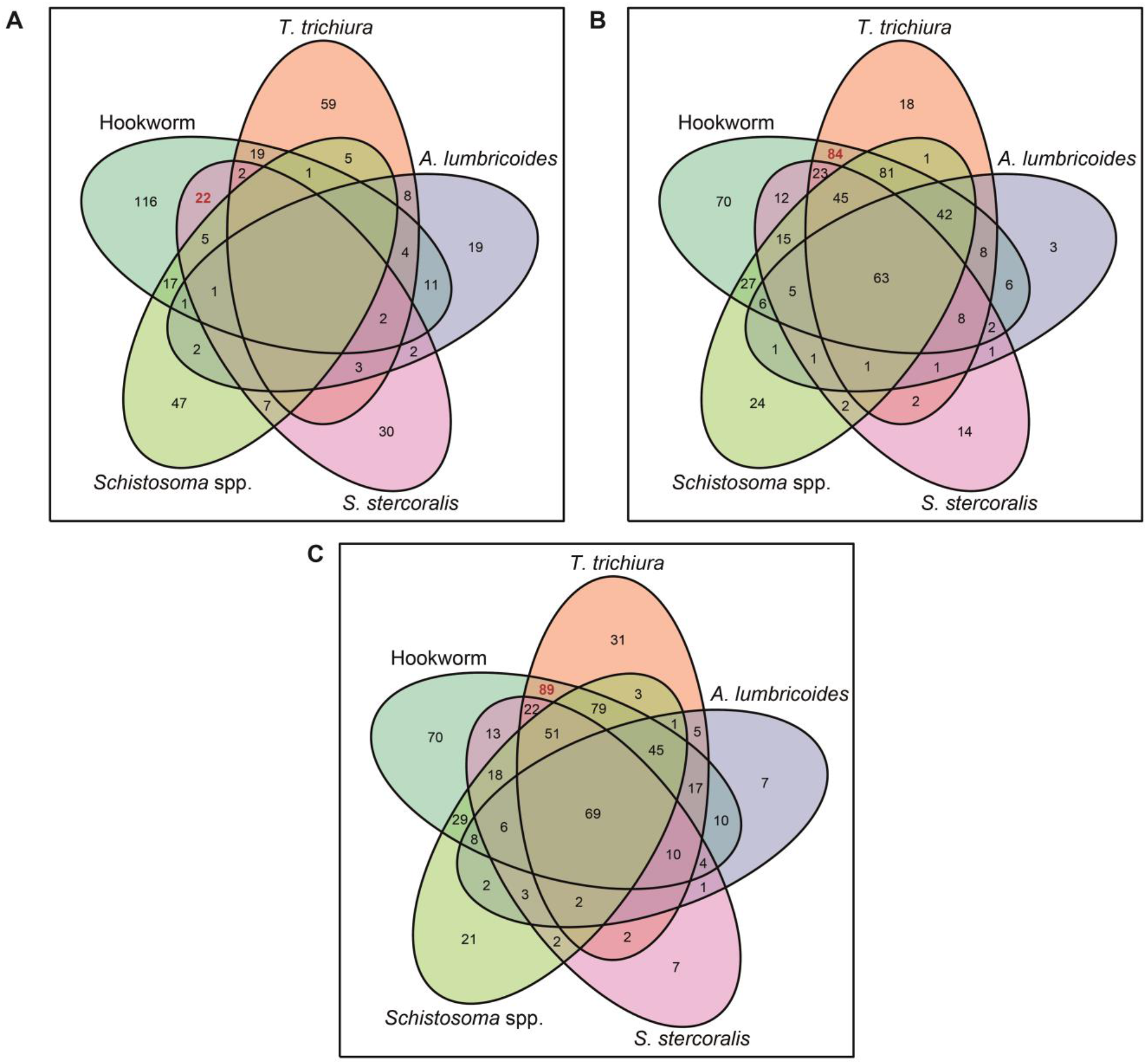
Venn diagrams showing the combinations of (A) coinfections, (B) coexposure and (C) coexposure/coinfection of helminth species. The most common combination of species in each classification is highlighted in red.

**Figure S2.**
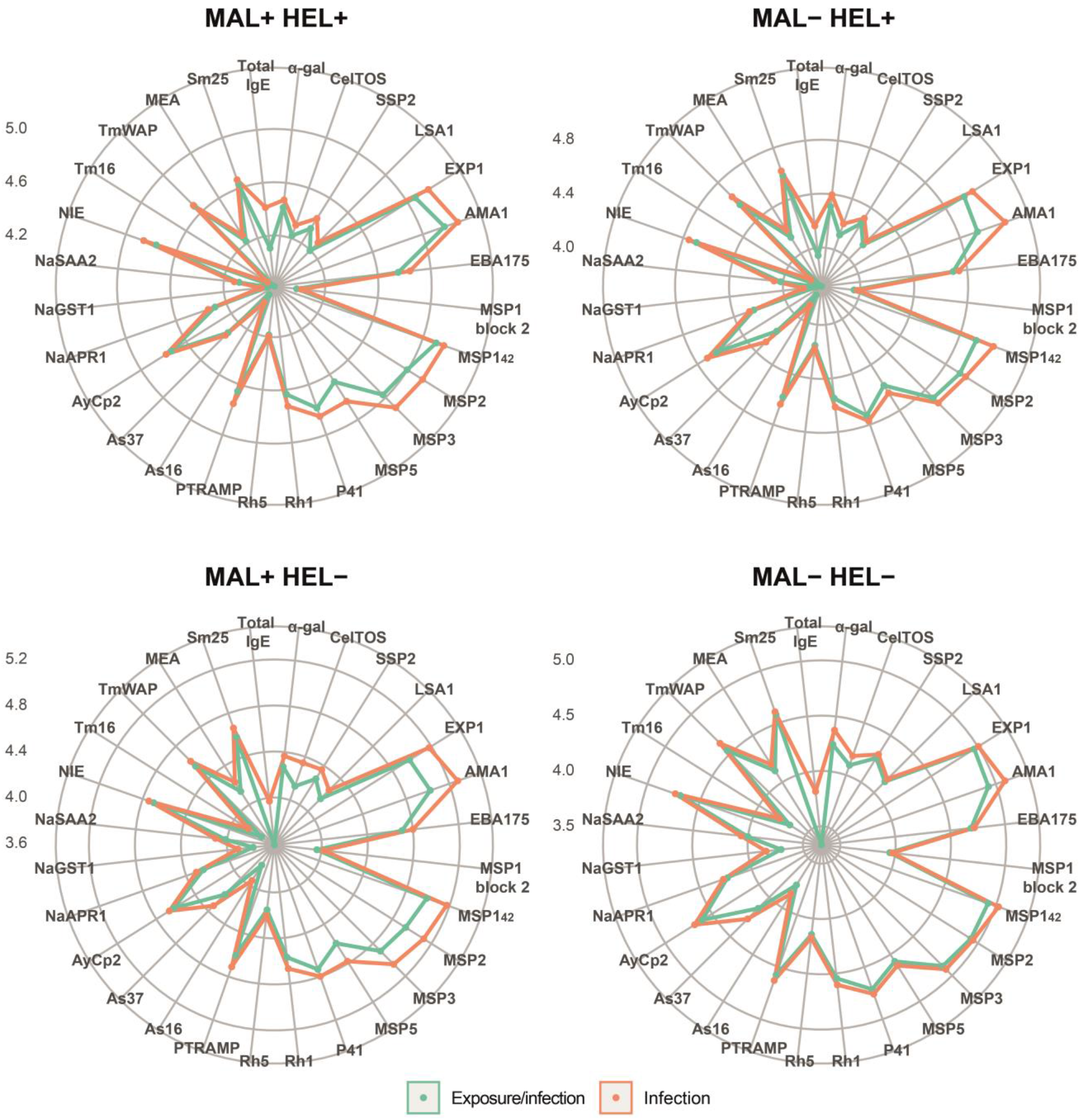
Comparison of antibody responses between groups classified by exposure/infection and only infection. In the vertices, the antibody responses are grouped by specific IgG to *P. falciparum* antigens (α-gal, CelTOS, SSP2, LSA1, EXP1, AMA1, EBA175, MSP1 block 2, MSP1_42_, MSP2, MSP3, MSP5, P41, Rh5, PTRAMP), helminth antigens (As16 and As37 [*Ascaris lumbricoides*], AyCp2, NaAPR1, NaGST1, NaSAA2 [Hookworm], NIE [*Strongyloides stercoralis*], Tm16 and TmWAP [*Trichuris trichiura*], MEA and Sm25 [*Schistosoma* spp.] or total IgE. The colored lines represent the median of antibody levels expressed as log_10_-transformed median fluorescence levels (MFI). Comparisons were made within study groups: no exposure/infection (green) or no infection (orange) (MAL- HEL-), only exposure/infection (green) or only infection (orange) with helminths (MAL- HEL+), only exposure/infection (green) or only infection (orange) with *P. falciparum* (MAL+ HEL-) and coexposure/coinfection (green) or coinfection (orange) with *P. falciparum* and helminths (MAL+ HEL+).

**Figure S3.**
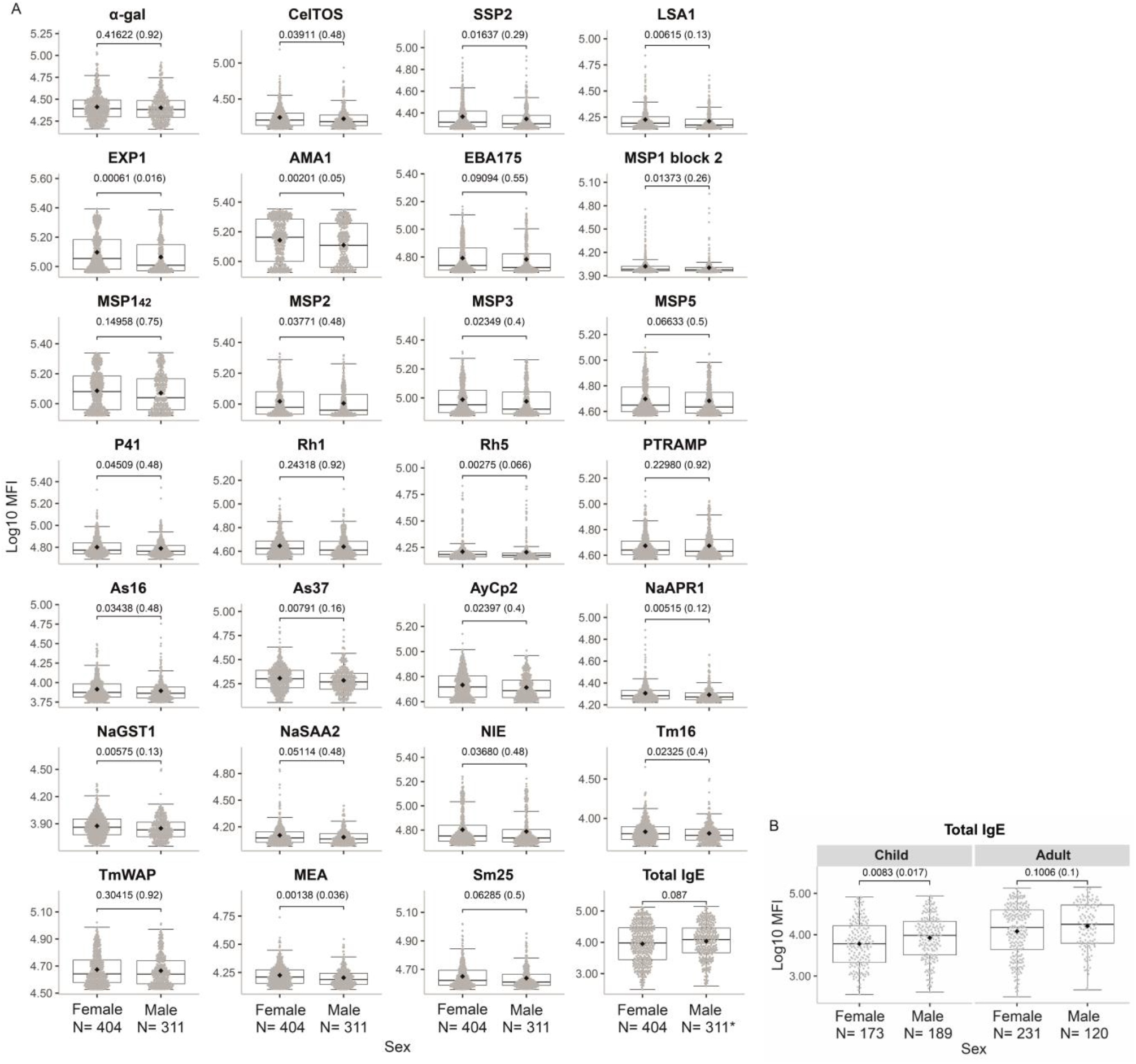
Comparison of antibody responses by sex. **(A)** Antibody levels expressed as the log_10_- transformed median fluorescence (MFI) of specific IgG to *P. falciparum* antigens (α-gal, CelTOS, SSP2, LSA1, EXP1, AMA1, EBA175, MSP1 block 2, MSP1_42_, MSP2, MSP3, MSP5, P41, Rh5, PTRAMP), helminth antigens (As16 and As37 [*Ascaris lumbricoides*], AyCp2, NaAPR1, NaGST1, NaSAA2 [Hookworm], NIE [*Strongyloides stercoralis*], Tm16 and TmWAP [*Trichuris trichiura*], MEA and Sm25 [*Schistosoma* spp.] or total IgE. **(B)** Log_10_-transformed MFI levels of total IgE stratified by age. The boxplots represent the median (bold line), the mean (black diamond), the 1^st^ and 3^rd^ quartiles (box) and the largest and smallest values within 1.5 times the inter-quartile range (whiskers). Data beyond the end of the whiskers are outliers. Statistical comparison between groups was performed by Wilcoxon rank sum test and the exact p-values are shown. Adjusted p-values by the Benjamini-Hochberg approach are shown in brackets Wilcoxon rank sum test. *N= 309.

**Figure S4.**
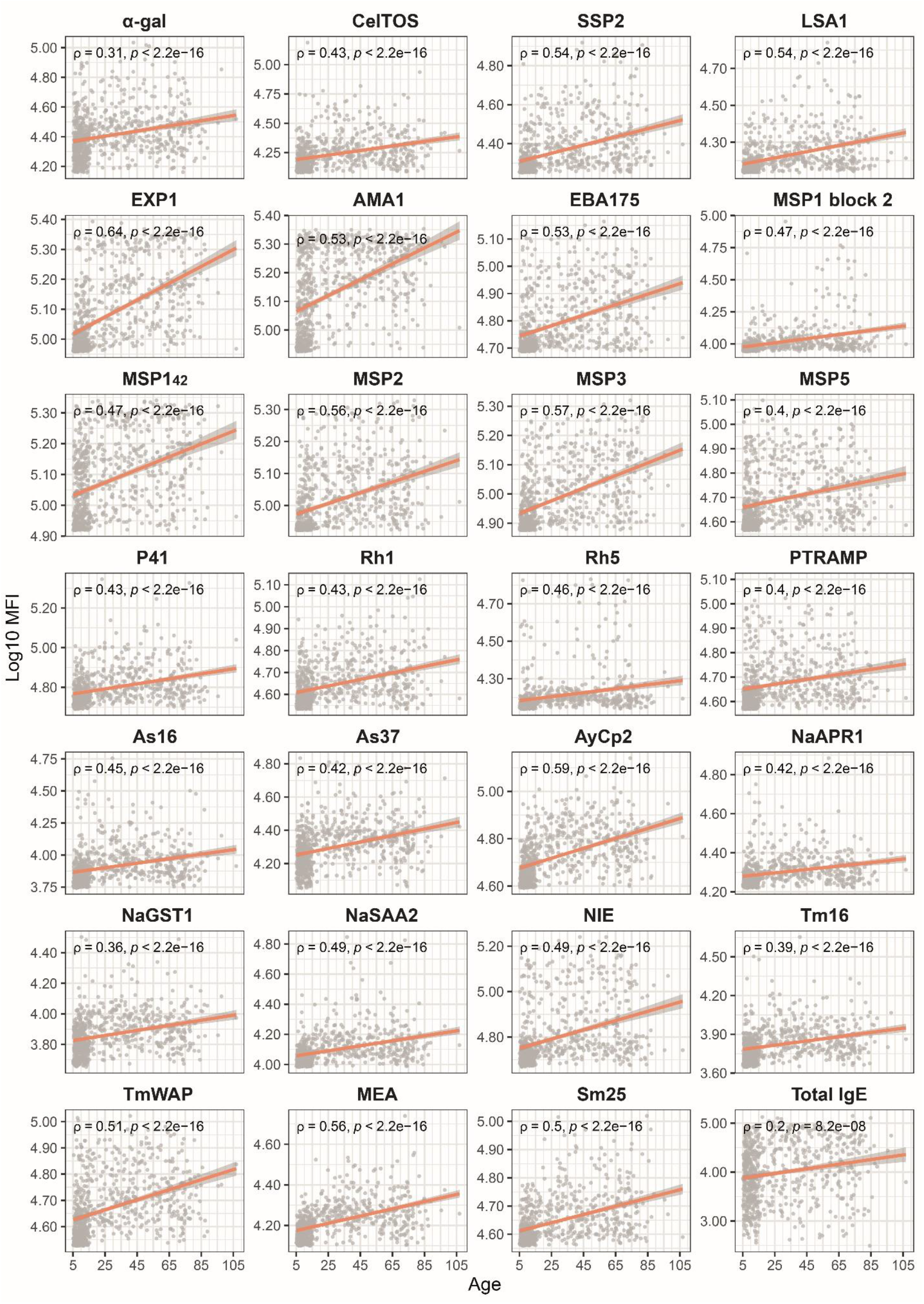
Antibody levels correlation with age. Antibody levels expressed as the log_10_-transformed median fluorescence (MFI) of specific IgG to *P. falciparum* antigens (α-gal, CelTOS, SSP2, LSA1, EXP1, AMA1, EBA175, MSP1 block 2, MSP1_42_, MSP2, MSP3, MSP5, P41, Rh5, PTRAMP), helminth antigens (As16 and As37 [*Ascaris lumbricoides*], AyCp2, NaAPR1, NaGST1, NaSAA2 [Hookworm], NIE [*Strongyloides stercoralis*], Tm16 and TmWAP [*Trichuris trichiura*], MEA and Sm25 [*Schistosoma* spp.] or total IgE. Age is expressed in years. ρ (rho) and p-values were calculated by Spearman, shaded areas represent 0.95 confidence intervals.

**Figure S5.**
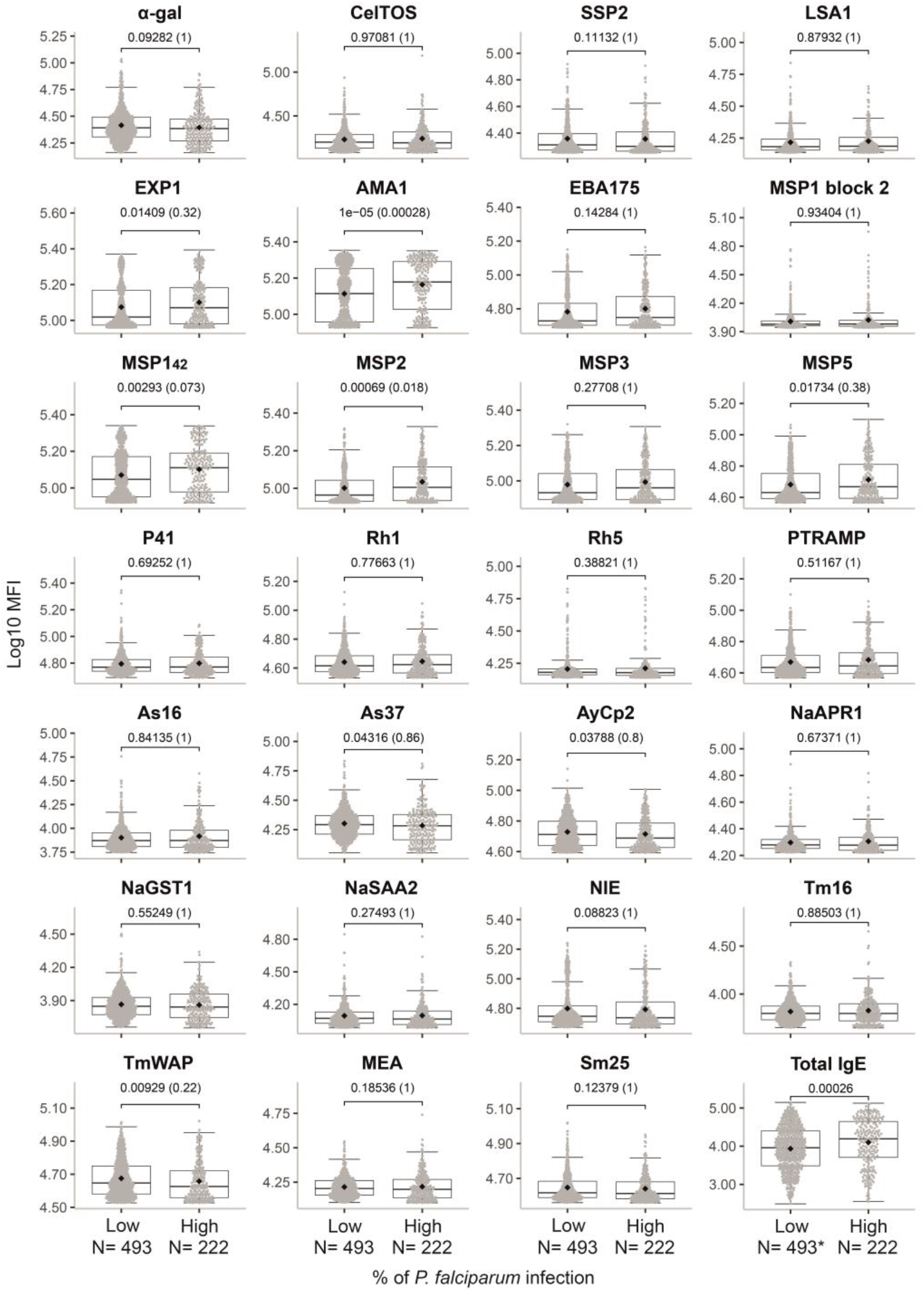
Comparison of antibody responses by percentage of *P. falciparum* infection by location. Antibody levels expressed as the log_10_-transformed median fluorescence (MFI) of specific IgG to *P. falciparum* antigens (α-gal, CelTOS, SSP2, LSA1, EXP1, AMA1, EBA175, MSP1 block 2, MSP1_42_, MSP2, MSP3, MSP5, P41, Rh5, PTRAMP), helminth antigens (As16 and As37 [*Ascaris lumbricoides*], AyCp2, NaAPR1, NaGST1, NaSAA2 [Hookworm], NIE [*Strongyloides stercoralis*], Tm16 and TmWAP [*Trichuris trichiura*], MEA and Sm25 [*Schistosoma* spp.] or total IgE. In the X-axis, locations were classified into “high” and “low” prevalence based on whether the percentage of *P. falciparum* infection was lower or higher than the median percentage of *P. falciparum* infections in all locations. The boxplots represent the median (bold line), the mean (black diamond), the 1^st^ and 3^rd^ quartiles (box) and the largest and smallest values within 1.5 times the inter-quartile range (whiskers). Data beyond the end of the whiskers are outliers. Statistical comparison between groups was performed by Wilcoxon rank sum test and the exact p-values are shown. Adjusted p-values by the Benjamini-Hochberg approach are shown in brackets. *N= 491.

**Figure S6.**
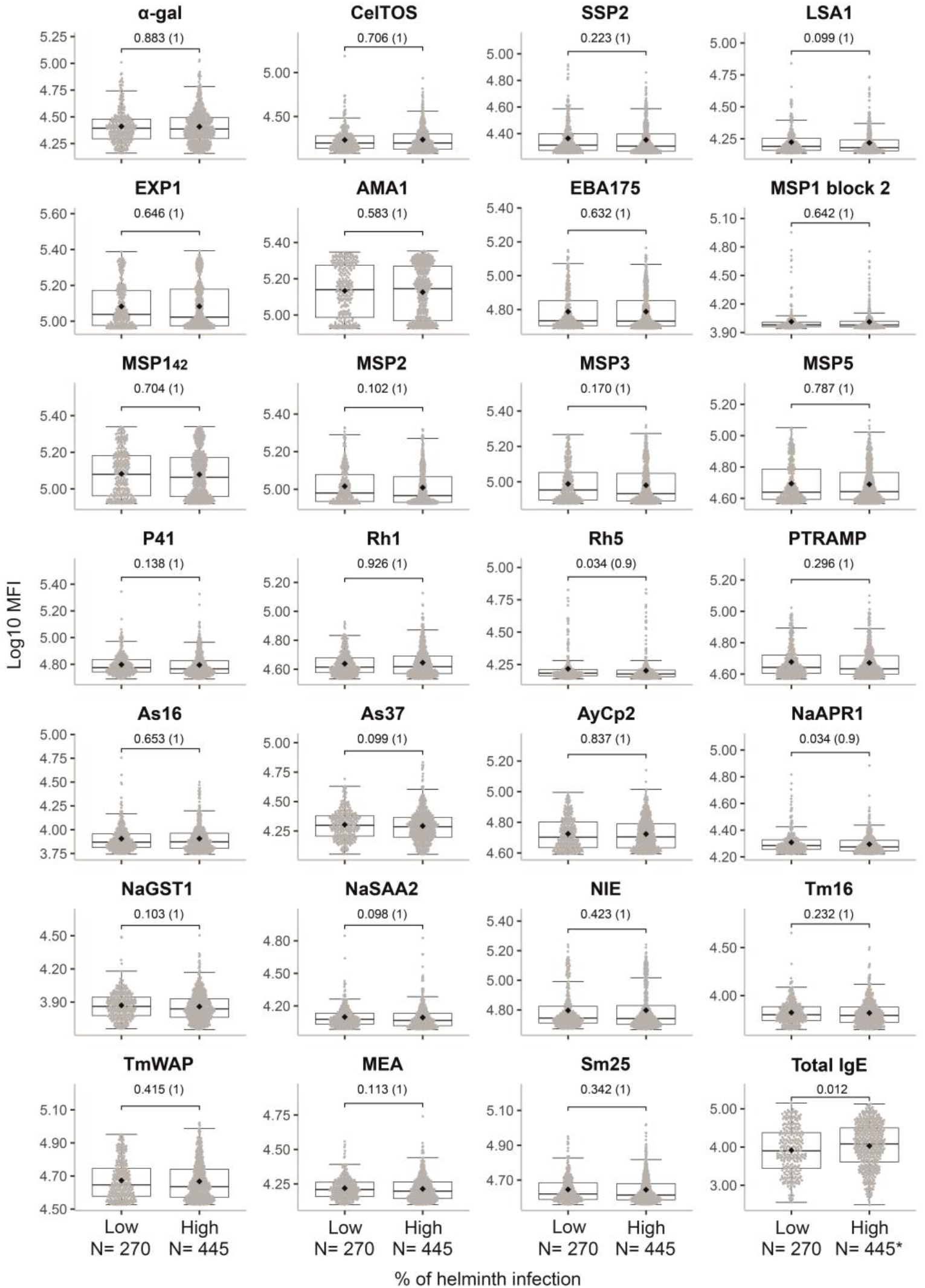
Comparison of antibody responses by percentage of helminth infection by location. Antibody levels expressed as the log_10_-transformed median fluorescence (MFI) of specific IgG to *P. falciparum* antigens (α-gal, CelTOS, SSP2, LSA1, EXP1, AMA1, EBA175, MSP1 block 2, MSP1_42_, MSP2, MSP3, MSP5, P41, Rh5, PTRAMP), helminth antigens (As16 and As37 [*Ascaris lumbricoides*], AyCp2, NaAPR1, NaGST1, NaSAA2 [Hookworm], NIE [*Strongyloides stercoralis*], Tm16 and TmWAP [*Trichuris trichiura*], MEA and Sm25 [*Schistosoma* spp.] or total IgE. . In the X-axis, locations were classified into “high” and “low” prevalence based on whether the percentage of helminth infection was lower or higher than the median percentage of helminth infections in all locations. The boxplots represent the median (bold line), the mean (black diamond), the 1^st^ and 3^rd^ quartiles (box) and the largest and smallest values within 1.5 times the inter-quartile range (whiskers). Data beyond the end of the whiskers are outliers. Statistical comparison between groups was performed by Wilcoxon rank sum test and the exact p-values are shown. Adjusted p-values by the Benjamini-Hochberg approach are shown in brackets. *N= 443.

**Figure S7.**
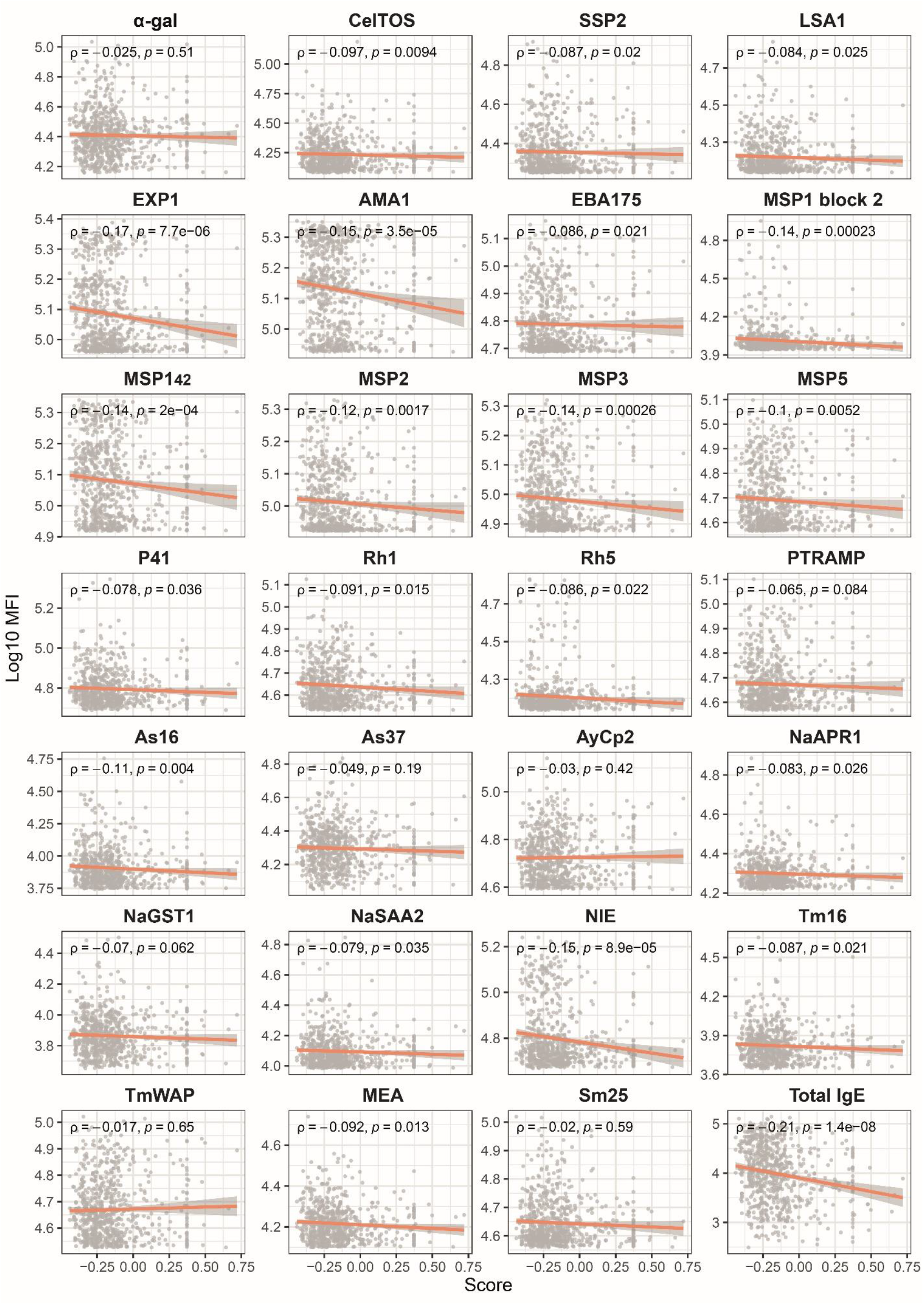
Antibody levels correlation with socioeconomic score. Antibody levels expressed as the log_10_- transformed median fluorescence (MFI) of specific IgG to *P. falciparum* antigens (α-gal, CelTOS, SSP2, LSA1, EXP1, AMA1, EBA175, MSP1 block 2, MSP1_42_, MSP2, MSP3, MSP5, P41, Rh5, PTRAMP), helminth antigens (As16 and As37 [*Ascaris lumbricoides*], AyCp2, NaAPR1, NaGST1, NaSAA2 [Hookworm], NIE [*Strongyloides stercoralis*], Tm16 and TmWAP [*Trichuris trichiura*], MEA and Sm25 [*Schistosoma* spp.] or total IgE. ρ (rho) and p-values were calculated by Spearman, shaded areas represent 0.95 confidence intervals.

**Figure S8.**
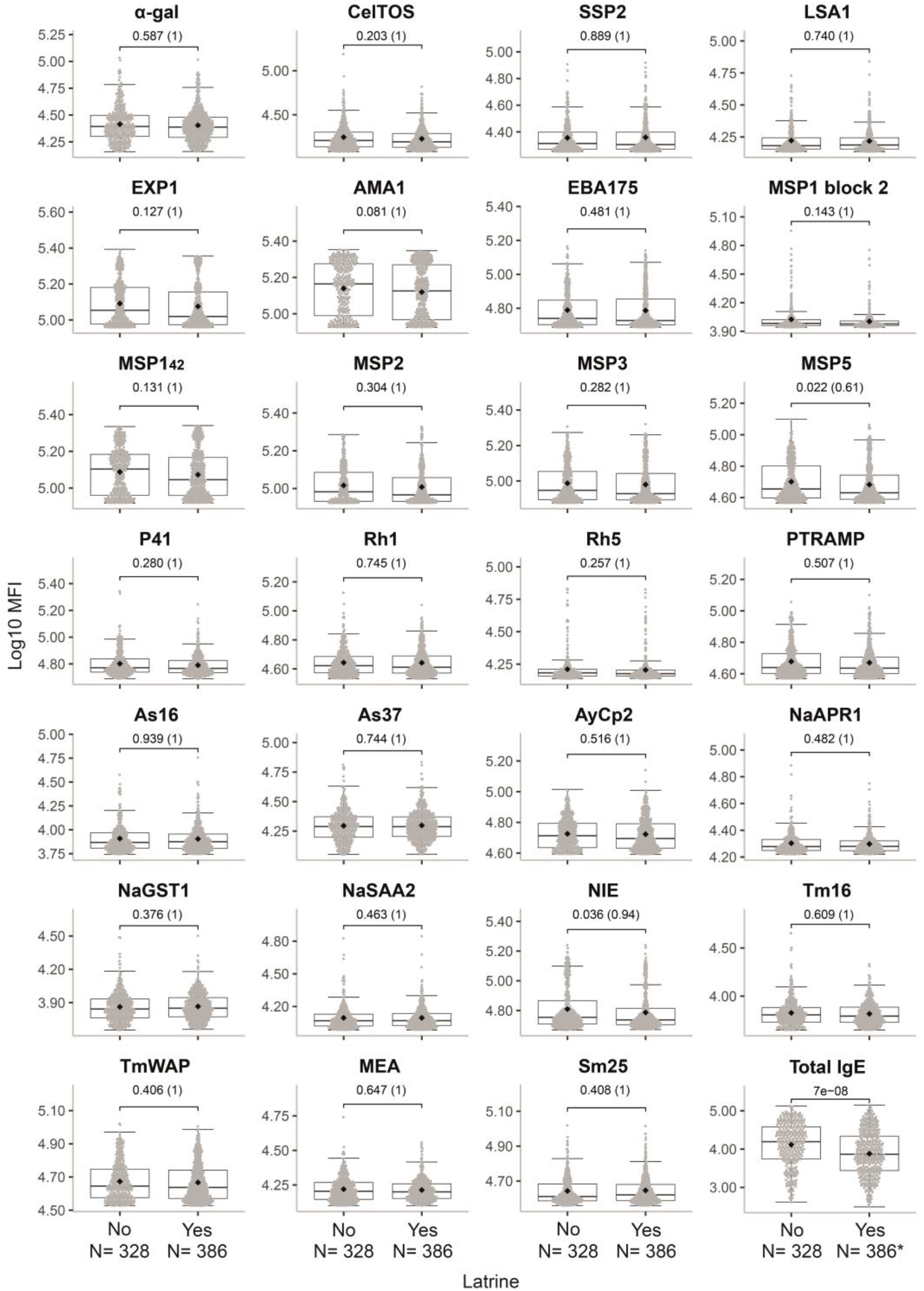
Comparison of antibody responses by ownership of latrine. Antibody levels expressed as the log_10_-transformed median fluorescence (MFI) of specific IgG to *P. falciparum* antigens (α-gal, CelTOS, SSP2, LSA1, EXP1, AMA1, EBA175, MSP1 block 2, MSP1_42_, MSP2, MSP3, MSP5, P41, Rh5, PTRAMP), helminth antigens (As16 and As37 [*Ascaris lumbricoides*], AyCp2, NaAPR1, NaGST1, NaSAA2 [Hookworm], NIE [*Strongyloides stercoralis*], Tm16 and TmWAP [*Trichuris trichiura*], MEA and Sm25 [*Schistosoma* spp.] or total IgE. The boxplots represent the median (bold line), the mean (black diamond), the 1^st^ and 3^rd^ quartiles (box) and the largest and smallest values within 1.5 times the inter-quartile range (whiskers). Data beyond the end of the whiskers are outliers. Statistical comparison between groups was performed by Wilcoxon rank sum test and the exact p-values are shown. Adjusted p-values by the Benjamini-Hochberg approach are shown in brackets. *N= 384.

**Figure S9.**
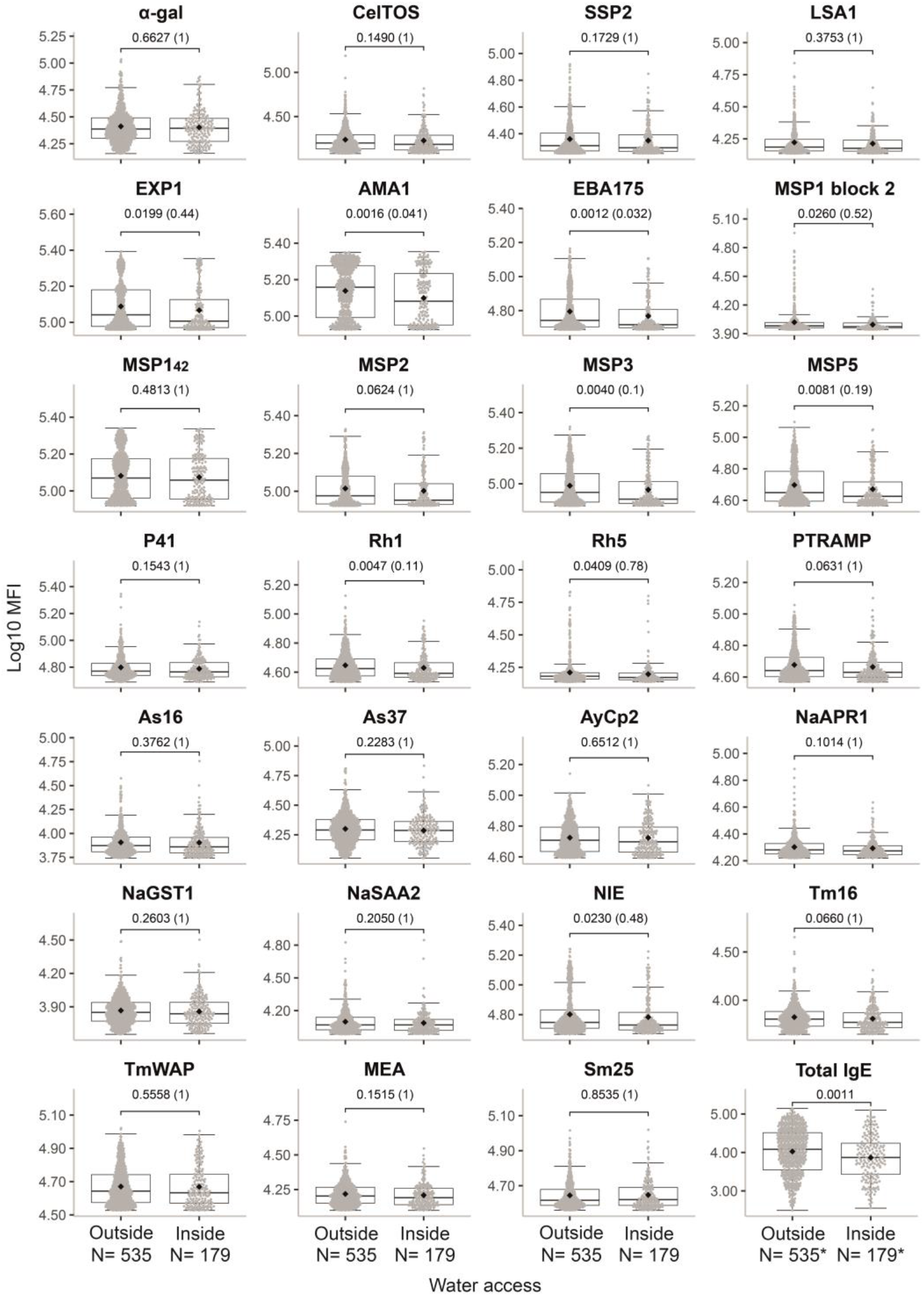
Comparison of antibody responses by accessibility to piped water in the household. Antibody levels expressed as the log_10_-transformed median fluorescence (MFI) of specific IgG to *P. falciparum* antigens (α-gal, CelTOS, SSP2, LSA1, EXP1, AMA1, EBA175, MSP1 block 2, MSP1_42_, MSP2, MSP3, MSP5, P41, Rh5, PTRAMP), helminth antigens (As16 and As37 [*Ascaris lumbricoides*], AyCp2, NaAPR1, NaGST1, NaSAA2 [Hookworm], NIE [*Strongyloides stercoralis*], Tm16 and TmWAP [*Trichuris trichiura*], MEA and Sm25 [*Schistosoma* spp.] or total IgE. The boxplots represent the median (bold line), the mean (black diamond), the 1^st^ and 3^rd^ quartiles (box) and the largest and smallest values within 1.5 times the inter-quartile range (whiskers). Data beyond the end of the whiskers are outliers. Statistical comparison between groups was performed by Wilcoxon rank sum test and the exact p-values are shown. Adjusted p-values by the Benjamini-Hochberg approach are shown in brackets. **\***N= 535 for Outside and N=179 for Inside.

**Figure S10.**
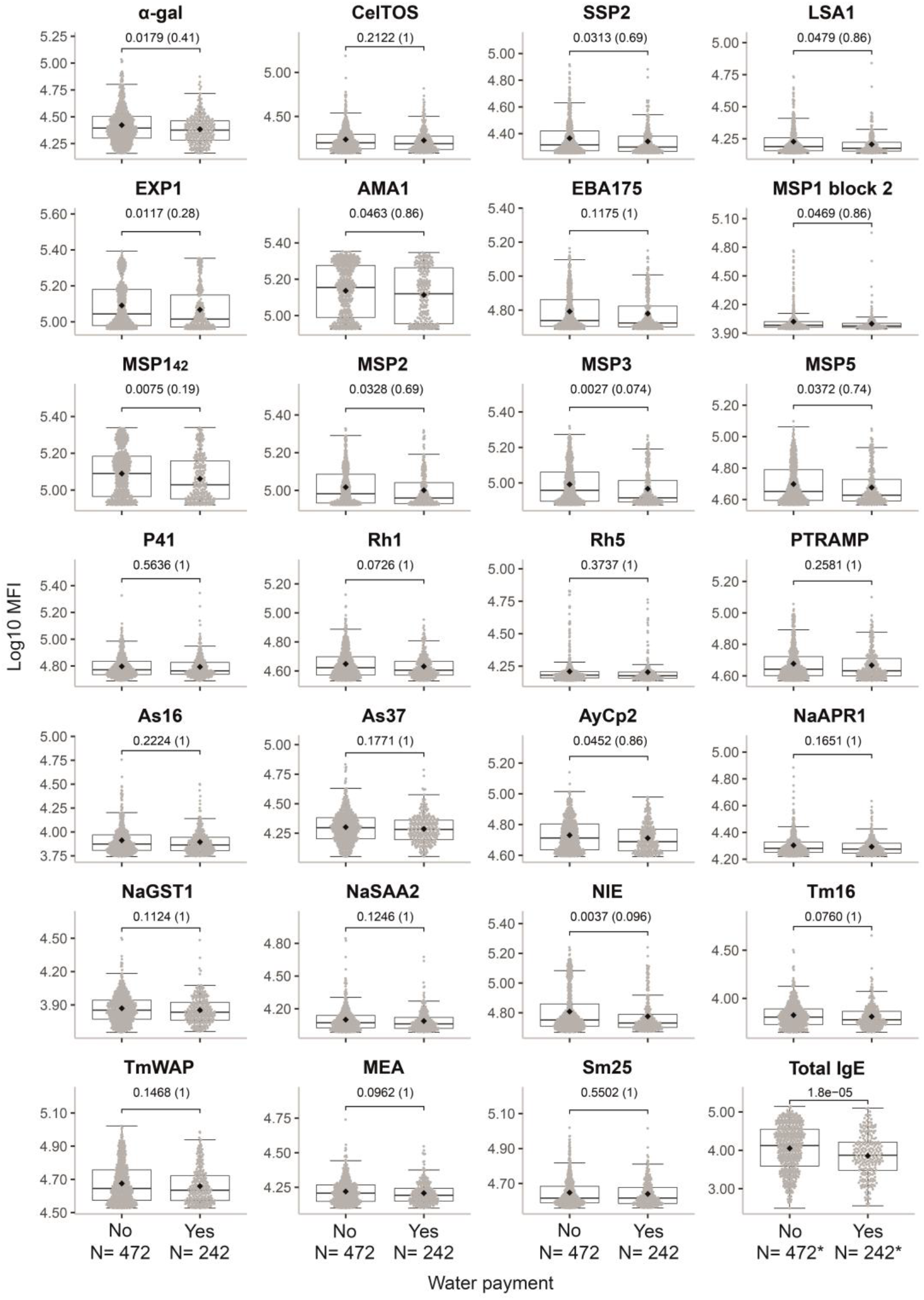
Comparison of antibody responses by payment of piped water. Antibody levels expressed as the log_10_-transformed median fluorescence (MFI) of specific IgG to *P. falciparum* antigens (α-gal, CelTOS, SSP2, LSA1, EXP1, AMA1, EBA175, MSP1 block 2, MSP1_42_, MSP2, MSP3, MSP5, P41, Rh5, PTRAMP), helminth antigens (As16 and As37 [*Ascaris lumbricoides*], AyCp2, NaAPR1, NaGST1, NaSAA2 [Hookworm], NIE [*Strongyloides stercoralis*], Tm16 and TmWAP [*Trichuris trichiura*], MEA and Sm25 [*Schistosoma* spp.] or total IgE. The boxplots represent the median (bold line), the mean (black diamond), the 1^st^ and 3^rd^ quartiles (box) and the largest and smallest values within 1.5 times the inter-quartile range (whiskers). Data beyond the end of the whiskers are outliers. Statistical comparison between groups was performed by Wilcoxon rank sum test and the exact p-values are shown. Adjusted p-values by the Benjamini-Hochberg approach are shown in brackets. **\***N= 471 for No and N= 241 for Yes.

**Figure S11.**
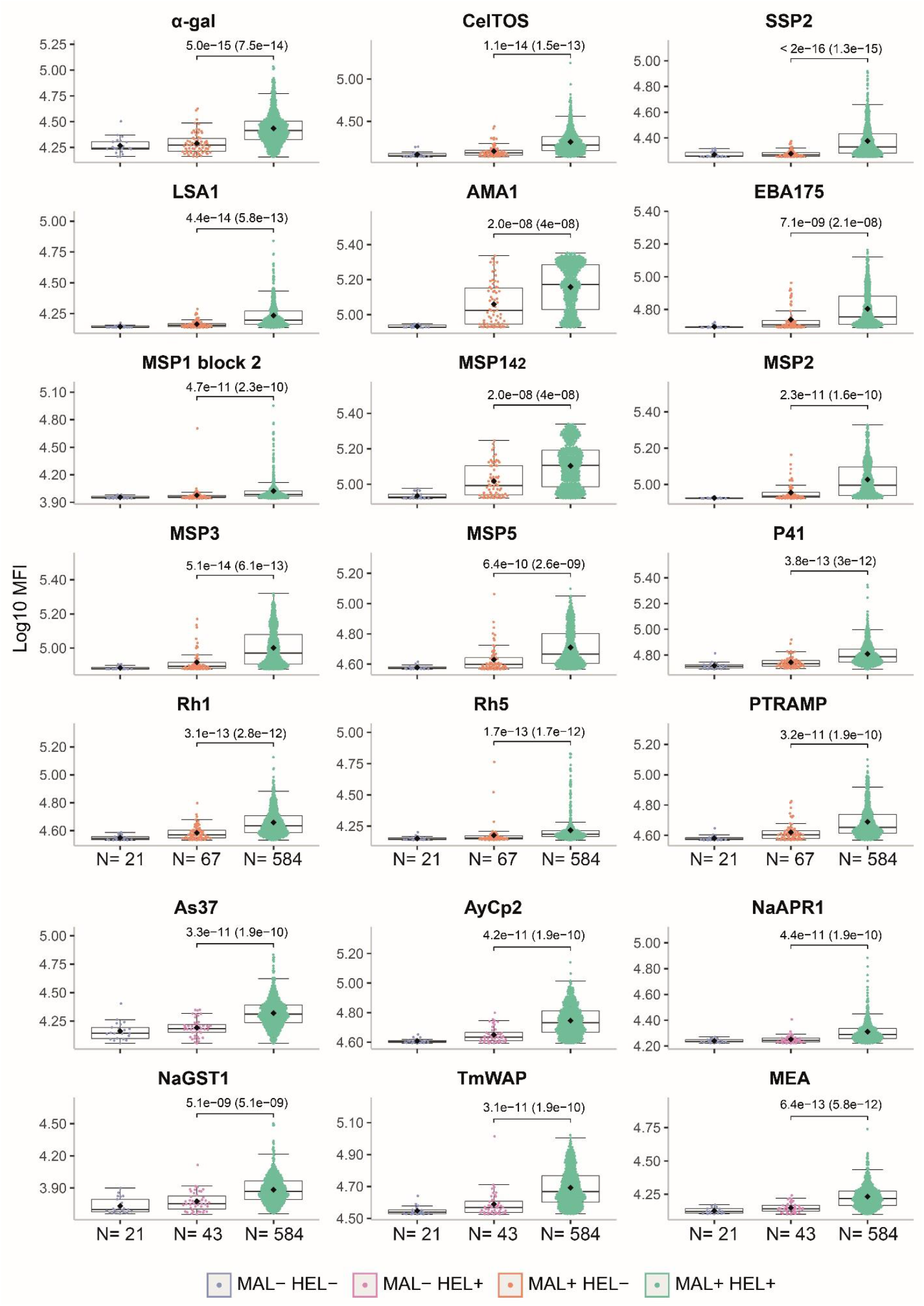
Comparison of antibody responses between exposure/infection groups. Antibody levels expressed as the log_10_-transformed median fluorescence (MFI) of specific IgG to *P. falciparum* antigens (α-gal, CelTOS, SSP2, LSA1, AMA1, EBA175, MSP1 block 2, MSP1_42_, MSP2, MSP3, MSP5, P41, Rh5, PTRAMP), helminth antigens (As37 [*Ascaris lumbricoides*], AyCp2, NaAPR1, NaGST1 [Hookworm], TmWAP [*Trichuris trichiura*], Sm25 [*Schistosoma* spp.]. The boxplots represent the median (bold line), the mean (black diamond), the 1^st^ and 3^rd^ quartiles (box) and the largest and smallest values within 1.5 times the inter-quartile range (whiskers). Data beyond the end of the whiskers are outliers. Statistical comparison between groups was performed by Wilcoxon rank sum test and the exact p-values are shown. Adjusted p-values by the Benjamini-Hochberg approach are shown in brackets. Study groups are shown in colors: no exposure/infection (violet) (MAL- HEL-), only exposure/infection with helminths (pink) (MAL- HEL+), only exposure/infection with *P. falciparum* (orange) (MAL+ HEL-) and coexposure/coinfection with *P. falciparum* and helminths (green) (MAL+ HEL+).

**Figure S12.**
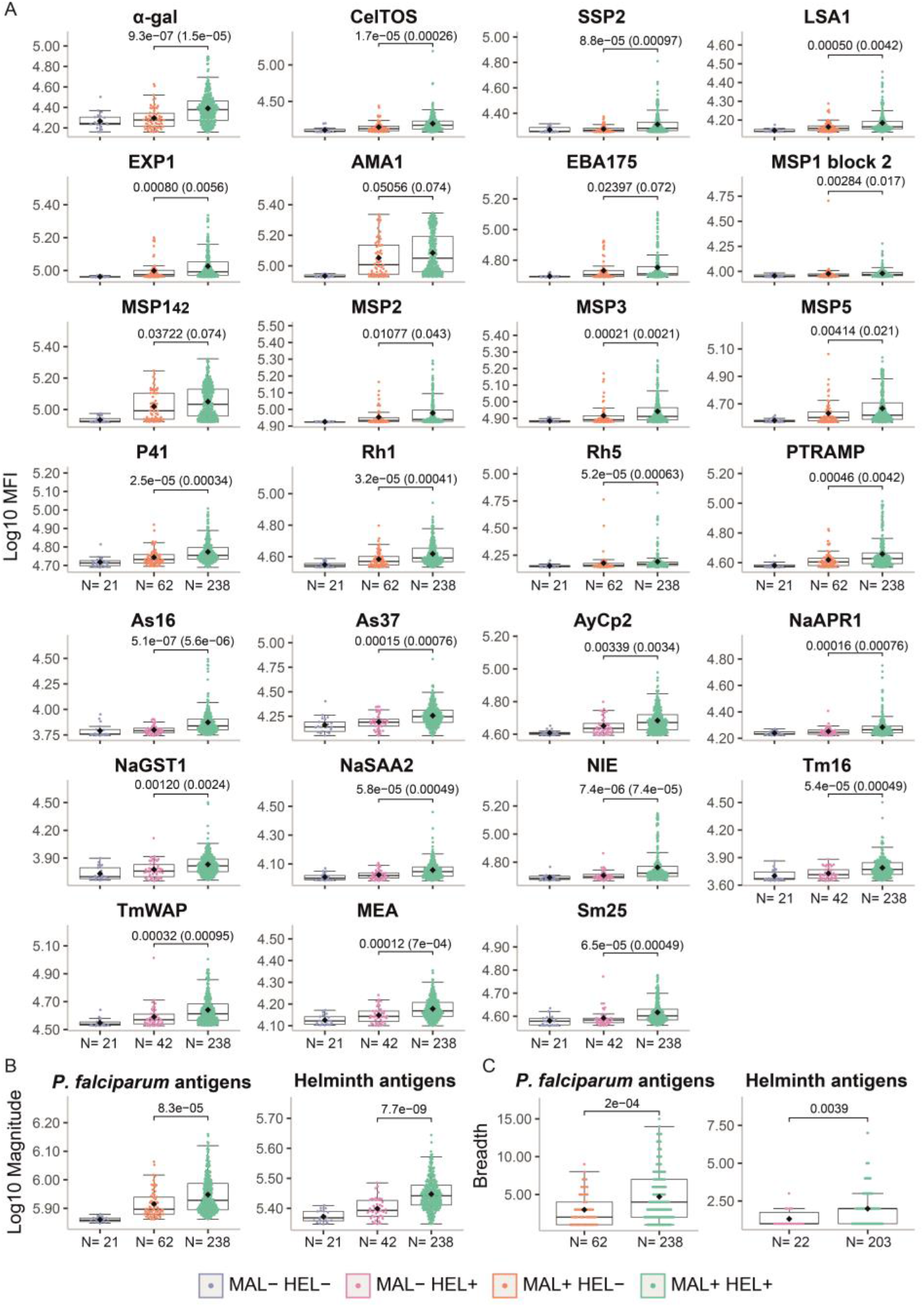
Comparison of antibody responses between exposure/infection groups in children. Antibody responses are expressed as **(A)** the log_10_-transformed median fluorescence (MFI) of specific IgG to *P. falciparum* antigens (α-gal, CelTOS, SSP2, LSA1, EXP1, AMA1, EBA175, MSP1 block 2, MSP1_42_, MSP2, MSP3, MSP5, P41, Rh5, PTRAMP), helminth antigens (As16 and As37 [*Ascaris lumbricoides*], AyCp2, NaAPR1, NaGST1, NaSAA2 [Hookworm], NIE [*Strongyloides stercoralis*], Tm16 and TmWAP [*Trichuris trichiura*], MEA and Sm25 [*Schistosoma* spp.] or total IgE, **(B)** the log_10_-transformed magnitude of response (sum of MFI), and **(C)** the breadth of response (number of seropositive antigen-specific IgG responses). The boxplots represent the median (bold line), the mean (black diamond), the 1^st^ and 3^rd^ quartiles (box) and the largest and smallest values within 1.5 times the inter-quartile range (whiskers). Data beyond the end of the whiskers are outliers. Statistical comparison between groups was performed by Wilcoxon rank sum test and the exact p-values are shown. Adjusted p-values by the Benjamini-Hochberg approach are shown in brackets. Study groups are shown in colors: no exposure/infection (violet) (MAL- HEL-), only exposure/infection with helminths (pink) (MAL- HEL+), only exposure/infection with *P. falciparum* (orange) (MAL+ HEL-) and coexposure/coinfection with *P. falciparum* and helminths (green) (MAL+ HEL+).

**Figure S13.**
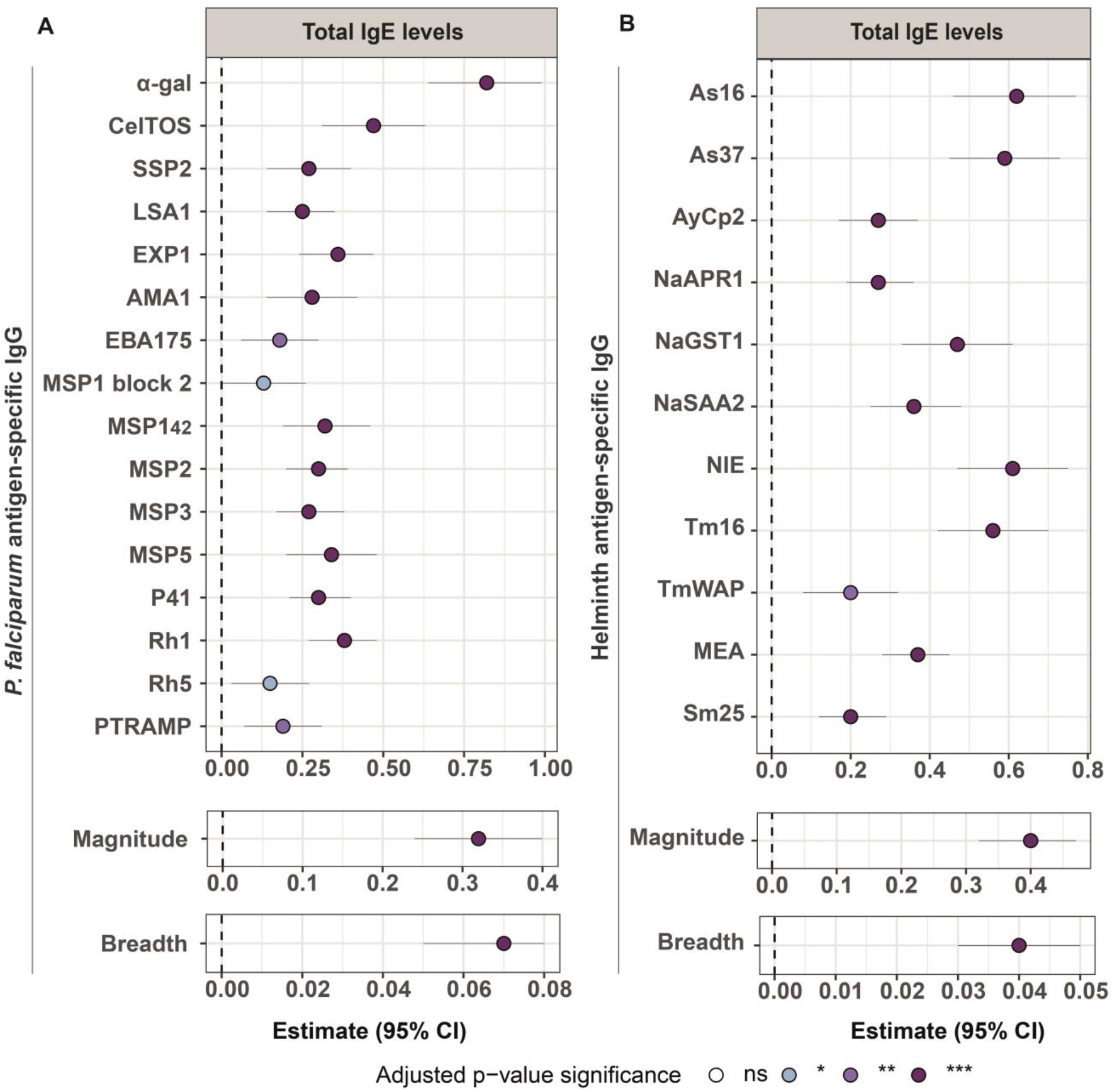
Association of total IgE levels with antibody levels in multivariable linear regression models. Forest plots show the association of total IgE levels with **(A)** *P. falciparum* antigens, magnitude and breadth of response or **(B)** helminth antigens, magnitude (sum of antigen-specific IgG levels) and breadth (number of seropositive antigen-specific IgG responses) of response. Multivariable linear regression models were fitted to calculate the estimates (dots) and 95% confidence intervals (CI) (lines). The represented values are the transformed betas and CI in percentage for antigen and magnitude of response (log-log models). For breadth of response (linear-log models), the betas and 95% CI were transformed to represent the additive effect on the breadth of response of a 10% increase in total IgE levels. The color of the dots represents the p-value significance after adjustment for multiple testing by Benjamini-Hochberg, where ns= not significant, * = p-value ≤ 0.05, ** = p-value ≤ 0.01 and *** = p-value ≤ 0.001. Models were adjusted by age and percentage of *P. falciparum* infection by location for *P. falciparum* antigens and age for helminth antigens.

**Figure S14.**
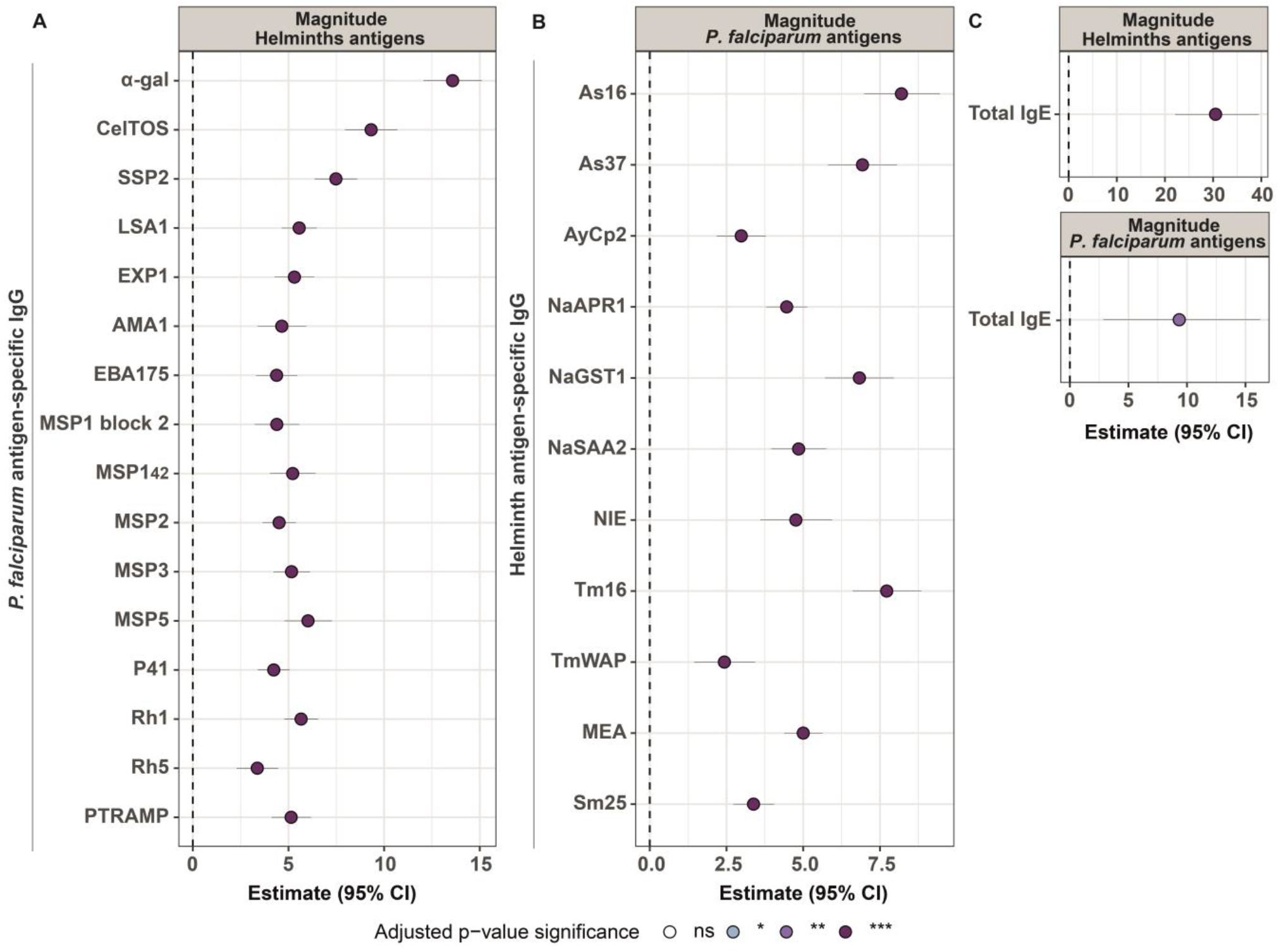
Association of magnitude of response with antibody levels in multivariable linear regression models. Forest plots show the association of **(A)** magnitude of response to helminth antigens with *P. falciparum* antigens, **(B)** magnitude of response to *P. falciparum* antigens with helminth antigens and **(C)** magnitude of response to helminth and *P. falciparum* antigens with total IgE levels. The magnitude of response was calculated as the sum of all specific IgG levels (MFI) to the different antigens belonging to *P. falciparum* or helminths. Multivariable linear regression models were fitted to calculate the estimates (dots) and 95% confidence intervals (CI) (lines). The represented values are the transformed betas and CI in percentage (log-log models). The color of the dots represents the p-value significance after adjustment for multiple testing by Benjamini-Hochberg, where ns= not significant, * = p-value ≤ 0.05, ** = p-value ≤ 0.01 and *** = p-value ≤ 0.001. Models were adjusted by age and percentage of *P. falciparum* infection by location for *P. falciparum* antigens, age for helminth antigens and sex, percentage of helminth infection by location, ownership of latrine, piped water accessibility, payment for piped water and socioeconomic score for total IgE.

**Figure S15.**
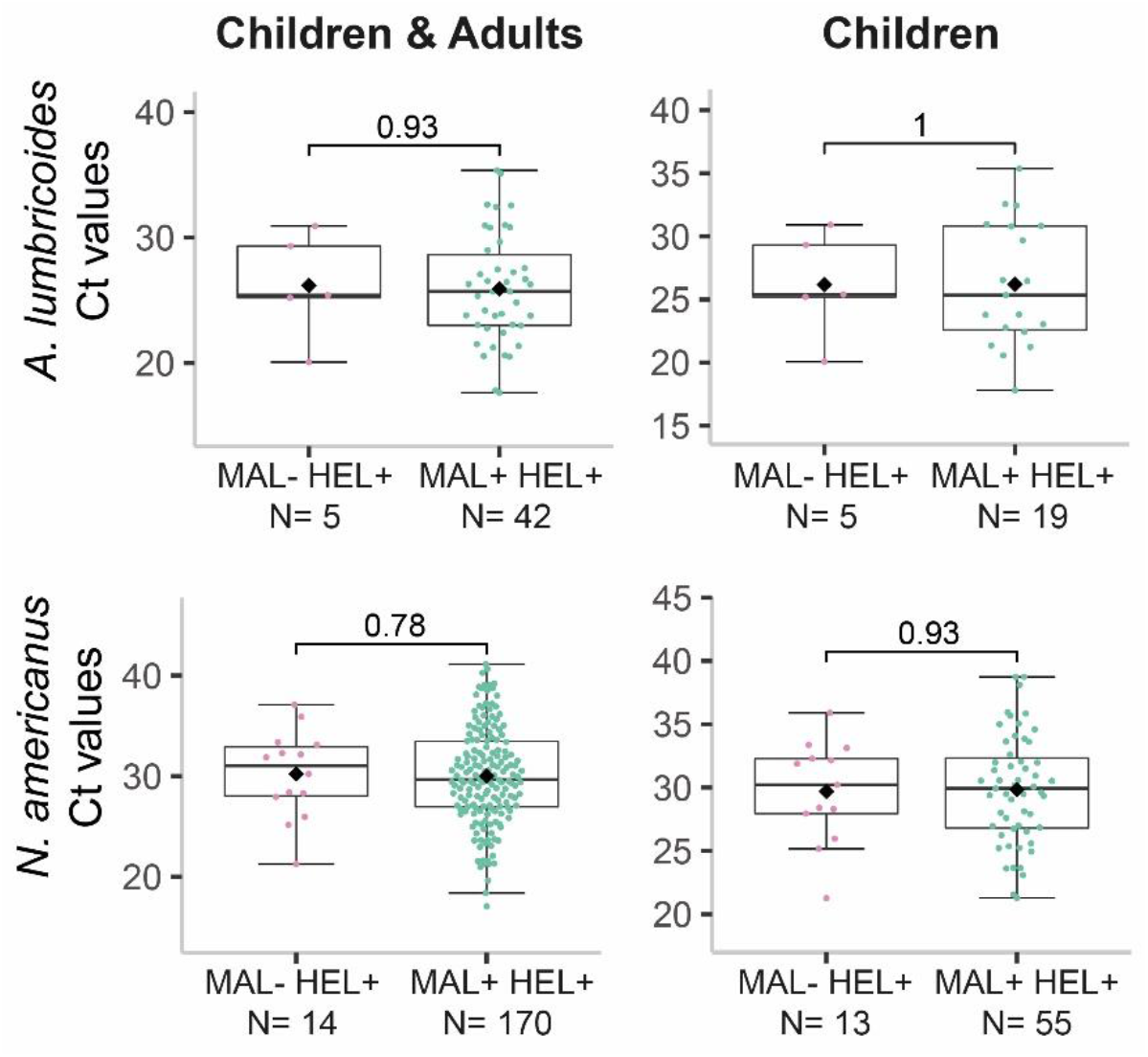
Parasite burden by infection groups. The levels of parasite burden are represented as the Ct values and are compared between the coexposure/coinfection group (MAL+ HEL+) and the single helminth exposure/infection group (MAL- HEL+) for *Ascaris lumbricoides and Necator americanus*. For each parasite, data for all individuals with parasite burden information or only children are displayed. The boxplots represent the median (bold line), the mean (black diamond), the 1^st^ and 3^rd^ quartiles (box) and the largest and smallest values within 1.5 times the inter-quartile range (whiskers). Data beyond the end of the whiskers are outliers. Statistical comparison between groups was performed by Wilcoxon rank sum test and the exact p-values are shown.

**Figure S16.**
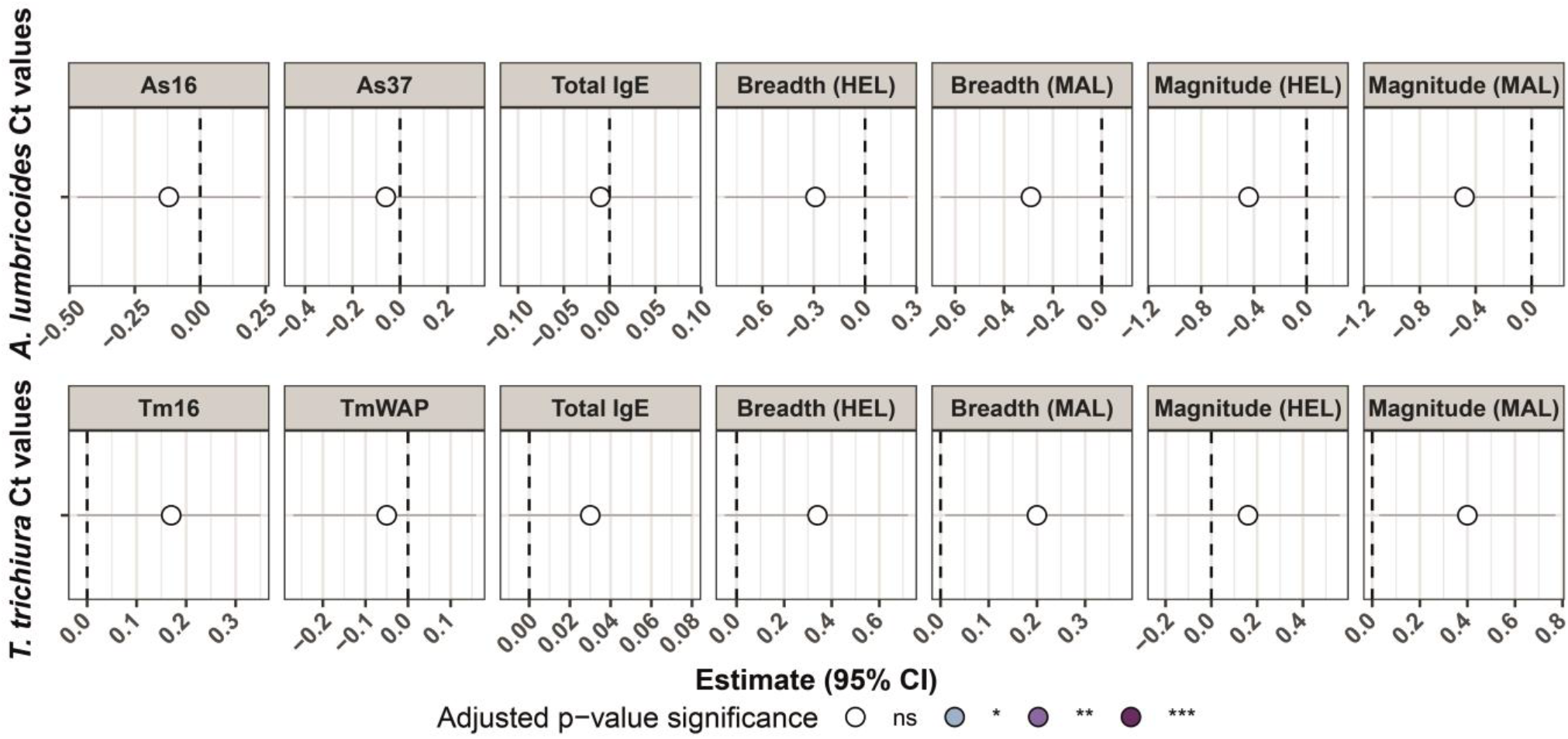
Association of antibody responses with parasite burden in multivariable linear regression models. Forest plots show the association of antibody responses with parasite burden represented as the Ct values of *Ascaris lumbricoides* and *Trichuris trichiura*. For each parasite, linear regression models adjusted by age were fitted, where specific IgG against the antigens from the corresponding parasite, total IgE and magnitude (sum of antigen-specific IgG levels) and breadth (number of seropositive antigen-specific IgG responses) of response were the predictor variables, and the parasite burden was the outcome variable. When the predictor variables were antibody levels or magnitude of response (linear-log models), the estimates (dots) and 95% confidence intervals (CI) (lines) of the models were transformed to represent the additive effect on the Ct values of a 10% increase in the predictor variable. For the breadth of response (linear-linear models), the untransformed betas were used. The color of the dots represents the adjusted p-value significance, where ns= not significant, * = p-value ≤ 0.05, ** = p-value ≤ 0.01 and *** = p-value ≤ 0.001.

## SUMPPLEMENTARY METHODS

### Data pre-processing

For antigen-specific IgG quantification results, we selected the dilution that was within the quantitative range of the PC curve, ideally in the linear part of it. Briefly, the point of the PC curve with the maximum slope was calculated per plate and antigen. The dilution of the sample with the MFI value more similar to this maximum slope point was chosen, as the maximum slope is the most reliable part of the PC curve for quantification. The selection of the dilution was performed after checking for hook effect, since it could disturb the results.

Then, we performed a double normalization to correct a batch effect caused by the use of two different streptavidin-R-phycoerythrin lots. The first normalization was intra-batch to correct for technical artefacts between plates within the same streptavidin-R-phycoerythrin lot. The second normalization was inter-batch to correct for the differences due to the different lots of streptavidin-R-phycoerythrin. First, MFI values after the selection of sample dilution were normalized by multiplying each MFI by the first normalization factor estimated by plate and antigen within each batch. To do so, first an average curve per antigen and batch was calculated by performing the mean of each of the dilution points of the PC curves (i.e.: mean of dilution 1 of all plates within batch 1 for AMA1). Next, the dilution point closest to the maximum slope of the average curve per antigen and batch was identified and denominated as the “normalization dilution point”. This normalization dilution point was then used to calculate the first normalization factor, which was the ratio of the normalization dilution point of each plate curve between the normalization point of the average curve per antigen, plate, and batch. The second normalization factor was obtained by calculating the ratio of the normalization dilution point per antigen between the two batches. Given that the batch 1 included 15 plates and the batch 2 only 6, the reference was batch 1 because its sample size was bigger. Therefore, the second normalization factor was applied to the batch 2 by multiplying the MFI values by the second normalization factor. In this step, the hook effect was also considered. Finally, a correction for the dilution factor was applied to be able to compare samples tested at different dilutions for IgG. The correction factor was obtained with a regression method based on the maximum slope of the batch 1 average curve for each antigen. The regression line has the next formula: ***y*=*mx*+*b***, where ***y*** is the final corrected MFI value, ***m*** is the maximum slope, ***x*** the log10-transformed dilution factor and ***b*** the normalized MFI.

For total IgE, neither selection of dilution, nor a correction by dilution was needed for the data pre-processing. However, normalization was also required due to the batch effect caused by the different streptavidin-R-phycoerythrin lots. MFI values were normalized by calculating the normalization factor of each plate. This factor was obtained by calculating the ratio of the MFI values from the elution control (EC) 1 per plate between the average of all plates. The EC 1 was selected for normalization since the performance of the PC curve was not good enough to use as a reference. Among the ECs, EC 1 had the lowest CV% (EC1 = 6.28%, EC2 = 23.47% and EC3 = 20.86%).

Data from blanks were excluded since they were negligible.

**Table S6.**
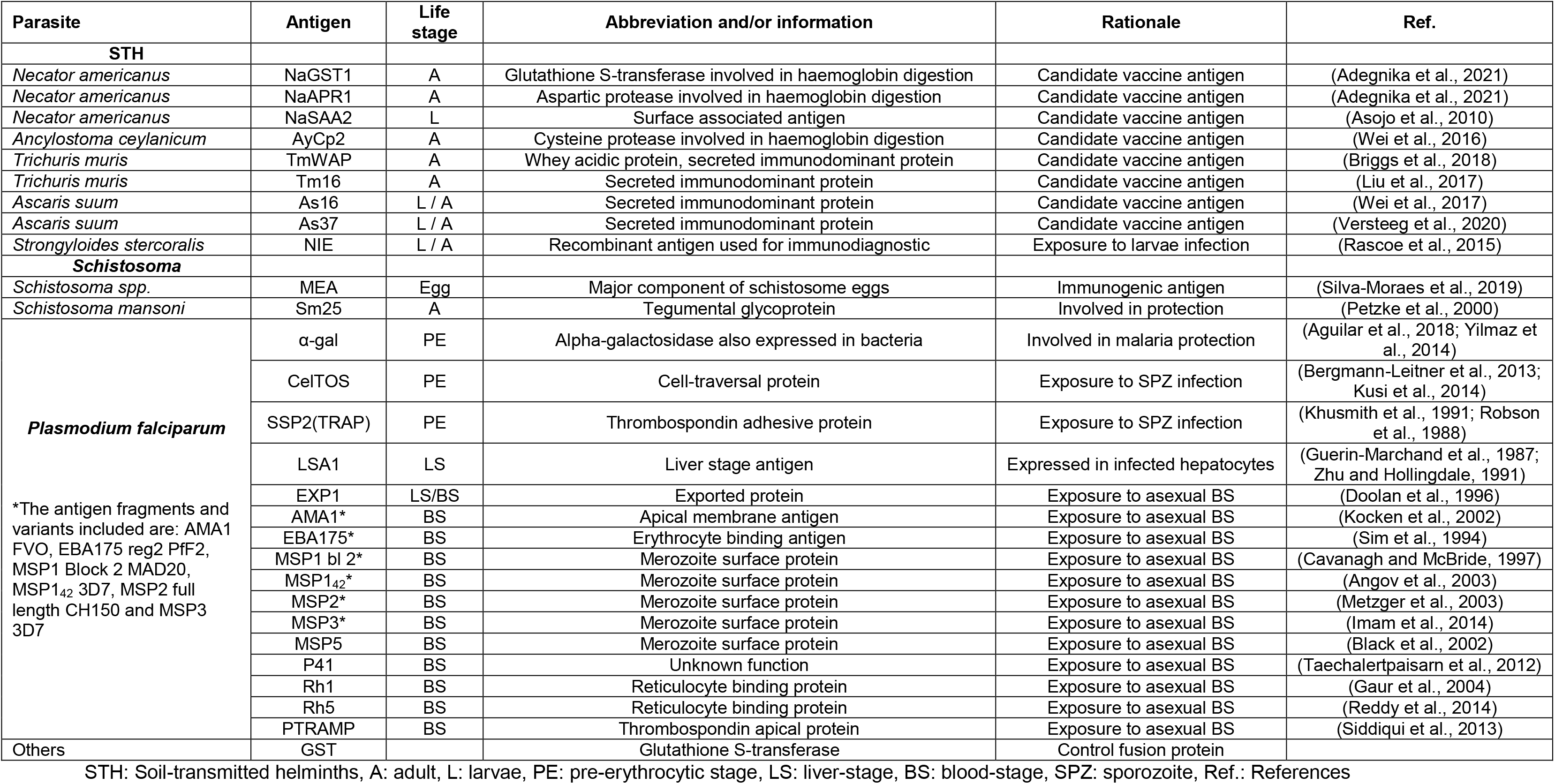
Antigens included in the multiplex panel.

**Table S7.** Panel of antigens with their corresponding optimal coupling conditions.

| Antigen | Coupling concentration | Buffer |
| --- | --- | --- |
| NaGST1 | 100 µg/ml | MES |
| NaAPR1 | 30 µg/ml | MES |
| NaSAA2 | 100 µg/ml | MES |
| AyCp2 | 50 µg/ml | MES |
| TmWAP | 30 µg/ml | MES |
| Tm16 | 50 µg/ml | MES |
| As16 | 50 µg/ml | MES |
| As37 | 75 µg/ml | MES |
| NIE | 50 µg/ml | MES |
| MEA | 30 µg/ml | MES |
| Sm25 | 10 µg/ml | MES |
| α-gal | 30 µg/ml | MES |
| CelTOS | 50 µg/ml | PBS |
| SSP2 or TRAP | 30 µg/ml | PBS |
| LSA1 | 30 µg/ml | PBS |
| EXP1 | 30 µg/ml | MES |
| AMA1 FVO | 30 µg/ml | MES |
| EBA175 reg2 PfF2 | 30 µg/ml | MES |
| MSP1 Block2 MAD20 | 50 µg/ml | MES |
| MSP1 42 3D7 | 30 µg/ml | MES |
| MSP2 full length CH150 | 30 µg/ml | MES |
| MSP3 3D7 | 30 µg/ml | MES |
| MSP5 | 30 µg/ml | MES |
| P41 | 50 µg/ml | MES |
| PfRh1 | 30 µg/ml | MES |
| PfRh5 | 30 µg/ml | MES |
| PTRAMP | 30 µg/ml | MES |
| GST | 50 µg/ml | MES |
| anti-IgE | 25 µg/ml | MES |

MES: Morpholineethane sulfonic acid

## REFERENCES

Abbate JL, Ezenwa VO, Guégan J-F, Choisy M, Nacher M, Roche B. 2018. Disentangling complex parasite interactions: Protection against cerebral malaria by one helminth species is jeopardized by co-infection with another. PLoS Negl Trop Dis 12:e0006483.

Afolabi MO, Ale BM, Dabira ED, Agbla SC, Bustinduy AL, Ndiaye JLA, Greenwood B. 2021. Malaria and helminth co-infections in children living in endemic countries: A systematic review with meta-analysis. PLoS Negl Trop Dis 15:e0009138. doi:10.1371/journal.pntd.0009138

Amoani B, Adu B, Frempong MT, Sarkodie-Addo T, Victor Nuvor S, Abu EK, Harrison LM, Cappello M, Gyan B, Wilson MD. 2019. Cytokine profiles of Necator americanus and Plasmodium falciparum co-infected patients in rural Ghana. Cytokine X 1:100014. doi:10.1016/j.cytox.2019.100014

Amoani B, Gyan B, Sakyi SA, Abu EK, Nuvor SV, Barnes P, Sarkodie-Addo T, Ahenkorah B, Sewor C, Dwomoh D, Theisen M, Cappello M, Wilson MD, Adu B. 2021. Effect of hookworm infection and anthelmintic treatment on naturally acquired antibody responses against the GMZ2 malaria vaccine candidate and constituent antigens. BMC Infect Dis 21:332. doi:10.1186/s12879-021-06027-5

Artis D, Grencis RK. 2008. The intestinal epithelium: sensors to effectors in nematode infection. Mucosal Immunol 1:252–264. doi:10.1038/mi.2008.21

Ateba-Ngoa U, Jones S, Zinsou JF, Honkpehedji J, Adegnika AA, Agobe JCD, Massinga-Loembe M, Mordmüller B, Bousema T, Yazdanbakhsh M. 2016. Associations between Helminth Infections, Plasmodium falciparum Parasite Carriage and Antibody Responses to Sexual and Asexual Stage Malarial Antigens. Am J Trop Med Hyg 95:394–400. doi:10.4269/ajtmh.15-0703

Babamale OA, Ugbomoiko US, Heukelbach J. 2018. High prevalence of Plasmodium falciparum and soil-transmitted helminth co-infections in a periurban community in Kwara State, Nigeria. J Infect Public Health 11:48–53. doi:10.1016/j.jiph.2017.03.002

Bereczky S, Montgomery SM, Troye-Blomberg M, Rooth I, Shaw M-A, Färnert A. 2004. Elevated anti-malarial IgE in asymptomatic individuals is associated with reduced risk for subsequent clinical malaria. Int J Parasitol 34:935–942. doi:10.1016/j.ijpara.2004.04.007

Bouharoun-Tayoun H, Druilhe P. 1992. Plasmodium falciparum malaria: evidence for an isotype imbalance which may be responsible for delayed acquisition of protective immunity. Infect Immun 60:1473–1481. doi:10.1128/IAI.60.4.1473-1481.1992

Briand V, Watier L, LE Hesran J-Y, Garcia A, Cot M. 2005. Coinfection with Plasmodium falciparum and schistosoma haematobium: protective effect of schistosomiasis on malaria in senegalese children? Am J Trop Med Hyg 72:702–707.

Brooker S, Akhwale W, Pullan R, Estambale B, Clarke SE, Snow RW, Hotez PJ. 2007. Epidemiology of plasmodium-helminth co-infection in Africa: populations at risk, potential impact on anemia, and prospects for combining control. Am J Trop Med Hyg 77:88–98.

Brooker S, Clements ACA, Hotez PJ, Hay SI, Tatem AJ, Bundy DAP, Snow RW. 2006. The co-distribution of Plasmodium falciparum and hookworm among African schoolchildren. Malar J 5:99. doi:10.1186/1475-2875-5-99

Brutus L, Watier L, Briand V, Hanitrasoamampionona V, Razanatsoarilala H, Cot M. 2006. Parasitic co-infections: does Ascaris lumbricoides protect against Plasmodium falciparum infection? Am J Trop Med Hyg 75:194–198.

Brutus L, Watier L, Hanitrasoamampionona V, Razanatsoarilala H, Cot M. 2007. Confirmation of the protective effect of Ascaris lumbricoides on Plasmodium falciparum infection: results of a randomized trial in Madagascar. Am J Trop Med Hyg 77:1091–1095.

Calissano C, Modiano D, Sirima BS, Konate A, Sanou I, Sawadogo A, Perlmann H, Troye-Blomberg M, Perlmann P. 2003. IgE antibodies to Plasmodium falciparum and severity of malaria in children of one ethnic group living in Burkina Faso. Am J Trop Med Hyg 69:31–35.

Clarke E, Sherrill-Mix SA. 2017. ggbeeswarm: Categorical scatter (violin point) plots. R Packag version 060. https://rdrr.io/cran/ggbeeswarm/

Cohen S, Mcgregor IA, Carrington S. 1961. Gamma-globulin and acquired immunity to human malaria. Nature 192:733–737. doi:10.1038/192733a0

Cooper PJ, Alexander N, Moncayo A-L, Benitez SM, Chico ME, Vaca MG, Griffin GE. 2008. Environmental determinants of total IgE among school children living in the rural Tropics: importance of geohelminth infections and effect of anthelmintic treatment. BMC Immunol 9:33. doi:10.1186/1471-2172-9-33

Courtin D, Djilali-Saïah A, Milet J, Soulard V, Gaye O, Migot-Nabias F, Sauerwein R, Garcia A, Luty AJF. 2011. Schistosoma haematobium infection affects Plasmodium falciparum-specific IgG responses associated with protection against malaria. Parasite Immunol 33:124–131. doi:10.1111/j.1365-3024.2010.01267.x

Degarege A, Animut A, Legesse M, Erko B. 2009. Malaria severity status in patients with soil-transmitted helminth infections. Acta Trop 112:8–11. doi:10.1016/j.actatropica.2009.05.019

Degarege A, Erko B. 2016. Epidemiology of Plasmodium and Helminth Coinfection and Possible Reasons for Heterogeneity. Biomed Res Int 2016:3083568. doi:10.1155/2016/3083568

Degarege A, Legesse M, Medhin G, Animut A, Erko B. 2012. Malaria and related outcomes in patients with intestinal helminths: a cross-sectional study. BMC Infect Dis 12:291. doi:10.1186/1471-2334-12-291

Degarege A, Veledar E, Degarege D, Erko B, Nacher M, Madhivanan P. 2016. Plasmodium falciparum and soil-transmitted helminth co-infections among children in sub-Saharan Africa: a systematic review and meta-analysis. Parasit Vectors 9:344. doi:10.1186/s13071-016-1594-2

Diallo TO, Remoue F, Gaayeb L, Schacht AM, Charrier N, de Clerck D, Dompnier JP, Pillet S, Garraud O, N’Diaye AA, Riveau G. 2010. Schistosomiasis coinfection in children influences acquired immune response against plasmodium falciparum Malaria antigens. PLoS One 5:1–7. doi:10.1371/journal.pone.0012764

Donati D, Zhang LP, Chêne A, Chen Q, Flick K, Nyström M, Wahlgren M, Bejarano MT. 2004. Identification of a polyclonal B-cell activator in Plasmodium falciparum. Infect Immun 72:5412–5418. doi:10.1128/IAI.72.9.5412-5418.2004

Druilhe P, Tall A, Sokhna C. 2005. Worms can worsen malaria: towards a new means to roll back malaria? Trends Parasitol 21:359–362. doi:10.1016/j.pt.2005.06.011

Duarte J, Deshpande P, Guiyedi V, Mécheri S, Fesel C, Cazenave P-A, Mishra GC, Kombila M, Pied S. 2007. Total and functional parasite specific IgE responses in Plasmodium falciparum-infected patients exhibiting different clinical status. Malar J 6:1. doi:10.1186/1475-2875-6-1

Duarte J, Herbert F, Guiyedi V, Franetich J-F, Roland J, Cazenave P-A, Mazier D, Kombila M, Fesel C, Pied S. 2012. High levels of immunoglobulin E autoantibody to 14-3-3 epsilon protein correlate with protection against severe Plasmodium falciparum malaria. J Infect Dis 206:1781–1789. doi:10.1093/infdis/jis595

Easton A V, Raciny-Aleman M, Liu V, Ruan E, Marier C, Heguy A, Yasnot MF, Rodriguez A, Loke P. 2020. Immune Response and Microbiota Profiles during Coinfection with Plasmodium vivax and Soil-Transmitted Helminths. MBio 11:e01705–20. doi:10.1128/mBio.01705-20

Efunshile AM, Olawale T, Stensvold CR, Kurtzhals JAL, König B. 2015. Epidemiological study of the association between malaria and helminth infections in Nigeria. Am J Trop Med Hyg 92:578–582. doi:10.4269/ajtmh.14-0548

Farouk SE, Dolo A, Bereczky S, Kouriba B, Maiga B, Färnert A, Perlmann H, Hayano M, Montgomery SM, Doumbo OK, Troye-Blomberg M. 2005. Different antibody- and cytokine-mediated responses to Plasmodium falciparum parasite in two sympatric ethnic tribes living in Mali. Microbes Infect 7:110–117. doi:10.1016/j.micinf.2004.09.012

Fonseca AM, Quinto L, Jiménez A, González R, Bardají A, Maculuve S, Dobaño C, Rupérez M, Vala A, Aponte JJ, Sevene E, Macete E, Menéndez C, Mayor A. 2017. Multiplexing detection of IgG against Plasmodium falciparum pregnancy-specific antigens. PLoS One 12:e0181150. doi:10.1371/journal.pone.0181150

Geha RS. 1984. Human IgE. J Allergy Clin Immunol 74:109–120. doi:10.1016/0091-6749(84)90270-7

Grau-Pujol B, Marti-Soler H, Escola V, Demontis M, Jamine JC. 2021. Towards soil-transmitted helminths transmission interruption: the impact of diagnostic tools on infection prediction in a low intensity setting in Southern Mozambique.

Harris N, Gause WC. 2011. To B or not to B: B cells and the Th2-type immune response to helminths. Trends Immunol 32:80–88. doi:10.1016/j.it.2010.11.005

Hartgers FC, Yazdanbakhsh M. 2006. Co-infection of helminths and malaria: modulation of the immune responses to malaria. Parasite Immunol 28:497–506. doi:https://doi.org/10.1111/j.1365-3024.2006.00901.x

Kaisar MMM, Brienen EAT, Djuardi Y, Sartono E, Yazdanbakhsh M, Verweij JJ, Supali T, VAN Lieshout L. 2017. Improved diagnosis of Trichuris trichiura by using a bead-beating procedure on ethanol preserved stool samples prior to DNA isolation and the performance of multiplex real-time PCR for intestinal parasites. Parasitology 144:965–974. doi:10.1017/S0031182017000129

Kassambara A. 2020. “ggplot2” Based Publication Ready Plots [R package ggpubr version 0.4.0].

Kotowicz K, Callard RE. 1993. Human immunoglobulin class and IgG subclass regulation: dual action of interleukin-4. Eur J Immunol 23:2250–2256. doi:10.1002/eji.1830230930

Langhorne J, Cross C, Seixas E, Li C, von der Weid T. 1998. A role for B cells in the development of T cell helper function in a malaria infection in mice. Proc Natl Acad Sci U S A 95:1730–1734. doi:10.1073/pnas.95.4.1730

Le Hesran J-Y, Akiana J, Ndiaye EHM, Dia M, Senghor P, Konate L. 2004. Severe malaria attack is associated with high prevalence of Ascaris lumbricoides infection among children in rural Senegal. Trans R Soc Trop Med Hyg 98:397–399. doi:10.1016/j.trstmh.2003.10.009

Lemaitre M, Watier L, Briand V, Garcia A, Le Hesran JY, Cot M. 2014. Coinfection with Plasmodium falciparum and Schistosoma haematobium: Additional evidence of the protective effect of schistosomiasis on malaria in Senegalese children. Am J Trop Med Hyg 90:329–334. doi:10.4269/ajtmh.12-0431

Lyke KE, Dicko A, Dabo A, Sangare L, Kone A, Coulibaly D, Guindo A, Traore K, Daou M, Diarra I, Sztein MB, Plowe C V, Doumbo OK. 2005. Association of Schistosoma haematobium infection with protection against acute Plasmodium falciparum malaria in Malian children. Am J Trop Med Hyg 73:1124–1130.

Maeno Y, Perlmann P, Perlmann H, Kusuhara Y, Taniguchi K, Nakabayashi T, Win K, Looareesuwan S, Aikawa M. 2000. IgE deposition in brain microvessels and on parasitized erythrocytes from cerebral malaria patients. Am J Trop Med Hyg 63:128–132. doi:10.4269/ajtmh.2000.63.128

Maizels RM, Balic A, Gomez-Escobar N, Nair M, Taylor MD, Allen JE. 2004. Helminth parasites – masters of regulation. Immunol Rev 201:89–116. doi:https://doi.org/10.1111/j.0105-2896.2004.00191.x

Maizels RM, Yazdanbakhsh M. 2003. Immune regulation by helminth parasites: cellular and molecular mechanisms. Nat Rev Immunol 3:733–744. doi:10.1038/nri1183

Malaguarnera L, Musumeci S. 2002. The immune response to Plasmodium falciparum malaria. Lancet Infect Dis 2:472–478. doi:https://doi.org/10.1016/S1473-3099(02)00344-4

Mangano VD, Bianchi C, Ouedraogo M, Kabore Y, Corran P, Silva N, Sirima SB, Nebie I, Bruschi F, Modiano D. 2020. Antibody response to Schistosoma haematobium and other helminth species in malaria-exposed populations from Burkina Faso. Acta Trop 205:105381. doi:https://doi.org/10.1016/j.actatropica.2020.105381

Mayor A, Serra-Casas E, Bardají A, Sanz S, Puyol L, Cisteró P, Sigauque B, Mandomando I, Aponte JJ, Alonso PL, Menéndez C. 2009. Sub-microscopic infections and long-term recrudescence of Plasmodium falciparum in Mozambican pregnant women. Malar J 8:9. doi:10.1186/1475-2875-8-9

McSorley HJ, Maizels RM. 2012. Helminth infections and host immune regulation. Clin Microbiol Rev 25:585–608. doi:10.1128/CMR.05040-11

Miranda GS, Resende SD, Cardoso DT, Camelo GMA, Silva JKAO, de Castro VN, Geiger SM, Carneiro M, Negrão-Corrêa D. 2021. Previous History of American Tegumentary Leishmaniasis Alters Susceptibility and Immune Response Against Schistosoma mansoni Infection in Humans. Front Immunol 12:630934. doi:10.3389/fimmu.2021.630934

Montes CL, Acosta-Rodríguez E V, Merino MC, Bermejo DA, Gruppi A. 2007. Polyclonal B cell activation in infections: infectious agents’ devilry or defense mechanism of the host? J Leukoc Biol 82:1027–1032. doi:10.1189/jlb.0407214

Moon K-W. 2020. Make Interactive “ggplot2”. Extension to “ggplot2” and “ggiraph” [R package ggiraphExtra version 0.3.0].

Mulu A, Kassu A, Legesse M, Erko B, Nigussie D, Shimelis T, Belyhun Y, Moges B, Ota F, Elias D. 2014. Helminths and malaria co-infections are associated with elevated serum IgE. Parasit Vectors 7:240. doi:10.1186/1756-3305-7-240

Murray J, Murray A, Murray M, Murray C. 1978. The biological suppression of malaria: an ecological and nutritional interrelationship of a host and two parasites. Am J Clin Nutr 31:1363–1366. doi:10.1093/ajcn/31.8.1363

Mwangi TW, Bethony JM, Brooker S. 2006. Malaria and helminth interactions in humans: an epidemiological viewpoint. Ann Trop Med Parasitol 100:551–570. doi:10.1179/136485906X118468

Nacher M, Gay F, Singhasivanon P, Krudsood S, Treeprasertsuk S, Mazier D, Vouldoukis I, Looareesuwan S. 2000. Ascaris lumbricoides infection is associated with protection from cerebral malaria. Parasite Immunol 22:107–113. doi:https://doi.org/10.1046/j.1365-3024.2000.00284.x

Nacher M, Singhasivanon P, Gay F, Silachomroon U, Phumratanaprapin W, Looareesuwan S. 2001a. Contemporaneous and successive mixed Plasmodium falciparum and Plasmodium vivax infections are associated with Ascaris lumbricoides: an immunomodulating effect? J Parasitol 87:912–915. doi:10.1645/0022-3395(2001)087[0912:CASMPF]2.0.CO;2

Nacher M, Singhasivanon P, Silachamroon U, Treeprasertsuk S, Vannaphan S, Traore B, Gay F, Looareesuwan S. 2001b. Helminth infections are associated with protection from malaria-related acute renal failure and jaundice in Thailand. Am J Trop Med Hyg 65:834–836. doi:10.4269/ajtmh.2001.65.834

Nacher M, Singhasivanon P, Traore B, Vannaphan S, Gay F, Chindanond D, Franetich J, Mazier D, Looareesuwan S. 2002a. Helminth infections are associated with protection from cerebral malaria and increased nitrogen derivatives concentrations in Thailand. Am J Trop Med Hyg 66:304–309. doi:10.4269/ajtmh.2002.66.304

Nacher M, Singhasivanon P, Yimsamran S, Manibunyong W, Thanyavanich N, Wuthisen R, Looareesuwan S. 2002b. Intestinal helminth infections are associated with increased incidence of Plasmodium falciparum malaria in Thailand. J Parasitol 88:55–58. doi:10.1645/0022-3395(2002)088[0055:IHIAAW]2.0.CO;2

Njenga SM, Kanyi HM, Arnold BF, Matendechero SH, Onsongo JK, Won KY, Priest JW. 2020. Integrated Cross-Sectional Multiplex Serosurveillance of IgG Antibody Responses to Parasitic Diseases and Vaccines in Coastal Kenya. Am J Trop Med Hyg 102:164–176. doi:10.4269/ajtmh.19-0365

Ntonifor HN, Chewa JS, Oumar M, Mbouobda HD. 2021. Intestinal helminths as predictors of some malaria clinical outcomes and IL-1β levels in outpatients attending two public hospitals in Bamenda, North West Cameroon. PLoS Negl Trop Dis 15:e0009174. doi:10.1371/journal.pntd.0009174

Pearce EJ, MacDonald AS. 2002. The immunobiology of schistosomiasis. Nat Rev Immunol 2:499–511. doi:10.1038/nri843

Perlmann H, Helmby H, Hagstedt M, Carlson J, Larsson PH, Troye-Blomberg M, Perlmann P. 1994. IgE elevation and IgE anti-malarial antibodies in Plasmodium falciparum malaria: association of high IgE levels with cerebral malaria. Clin Exp Immunol 97:284–292. doi:10.1111/j.1365-2249.1994.tb06082.x

Perlmann P, Perlmann H, Flyg BW, Hagstedt M, Elghazali G, Worku S, Fernandez V, Rutta AS, Troye-Blomberg M. 1997. Immunoglobulin E, a pathogenic factor in Plasmodium falciparum malaria. Infect Immun 65:116–121. doi:10.1128/IAI.65.1.116-121.1997

Perlmann P, Perlmann H, Looareesuwan S, Krudsood S, Kano S, Matsumoto Y, Brittenham G, Troye-Blomberg M, Aikawa M. 2000. Contrasting functions of IgG and IgE antimalarial antibodies in uncomplicated and severe Plasmodium falciparum malaria. Am J Trop Med Hyg 62:373–377. doi:10.4269/ajtmh.2000.62.373

Porto AF, Neva FA, Bittencourt H, Lisboa W, Thompson R, Alcântara L, Carvalho EM. 2001. HTLV-1 decreases Th2 type of immune response in patients with strongyloidiasis. Parasite Immunol 23:503–507. doi:10.1046/j.1365-3024.2001.00407.x

Quinnell RJ, Pritchard DI, Raiko A, Brown AP, Shaw M. 2004. Immune Responses in Human Necatoriasis: Association between Interleukin-5 Responses and Resistance to Reinfection. J Infect Dis 190:430–438. doi:10.1086/422256

Remoue F, Diallo TO, Angeli V, Hervé M, de Clercq D, Schacht AM, Charrier N, Capron M, Vercruysse J, Ly A, Capron A, Riveau G. 2003. Malaria co-infection in children influences antibody response to schistosome antigens and inflammatory markers associated with morbidity. Trans R Soc Trop Med Hyg 97:361–364. doi:10.1016/S0035-9203(03)90170-2

Roussilhon C, Brasseur P, Agnamey P, Pérignon J-L, Druilhe P. 2010. Understanding human-Plasmodium falciparum immune interactions uncovers the immunological role of worms. PLoS One 5:e9309. doi:10.1371/journal.pone.0009309

Salazar-Castañón VH, Juárez-Avelar I, Legorreta-Herrera M, Govezensky T, Rodriguez-Sosa M. 2018. Co-infection: the outcome of Plasmodium infection differs according to the time of pre-existing helminth infection. Parasitol Res 117:2767–2784. doi:10.1007/s00436-018-5965-9

Scrucca L, Fop M, Murphy TB, Raftery AE. 2016. mclust 5: Clustering, Classification and Density Estimation Using Gaussian Finite Mixture Models. R J 8:289–317.

Seka-Seka J, Brouh Y, Yapo-Crézoit AC, Atseye NH. 2004. The role of serum immunoglobulin E in the pathogenesis of Plasmodium falciparum malaria in Ivorian children. Scand J Immunol 59:228–230. doi:10.1111/j.0300-9475.2004.01337.x

Sokhna C, Le Hesran J-Y, Mbaye PA, Akiana J, Camara P, Diop M, Ly A, Druilhe P. 2004. Increase of malaria attacks among children presenting concomitant infection by Schistosoma mansoni in Senegal. Malar J 3:43. doi:10.1186/1475-2875-3-43

Spiegel A, Tall A, Raphenon G, Trape J-F, Druilhe P. 2003. Increased frequency of malaria attacks in subjects co-infected by intestinal worms and Plasmodium falciparum malaria. Trans R Soc Trop Med Hyg 97:198–199. doi:10.1016/S0035-9203(03)90117-9

Subirana I, Sanz H, Vila J. 2014. Building Bivariate tables: The compareGroups package for R. J Stat Softw 57:1–16. doi:10.18637/jss.v057.i12

Tangteerawatana P, Montgomery SM, Perlmann H, Looareesuwan S, Troye-Blomberg M, Khusmith S. 2007. Differential regulation of IgG subclasses and IgE antimalarial antibody responses in complicated and uncomplicated Plasmodium falciparum malaria. Parasite Immunol 29:475–483. doi:10.1111/j.1365-3024.2007.00965.x

Taylor SM, Mayor A, Mombo-Ngoma G, Kenguele HM, Ouédraogo S, Ndam NT, Mkali H, Mwangoka G, Valecha N, Singh JPN, Clark MA, Verweij JJ, Adegnika AA, Severini C, Menegon M, Macete E, Menendez C, Cisteró P, Njie F, Affara M, Otieno K, Kariuki S, ter Kuile FO, Meshnick SR. 2014. A quality control program within a clinical trial Consortium for PCR protocols to detect Plasmodium species. J Clin Microbiol 52:2144–2149. doi:10.1128/JCM.00565-14

Tokplonou L, Nouatin O, Sonon P, M’po G, Glitho S, Agniwo P, Gonzalez-Ortiz D, Tchégninougbo T, Ayitchédji A, Favier B, Donadi EA, Milet J, Luty AJF, Massougbodji A, Garcia A, Ibikounlé M, Courtin D. 2020. Schistosoma haematobium infection modulates Plasmodium falciparum parasite density and antimalarial antibody responses. Parasite Immunol 42:e12702. doi:https://doi.org/10.1111/pim.12702

Tran TM, Guha R, Portugal S, Skinner J, Ongoiba A, Bhardwaj J, Jones M, Moebius J, Venepally P, Doumbo S, DeRiso EA, Li S, Vijayan K, Anzick SL, Hart GT, O’Connell EM, Doumbo OK, Kaushansky A, Alter G, Felgner PL, Lorenzi H, Kayentao K, Traore B, Kirkness EF, Crompton PD. 2019. A Molecular Signature in Blood Reveals a Role for p53 in Regulating Malaria-Induced Inflammation. Immunity 51:750–765.e10. doi:10.1016/j.immuni.2019.08.009

Troye-Blomberg M, Riley EM, Kabilan L, Holmberg M, Perlmann H, Andersson U, Heusser CH, Perlmann P. 1990. Production by activated human T cells of interleukin 4 but not interferon-gamma is associated with elevated levels of serum antibodies to activating malaria antigens. Proc Natl Acad Sci U S A 87:5484–5488. doi:10.1073/pnas.87.14.5484

Tshikuka JG, Scott ME, Gray-Donald K, Kalumba ON. 1996. Multiple infection with Plasmodium and helminths in communities of low and relatively high socio-economic status. Ann Trop Med Parasitol 90:277–293. doi:10.1080/00034983.1996.11813053

Turner JD, Jackson JA, Faulkner H, Behnke J, Else KJ, Kamgno J, Boussinesq M, Bradley JE. 2008. Intensity of Intestinal Infection with Multiple Worm Species Is Related to Regulatory Cytokine Output and Immune Hyporesponsiveness. J Infect Dis 197:1204–1212. doi:10.1086/586717

Vidal M, Aguilar R, Campo JJ, Dobaño C. 2018. Development of quantitative suspension array assays for six immunoglobulin isotypes and subclasses to multiple Plasmodium falciparum antigens. J Immunol Methods 455:41–54. doi:10.1016/j.jim.2018.01.009

Wang Y, Jackson KJL, Chen Z, Gaëta BA, Siba PM, Pomat W, Walpole E, Rimmer J, Sewell WA, Collins AM. 2011. IgE sequences in individuals living in an area of endemic parasitism show little mutational evidence of antigen selection. Scand J Immunol 73:496–504. doi:10.1111/j.1365-3083.2011.02525.x

Wickham H, Averick M, Bryan J, Chang W, McGowan L, François R, Grolemund G, Hayes A, Henry L, Hester J, Kuhn M, Pedersen T, Miller E, Bache S, Müller K, Ooms J, Robinson D, Seidel D, Spinu V, Takahashi K, Vaughan D, Wilke C, Woo K, Yutani H. 2019. Welcome to the Tidyverse. J Open Source Softw 4:1686. doi:10.21105/joss.01686

World Health Organization. 2020a. World malaria report 2020: 20 years of global progress and challenges.

World Health Organization. 2020b. Soil-transmitted helminth infections. https://www.who.int/news-room/fact-sheets/detail/soil-transmitted-helminth-infections

World Health Organization. 2020c. Schistosomiasis. https://www.who.int/news-room/fact-sheets/detail/schistosomiasis

World Health Organization. 2021. Malaria. https://www.who.int/news-room/fact-sheets/detail/malaria

## References

Adegnika AA, de Vries SG, Zinsou FJ, Honkepehedji YJ, Dejon Agobé J-C, Vodonou KG, Bikangui R, Bouyoukou Hounkpatin A, Bache EB, Massinga Loembe M, van Leeuwen R, Molemans M, Kremsner PG, Yazdanbakhsh M, Hotez PJ, Bottazzi ME, Li G, Bethony JM, Diemert DJ, Grobusch MP. 2021. Safety and immunogenicity of co-administered hookworm vaccine candidates Na-GST-1 and Na-APR-1 in Gabonese adults: a randomised, controlled, double-blind, phase 1 dose-escalation trial. Lancet Infect Dis 21:275–285. doi:10.1016/S1473-3099(20)30288-7

Aguilar R, Ubillos I, Vidal M, Balanza N, Crespo N, Jiménez A, Nhabomba A, Jairoce C, Dosoo D, Gyan B, Ayestaran A, Sanz H, Campo JJ, Gómez-Pérez GP, Izquierdo L, Dobaño C. 2018. Antibody responses to α-Gal in African children vary with age and site and are associated with malaria protection. Sci Rep 8:9999. doi:10.1038/s41598-018-28325-w

Angov E, Aufiero BM, Turgeon AM, Van Handenhove M, Ockenhouse CF, Kester KE, Walsh DS, McBride JS, Dubois M-C, Cohen J, Haynes JD, Eckels KH, Heppner DG, Ballou WR, Diggs CL, Lyon JA. 2003. Development and pre-clinical analysis of a Plasmodium falciparum Merozoite Surface Protein-1(42) malaria vaccine. Mol Biochem Parasitol 128:195–204. doi:10.1016/s0166-6851(03)00077-x

Asojo OA, Goud GN, Zhan B, Ordonez K, Sedlacek M, Homma K, Deumic V, Gupta R, Brelsford J, Price MK, Ngamelue MN, Hotez PJ. 2010. Crystallization and preliminary X-ray analysis of Na-SAA-2 from the human hookworm parasite Necator americanus. *Acta Crystallogr Sect F*, Struct Biol Cryst Commun 66:172–176. doi:10.1107/S1744309109051616

Bergmann-Leitner ES, Hosie H, Trichilo J, Deriso E, Ranallo RT, Alefantis T, Savranskaya T, Grewal P, Ockenhouse CF, Venkatesan MM, Delvecchio VG, Angov E. 2013. Self-adjuvanting bacterial vectors expressing pre-erythrocytic antigens induce sterile protection against malaria. Front Immunol 4:176. doi:10.3389/fimmu.2013.00176

Black CG, Barnwell JW, Huber CS, Galinski MR, Coppel RL. 2002. The Plasmodium vivax homologues of merozoite surface proteins 4 and 5 from Plasmodium falciparum are expressed at different locations in the merozoite. Mol Biochem Parasitol 120:215–224. doi:10.1016/s0166-6851(01)00458-3

Briggs N, Wei J, Versteeg L, Zhan B, Keegan B, Damania A, Pollet J, Hayes KS, Beaumier C, Seid CA, Leong J, Grencis RK, Bottazzi ME, Sastry KJ, Hotez PJ. 2018. Trichuris muris whey acidic protein induces type 2 protective immunity against whipworm. PLoS Pathog 14:e1007273. doi:10.1371/journal.ppat.1007273

Cavanagh DR, McBride JS. 1997. Antigenicity of recombinant proteins derived from Plasmodium falciparum merozoite surface protein 1. Mol Biochem Parasitol 85:197–211. doi:10.1016/s0166-6851(96)02826-5

Doolan DL, Hedstrom RC, Rogers WO, Charoenvit Y, Rogers M, de la Vega P, Hoffman SL. 1996. Identification and characterization of the protective hepatocyte erythrocyte protein 17 kDa gene of Plasmodium yoelii, homolog of Plasmodium falciparum exported protein 1. J Biol Chem 271:17861–17868. doi:10.1074/jbc.271.30.17861

Gaur D, Mayer DCG, Miller LH. 2004. Parasite ligand-host receptor interactions during invasion of erythrocytes by Plasmodium merozoites. Int J Parasitol 34:1413–1429. doi:10.1016/j.ijpara.2004.10.010

Guerin-Marchand C, Druilhe P, Galey B, Londono A, Patarapotikul J, Beaudoin RL, Dubeaux C, Tartar A, Mercereau-Puijalon O, Langsley G. 1987. A liver-stage-specific antigen of Plasmodium falciparum characterized by gene cloning. Nature 329:164–167. doi:10.1038/329164a0

Imam M, Singh S, Kaushik NK, Chauhan VS. 2014. Plasmodium falciparum merozoite surface protein 3: oligomerization, self-assembly, and heme complex formation. J Biol Chem 289:3856–3868. doi:10.1074/jbc.M113.520239

Khusmith S, Charoenvit Y, Kumar S, Sedegah M, Beaudoin RL, Hoffman SL. 1991. Protection against malaria by vaccination with sporozoite surface protein 2 plus CS protein. Science 252:715–718. doi:10.1126/science.1827210

Kocken CHM, Withers-Martinez C, Dubbeld MA, van der Wel A, Hackett F, Valderrama A, Blackman MJ, Thomas AW. 2002. High-level expression of the malaria blood-stage vaccine candidate Plasmodium falciparum apical membrane antigen 1 and induction of antibodies that inhibit erythrocyte invasion. Infect Immun 70:4471–4476. doi:10.1128/iai.70.8.4471-4476.2002

Kusi KA, Bosomprah S, Dodoo D, Kyei-Baafour E, Dickson EK, Mensah D, Angov E, Dutta S, Sedegah M, Koram KA. 2014. Anti-sporozoite antibodies as alternative markers for malaria transmission intensity estimation. Malar J 13:103. doi:10.1186/1475-2875-13-103

Liu Z, Kelleher A, Tabb S, Wei J, Pollet J, Hotez PJ, Bottazzi ME, Zhan B, Asojo OA. 2017. Identification, Characterization, and Structure of Tm16 from Trichuris muris. J Parasitol Res 2017:4342789. doi:10.1155/2017/4342789

Metzger WG, Okenu DMN, Cavanagh DR, Robinson J V, Bojang KA, Weiss HA, McBride JS, Greenwood BM, Conway DJ. 2003. Serum IgG3 to the Plasmodium falciparum merozoite surface protein 2 is strongly associated with a reduced prospective risk of malaria. Parasite Immunol 25:307–312. doi:10.1046/j.1365-3024.2003.00636.x

Petzke MM, Suri PK, Bungiro R, Goldberg M, Taylor SF, Ranji S, Taylor H, McCray JW, Knopf PM. 2000. Schistosoma mansoni gene GP22 encodes the tegumental antigen sm25: (1) antibodies to a predicted B-cell epitope of Sm25 cross-react with other candidate vaccine worm antigens; (2) characterization of a recombinant product containing tandem-repeats of thi. Parasite Immunol 22:381–395. doi:10.1046/j.1365-3024.2000.00316.x

Rascoe LN, Price C, Shin SH, McAuliffe I, Priest JW, Handali S. 2015. Development of Ss-NIE-1 recombinant antigen based assays for immunodiagnosis of strongyloidiasis. PLoS Negl Trop Dis 9:e0003694. doi:10.1371/journal.pntd.0003694

Reddy KS, Pandey AK, Singh H, Sahar T, Emmanuel A, Chitnis CE, Chauhan VS, Gaur D. 2014. Bacterially expressed full-length recombinant Plasmodium falciparum RH5 protein binds erythrocytes and elicits potent strain-transcending parasite-neutralizing antibodies. Infect Immun 82:152–164. doi:10.1128/IAI.00970-13

Robson KJ, Hall JR, Jennings MW, Harris TJ, Marsh K, Newbold CI, Tate VE, Weatherall DJ. 1988. A highly conserved amino-acid sequence in thrombospondin, properdin and in proteins from sporozoites and blood stages of a human malaria parasite. Nature 335:79–82. doi:10.1038/335079a0

Siddiqui FA, Dhawan S, Singh S, Singh B, Gupta P, Pandey A, Mohmmed A, Gaur D, Chitnis CE. 2013. A thrombospondin structural repeat containing rhoptry protein from Plasmodium falciparum mediates erythrocyte invasion. Cell Microbiol 15:1341–1356. doi:10.1111/cmi.12118

Silva-Moraes V, Shollenberger LM, Castro-Borges W, Rabello ALT, Harn DA, Medeiros LCS, Jeremias W de J, Siqueira LMV, Pereira CSS, Pedrosa MLC, Almeida NBF, Almeida A, Lambertucci JR, Carneiro NF de F, Coelho PMZ, Grenfell RFQ. 2019. Serological proteomic screening and evaluation of a recombinant egg antigen for the diagnosis of low-intensity Schistosoma mansoni infections in endemic area in Brazil. PLoS Negl Trop Dis 13:e0006974. doi:10.1371/journal.pntd.0006974

Sim BK, Chitnis CE, Wasniowska K, Hadley TJ, Miller LH. 1994. Receptor and ligand domains for invasion of erythrocytes by Plasmodium falciparum. Science 264:1941–1944. doi:10.1126/science.8009226

Taechalertpaisarn T, Crosnier C, Bartholdson SJ, Hodder AN, Thompson J, Bustamante LY, Wilson DW, Sanders PR, Wright GJ, Rayner JC, Cowman AF, Gilson PR, Crabb BS. 2012. Biochemical and functional analysis of two Plasmodium falciparum blood-stage 6-cys proteins: P12 and P41. PLoS One 7:e41937. doi:10.1371/journal.pone.0041937

Versteeg L, Wei J, Liu Z, Keegan B, Fujiwara RT, Jones KM, Asojo O, Strych U, Bottazzi ME, Hotez PJ, Zhan B. 2020. Protective immunity elicited by the nematode-conserved As37 recombinant protein against Ascaris suum infection. PLoS Negl Trop Dis 14:e0008057. doi:10.1371/journal.pntd.0008057

Wei J, Damania A, Gao X, Liu Z, Mejia R, Mitreva M, Strych U, Bottazzi ME, Hotez PJ, Zhan B. 2016. The hookworm Ancylostoma ceylanicum intestinal transcriptome provides a platform for selecting drug and vaccine candidates. Parasit Vectors 9:518. doi:10.1186/s13071-016-1795-8

Wei J, Versteeg L, Liu Z, Keegan B, Gazzinelli-Guimarães AC, Fujiwara RT, Briggs N, Jones KM, Strych U, Beaumier CM, Bottazzi ME, Hotez PJ, Zhan B. 2017. Yeast-expressed recombinant As16 protects mice against Ascaris suum infection through induction of a Th2-skewed immune response. PLoS Negl Trop Dis 11:e0005769. doi:10.1371/journal.pntd.0005769

Yilmaz B, Portugal S, Tran TM, Gozzelino R, Ramos S, Gomes J, Regalado A, Cowan PJ, d’Apice AJF, Chong AS, Doumbo OK, Traore B, Crompton PD, Silveira H, Soares MP. 2014. Gut microbiota elicits a protective immune response against malaria transmission. Cell 159:1277–1289. doi:10.1016/j.cell.2014.10.053

Zhu J, Hollingdale MR. 1991. Structure of Plasmodium falciparum liver stage antigen-1. Mol Biochem Parasitol 48:223–226. doi:10.1016/0166-6851(91)90117-o

